# A spatial long-read approach at near-single-cell resolution reveals developmental regulation of splicing and polyadenylation sites in distinct cortical layers and cell types

**DOI:** 10.1101/2025.06.10.658877

**Authors:** Careen Foord, Andrey D Prjibelski, Wen Hu, Lieke Michielsen, Andrea Vandelli, Oleksandr Narykov, Brian Evans, Justine Hsu, Natan Belchikov, Julien Jarroux, Yi He, M Elizabeth Ross, Iman Hajirasouliha, Gian Gaetano Tartaglia, Dmitry Korkin, Alexandru I Tomescu, Hagen U Tilgner

**Affiliations:** Feil Family Brain and Mind Research Institute, Weill Cornell Medicine, New York, NY, USA; Center for Neurogenetics, Weill Cornell Medicine, New York, NY, USA; Department of Computer Science, University of Helsinki, Helsinki, Finland; RNA Systems Biology Lab, Center for Human Technologies, Istituto Italiano di Tecnologia; Bioinformatics and Computational Biology Program, Worcester Polytechnic Institute, Worcester, MA, USA; Computer Science Department, Worcester Polytechnic Institute, Worcester, MA, USA; Data Science Program, Worcester Polytechnic Institute, Worcester, MA, USA; Institute for Computational Biomedicine, Department of Physiology and Biophysics, Weill Cornell Medicine of Cornell University, New York, NY, USA; Caryl and Israel Englander Institute for Precision Medicine, The Meyer Cancer Center, Weill Cornell Medicine, New York, NY, USA; Physiology, Biophysics & Systems Biology Program, Weill Cornell Medicine, New York, NY, USA

## Abstract

Genome-wide single-cell and spatial long-read approaches have gained traction, but mostly lack single-cell resolution - and yield limited read lengths. Here, we introduce spatial ISOform sequencing (Spl-ISO-Seq), which reveals exons and polyadenylation sites from long reads with near-single-cell resolution. Spl-ISO-Seq selects long cDNAs and doubles to triples read lengths compared to standard preparations. Adding a highly specific software tool (Spl-ISOquant) and comparing human post-mortem pre-puberty samples of the visual cortex (8-11 years) to post-puberty samples (16-19 years), we find that cortical layers harbor stronger splicing and poly(A)-site regulation than the adjacent white matter, with enrichment of multiple protein-domain types. For oligodendrocytes however, developmental splicing changes are stronger in white matter. Among cortical layers, layer 4 has the most developmental changes in alternative-exon inclusion in excitatory neurons and in poly(A) sites. We also find many repeat elements, especially ERV1 long terminal repeats downstream of developmentally-regulated layer 4 exons. Overall, alternative splicing changes are linked to synapses – specifically at the post-synapse. Age-linked splicing changes in layers 1-3 and 4 are associated with autism spectrum disorder but not with schizophrenia, amyotrophic lateral sclerosis and Alzheimer’s disease. These results root developmental splicing changes during puberty and the resulting protein changes in specific layers and cell types. More generally, our new technologies enable new observations for any complex tissue.

## INTRODUCTION

Alternative splicing (AS) affects nearly all human protein-coding genes^1–4^, impacting neuronal properties, growth, excitability, synapse specification, plasticity, and neuronal network function^5–8^. Moreover, AS is associated with aging and developmental brain diseases^6,7,9–14^. Likewise, polyadenylation (poly(A))-site changes are abundant genome-wide and have been linked to cellular function^15–19^. The naming of long-read sequencing as the method of the year in January of 2023^20–25^ highlights this technology’s ability to record such RNA variables and their combinations. Using our technology of single-cell isoform sequencing (ScISOr-Seq)^26^, we recently profiled single-cell isoforms in cerebellum at postnatal day 1 (P1)^26^ as well as in pre-frontal cortex (PFC) and hippocampus at P7^27^. Additionally, multiple groups have tied alternative splicing to neurogenesis and differentiation^28,29^ and splicing differences between neuronal subtypes have been linked to the distinct expression patters of the splicing factors *Nova*, *Rbfox*, *Mbnl*, and *Ptbp*^30^. Additionally, for the Brain Initiative, with an enhanced version of ScISOr-Seq (ScISOr-Seq2), we defined brain-region specific full-length isoforms at single-cell resolution, covering cerebellum, hippocampus, thalamus, striatum and visual cortex at mouse postnatal day 56 (P56) and additionally mouse visual cortex and hippocampus at P14, P21 and P28^31^. Collectively, these data have revealed widespread cell-type specific isoform expression and brain-region specificity in isoforms for matched cell types – and dramatic developmental isoform changes in the mouse visual cortex for matched cell types. While certain markers in the cortex correlate with layer-specific location of cells, it is currently rarely possible to conclusively determine the layer in which an individual cortical neuron resides based solely on marker expression – especially in the human brain^32–34^. Therefore, in order to answer the question of which cortical layers are most strongly altered by splicing and poly(A)-site choice, technologies with spatial resolution are required. To this end, we and others have previously developed spatial isoform sequencing^27,35^ which revealed spatially regulated isoform expression events. Many of these occurred at transitions from one brain region to another, but others, like *Snap25*, occurred in a gradient throughout the mouse brain. At the employed spatial resolution of 60um, however, most physical spots covered multiple cells, thus hindering a cell-type specific view of spatially regulated isoforms. Additionally, our single-cell isoform investigation in human PFC^36^ and hippocampus^31^ have revealed many human alternative splicing events that cannot be modeled in mouse brain. Here, we address these limitations by developing spatial isoform sequencing (Spl-ISO-Seq) that improved the previous 60um to now 10um resolution. We coupled the spatial Curio Biosciences technology, a slide-based whole-transcriptome sequencing platform with high spatial resolution stemming from Slide-SeqV2^37^, along with a 2-step protocol enriching for exon-containing and long cDNAs. We add a specialized software package for the analysis of spatially barcoded long reads (Spl-ISOquant). Using fresh frozen human visual cortex (Brodmann area 17) samples from pre-puberty donors aged 8-11 years (“children”) to post-puberty (16-19 years, “young adults”; Y.A.), we investigate RNA biology regulation during this developmental period. Collectively, these approaches reveal widespread regulation of alternative exon inclusion, alternative acceptor usage, and poly(A)-site usage. Overall developmental RNA regulation in the cortex is stronger than in the white matter and among the cortical layers, layer 4 shows the highest regulation – an observation traceable to excitatory neurons. For oligodendrocytes, however, regulation is stronger in white matter than in cortical layers. Overall, we present a cell-type specific spatial transcriptomic technology optimized for long-read sequencing and identify region and cell-type specific splicing trends across human development.

## RESULTS

### A spatial view of pre-puberty and post-puberty samples of the visual cortex

Our new protocol for spatial isoform sequencing (“Spl-ISO-Seq”) at near-single-cell resolution ensures that polyadenylated RNA molecules are spatially barcoded based on tissue placement on a barcoded slide. The workflow proceeds by producing spatially barcoded cDNA, which is split into two pools. (i) The first pool was fragmented for generating Illumina short reads for gene-expression estimation and cell-type deconvolution and (ii) the second pool for enrichment of exon-containing molecules, long-molecule selection, and long-read isoform sequencing with Oxford Nanopore Technology (ONT) (**Figure 1a**). We used this approach to compare a group of four human visual cortex samples from children (8-11yrs) and four post-puberty young adults (Y.A.; 16-19yrs). We sequenced ∼245 million Illumina reads per individual. On average, these reads recovered 86.2% of expected barcodes (**Supplemental Figure S1**). The visual system and specifically Brodmann area 17 (V1) are well studied in their architecture and function^38–40^. V1 is composed of 6 layers which aid in visual signaling and processing. White matter is mainly composed of myelinated axons and oligodendrocytes. The layers of the visual cortex are distinguishable by key attributes including cell density, layer thickness, and layer-specific markers. Illumina UMI counts per spot suggested the position of distinct cortical layers (**Figure 1b**), which was confirmed with H&E stains of an adjacent tissue slice (**Figure 1c, Supplemental Figure S2**), short-read deconvolution methods^41^ (Methods), and known layer-specific markers. These layer-specific markers were used to help identify layer cutoffs and were identified with Illumina sequencing, since Illumina was less sparse compared to ONT sequencing (**Supplemental Figure S3**). We also employed a cell-type deconvolution program^41^ which identified which spots entirely contained individual cell types. Most cell-type deconvolution programs require large numbers of reads to accurately deconvolve cell types, thus Illumina sequencing was required for this process as ONT isoform sequencing lacked sufficient depth (**Supplemental Figure S4**) and could not sufficiently identify layer patterns or cell types (**Supplemental Figure S5**). Among spots classified as representing a single cell type (“singlets”), excitatory neurons were most abundant across all regions, followed by oligodendrocytes and vascular/endothelial cells (VENCs) (**Figure 1d,e, Supplemental Figure S6**). Out of all Illumina-identified spatial barcodes (n=482,777), excitatory neurons had the highest rate of assignment (**Supplemental Figure S7a**). Spots which were identified to contain two cell types (“doublets”) were less common compared to assigned singlets and empty spots (“rejects”), and the majority consisted of excitatory neurons paired with another cell type (**Supplemental Figure S7b-c**). Of note, 87.9% of all Illumina-identified spatial barcodes were also detected in ONT data. The barcodes which were highly sequenced by Illumina but not ONT were significantly upregulated in mitochondrial genes which are not targeted by the exome enrichment **(Supplemental Figure S8).** Comparing short-read gene expression between children and young adults, in agreement with the literature, we found increased expression of genes linked to long-term potentiation in the visual cortex in younger samples^42–44^. Conversely, and again agreeing with the literature^45,46^, we found genes linked to oxidative phosphorylation to increase during puberty (**Figure 1f**). In neurons, KEGG analysis revealed synaptic-vesicle-cycle genes and glutamatergic-synapse genes were upregulated. Likewise, glutamatergic-synapse genes as well as aldosterone-regulated sodium reabsorption were upregulated in astrocytes. As expected, oligodendrocytes also showed enrichment for myelin related genes with a gene ontology analysis (**Figure 1g**). Clustering analysis of short-read gene expression in excitatory neurons revealed that the strongest split represented the difference between white matter and cortex, with the second strongest split separating age groups rather than cortical layers. This observation supports the idea of robust cortical gene regulation during puberty (**Figure 1h**). Performing principal component analysis, principal component 1 (PC1) and PC2 of singlets also demonstrate variability deriving from mitochondrial gene expression and cell-type differences predominantly between excitatory neurons and oligodendrocytes, which likely represent overall differences between cortex and white matter (**Supplemental Figure S9**).

**Figure 1.**
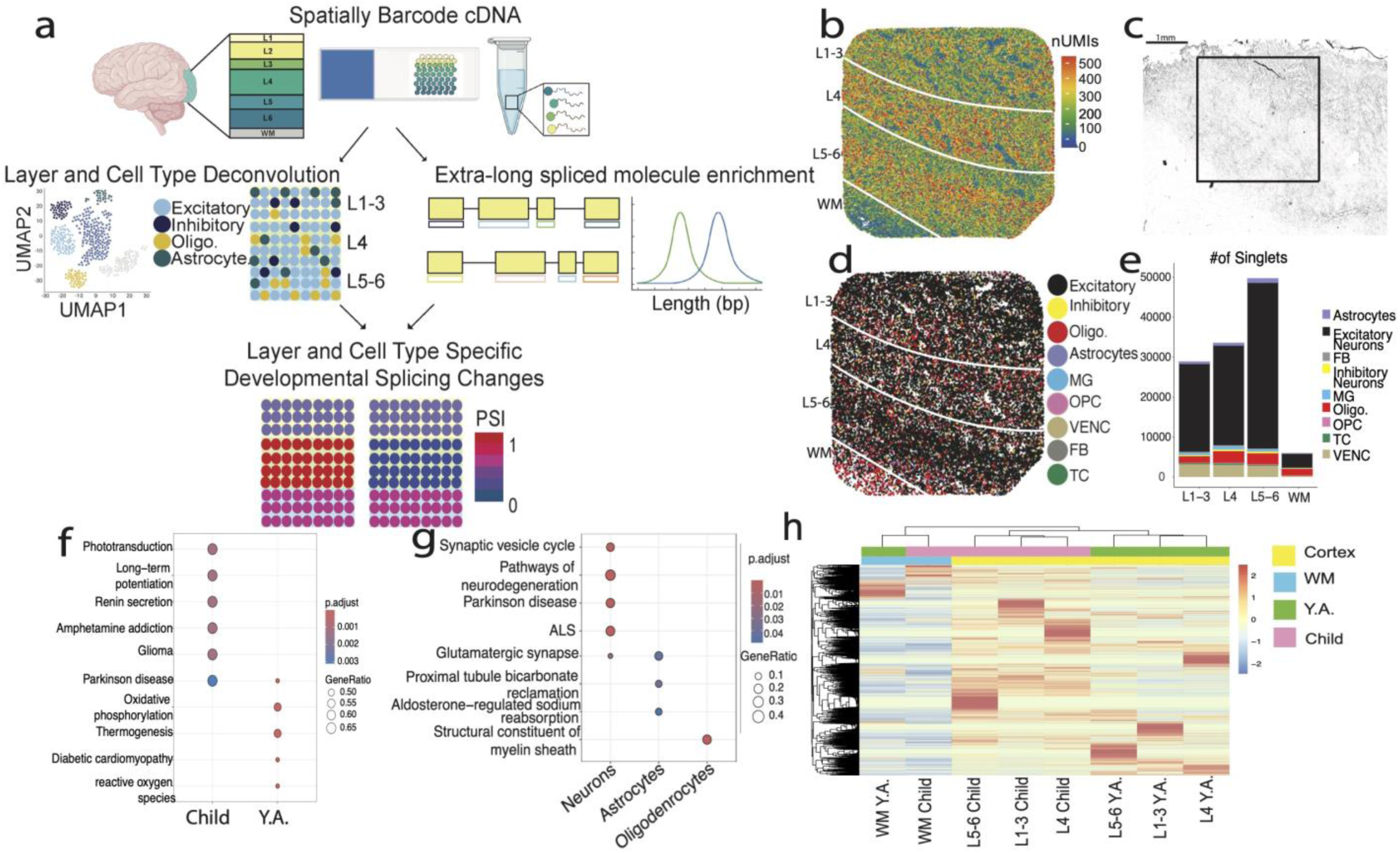
**a)** Experimental overview. Spatially barcoded cDNA is separated into 2 pools: one which is short-read sequenced and used for layer and cell-type deconvolution, and the other which undergoes exome enrichment, long-molecule selection, and is long-read sequenced. The two sets of data are then combined to examine layer and cell-type specific developmental splicing changes. **b)** UMI count per spot plotted by spatial location on sample 1115. **c)** Hematoxylin and Eosin stain on 10 uM thick slice of tissue following experimental section of sample 1115 with approximate area captured in black square. Scale bar indicates 1mm. **d)** RCTD defined singlets plotted by cell type and spatial location. Cell types include excitatory neurons, inhibitory neurons, oligodendrocytes (Oligo.), astrocytes, microglia (MG), oligodendrocyte precursor cells (OPC), vascular endothelial cells (VENC), fibroblasts (FB), and T cells (TC). **e)** Number of singlets across layers (L1-6) and white matter (WM) for all samples combined. **f)** KEGG enrichment of gene expression differences from short-read data when grouping by age groups, child (8-11) and young adult (16-19). **g)** KEGG enrichment of neuron and astrocyte singlets and GO enrichment for oligodendrocytes. h) Heatmap of short-read gene expression in excitatory

### A long cDNA enrichment approach enables improved spatial and single-cell isoform recovery

Long reads hold the promise to reveal the combination patterns of variable sites (for example TSS, all splice sites, poly(A)-sites on RNAs)^21^. However, cDNAs produced by spatial approaches or single-cell DNAs are commonly short, often due to truncation of full transcripts. In the case of Curio Bioscience’s library preparation, a random hexamer rather than a 5’ TSO molecule is employed to generate 2nd strands. Thus, most molecules are not complete as the library preparation is not designed to capture full length molecules, but rather only gene expression. We hence engineered cDNA amplification and purification methods to enrich for longer, spliced cDNAs. After isolating spatially barcoded cDNA following the Curio Bioscience’s pipeline, exon-spanning cDNA was enriched for using Agilent exome enrichment probes (Methods), which removes purely intronic molecules and enriches for exonic ones. Additionally, the PCR following exome enrichment was stopped at cycle 8 and cleaned up with 0.48x SPRIselect beads in 1.25M NaCl-20% PEG buffer to yield long cDNA. We then performed another PCR such that only a longer set of the cDNAs was amplified, followed by another cleanup (Methods). We optimized Spl-ISO-Seq using two pre-frontal cortex samples (one child and one young adult) (**Supplemental Figure S10**). Using the Curio spatial-genomics approach and naïve long-read sequencing, we see a median read length of 502 bp (“Standard”). Employing exome-enrichment probes to remove intronic cDNAs left this number largely unchanged (“Standard Exome”). Our new protocol, involving exome enrichment and long-cDNA enrichment (“Long Exome”), increases the median read length 2.7-fold to 1,358 bp (**Figure 2a**; Standard: Mean=775.61, Standard Deviation (SD) =941.65; Standard Exome: Mean=725.53, SD=455.23; Long Exome: Mean=1409.45, SD=673.90). The Long Exome dataset contained a slightly higher percentage of intron-retained reads compared to the other two protocols (**Supplemental Figure S11**). For spliced reads, however, the exome enrichment step yields a dramatic increase in spliced reads (**Figure 2b**). Taken together the long protocol increases the median exon number per spliced read from 4 to 5 (**Figure 2c**; Standard: Mean=4.43, SD=1.71; Standard Exome: Mean= 4.41, SD=1.67; Long Exome: Mean=5.84, SD=2.90). Furthermore, comparing the number of reads assigned to each gene between Standard and Long Exome experiments yielded a high correlation (**Figure 2d**; Pearson’s R=0.834, p<2.2e-16). Similarly, transcript expression is also correlated between the two datasets (**Supplemental Figure S12a**, Pearson’s R=0.69, p< 2.2e-16) and transcript expression is not related to transcript length (**Supplemental Figure S12b**). Thus, the enrichment for long fragments does not strongly shift gene expression patterns when comparing two exome enriched samples. Additionally, performing the exome enrichment itself does not significantly alter gene expression compared to the standard preparation (**Supplemental Figure S13**; Pearson’s R=0.814, p<2.2e-16). Not observing a strong shift in gene expression profile yet having longer reads could be explained by covering a larger fraction of the isoforms to which the molecules are assigned. We therefore calculated the fraction of bases of an annotated transcript covered by a read after the read was assigned to this transcript. Using the Long Exome approach, this fraction increased from 0.49 to 0.70 and from 0.53 to 0.59 in the two samples, respectively (**Supplemental Figure S14, Supplemental Figure S15a-b**). Of note, even in cDNAs of bulk full-length molecules^47^ the resulting fraction of transcript coverage is 0.89. The difference between sequencing methods becomes even more pronounced when only examining spliced reads (**Supplemental Figure S15c-d**). Overall, the Long Exome approach allows for the capture of larger portions of transcripts, which is greatly increased in longer transcripts compared to the Standard LR and Standard Exome LR (**Supplemental Figure S15e-h**). Importantly, a truncation simulation of ONT reads suggests that isoform-assignment recall and precision are affected by the coverage fraction per transcript (**Supplemental Table 1**). Thus, enrichment for longer molecules both tends to cover more of the original RNA isoforms that the reads represent, as well as maximizes the accuracy of isoform assignment and subsequent quantification. An example of this higher fraction of transcript bases covered can be found in the *RPS6KB2* gene. Before selection of long cDNAs, the sequenced reads cover an average of 5.4 exons per read, while with selection of long cDNAs, the sequenced reads cover an average of 12.3 exons per read and usually reach the first annotated exon^48^ (**Figure 2e**). We then explored whether short-read analysis alone could serve for splicing analysis in our brain slices. To this end, we counted the number of detected splicing events per barcode and added these from all barcodes, which we termed intron-barcode pairs. For the short-read experiment, we found an average of 108,343.6 events per sample while we found 5,910,532 for the Spl-ISO-Seq protocol – a 55-fold increase (**Figure 2f**; Naive Short Read (SR): SD =39672.25; Long Exome Long Read (LR): SD=3242296). Of note, spliced short reads showed dramatically lower gene coverage compared to both naïve short read and Long Exome LR sequencing (**Supplemental Figure S16**). Although Naïve Short read data showed high depth, gene coverage was limited to the regions near the Poly(A) site. In contrast, Long Exome LR data showed moderate, although more uniform coverage. We then tested whether the selection for long molecules had equally positive effects for our previously published single-cell experiments. Indeed, the long protocol has even more drastic increases in a separate, mouse single-cell sequencing experiment: the read length is increased from a median of 810 to 1,688 bp (both after exome enrichment, **Figure 2g**; Standard Exome: Mean=919.83, SD=463.88; Long Exome: Mean=1701.81, SD=592.45). While the spliced-read fraction increases only slightly (**Figure 2h**), the median exon number per read rises from 4 to 8 (**Figure 2i**; Standard Exome: Mean= 4.98, SD=2.46; Long Exome: Mean=8.19, SD=3.94). This demonstrates that this new protocol functions well on cDNA derived from multiple protocols and regions. Taken together, these findings show that both steps of our new protocol, namely exome enrichment and long-cDNA isolation are required for yielding spliced reads derived from transcripts of different sizes.

**Figure 2.**
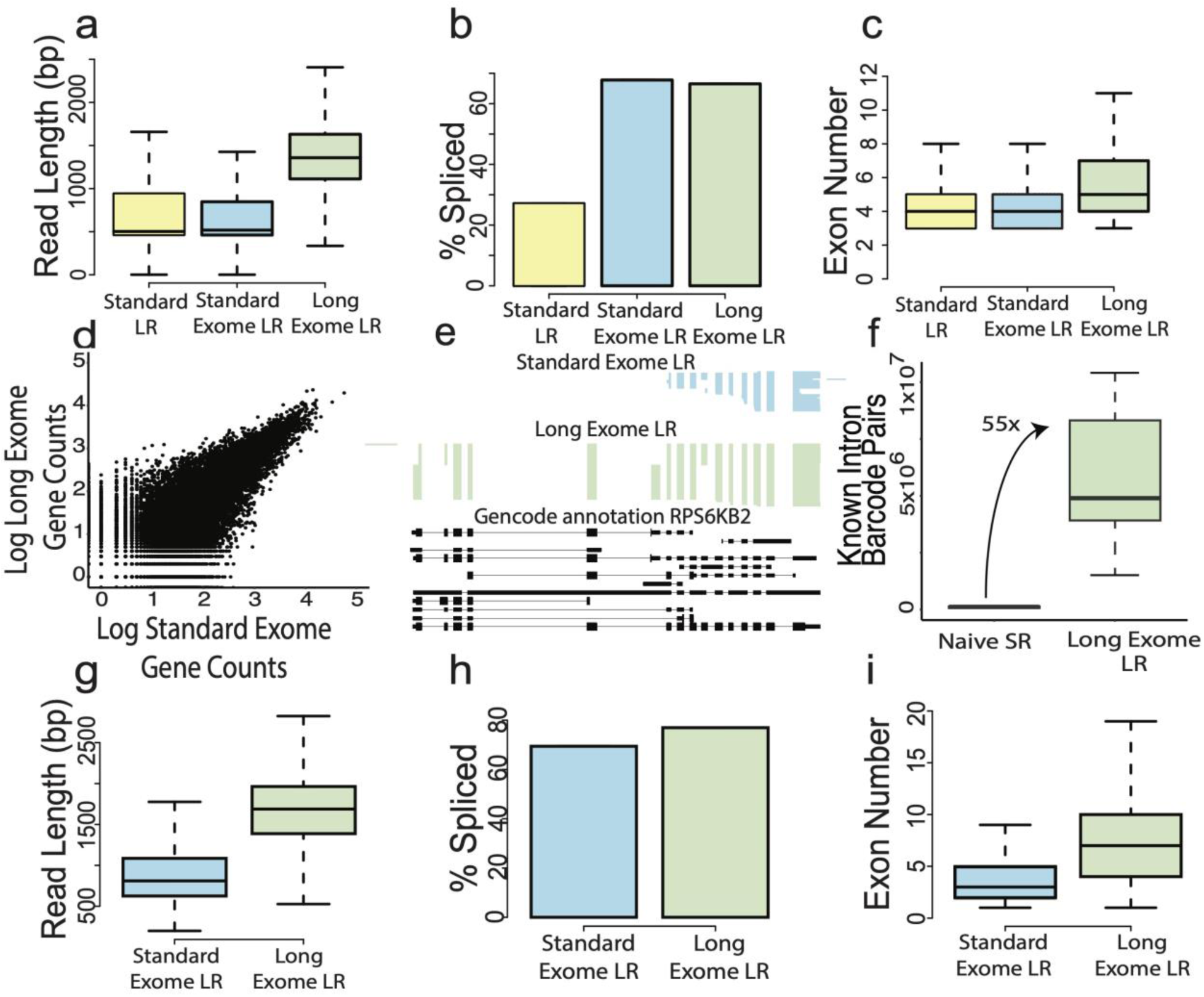
**a)** Read length of 100,000 long reads randomly sampled from each spatial dataset (n=2). Standard: cDNA from standard protocol prior to tagmentation; Standard Exome: cDNA from standard protocol which is enriched for exonic reads; Long Exome: cDNA from standard protocol which is enriched for exonic reads and longest molecules. **b)** Percent of spliced molecules from each group. **c)** Number of exons per read in barcoded, mapped, and spliced reads. **d)** Correlation of counts per gene of a Standard Exome dataset compared to the Long Exome dataset from the same spatial slide (n=1). Counts are log10 transformed. **e)** ScisorWiz^48^ plot of the gene RPS6KB2. Each horizontal line is a sequenced read from each respective dataset which come from the same slide. Gencode annotated transcripts are plotted in black. **f)** Total number of intron-barcode pairs from naive Illumina sequencing compared to Long Exome sequencing from all samples (n=8). **g)** Read length of 10,000 reads randomly sampled from each single-cell dataset (n=1). **h)** Percent of spliced molecules per group. **i)** Number of exons per read in barcoded, mapped, and spliced reads.

### Spl-ISO-quant enables long-read analysis including highly specific barcode deconvolution for spatial experiments with near-single-cell resolution

We have recently published IsoQuant and shown that it is highly specific in terms of isoform detection and read-to-isoform assignment for both Oxford Nanopore and PacBio long-read experiments^49^. IsoQuant has recently been shown - without our involvement - to exhibit very strong performance. Specifically, the authors found IsoQuant as “a highly effective tool for isoform detection with LRS, with Bambu and StringTie2 also exhibiting strong performance”^50^. Here, we added barcode deconvolution and UMI deduplication, as well as isoform quantification (**Figure 3a**). While these problems were previously addressed by various tools^51–54^ (BLAZE, SiCeLoRe, FLAMES, and scnanoseq), all of these methods are designed specifically for 10x Genomics long-read data and cannot be easily adapted for other protocols with a distinct molecule structure, such as Curio. Furthermore, Curio has about 10-fold more barcoded spots in a single experiment compared to a typical 10x Genomics single-cell experiment. Additionally, Curio features barcodes of 14 bp versus 16 bp long barcodes for 10x. Thus, the Curio data is characterized by substantially higher barcode collision probability (roughly 100-fold), which even further complicates the barcode demultiplexing process. Using the known barcode structure of the Curio protocol, we first detect PCR primer and barcode linker positions in the vicinity of poly(A)-tails. The known barcodes that come with each of the purchased slides are exploited to build a k-mer index, which allows to quickly obtain a limited number of candidate barcodes for every read. For these candidates a Smith-Waterman alignment is employed to assign a score to each barcode candidate (Methods). The maximal score of 14 represents 100%-identity of the long-read derived barcode candidate and a barcode in the prior list. This score is then used to determine the correct barcode, and the position of the barcode yields the candidate sequence of the UMI (**Figure 3b**; Methods). Subsequently, barcoded, spliced, and UMI filtered reads are used for downstream splicing analysis, where UMI filtering does not introduce any significant bias with respect to read length or splice sites (**Supplemental Figure S17, Supplemental Table 2**). We performed simulation experiments, introducing errors into in-silico reads according to an error profile learned from Oxford Nanopore alignments including read truncation, which is a commonly occurring phenomenon in ONT sequencing (Methods). We used these experiments to determine precision and recall of our barcode-calling strategy. Allowing barcode calls with a minimal score of 11, yielded a precision of 91.6%. This number rose to 97.3% (score=12), 99.5% (score=13) and 99.9% (score=14). As expected, recall also decreased with higher score requirements from 55.2% (score=11) down to 50.4% for score 13 (**Figure 3c**). Given the small precision difference between score=13 and score=14, we used score=13 in the following analyses. We also verified if specific barcodes with score=13 could be more problematic than others. Overall, close to all barcodes showed a precision of >=95%. However, a very few barcodes (118 of ∼70k) showed precision <85% and 14 barcodes precision of 0-5%. These few barcodes contribute a small fraction of reads and thus do not affect slide-wide conclusions. However, if any statements are made for a very small number of barcodes (<=10), our results indicate that such barcodes need to be tested for their individual precision to avoid false-positive results derived from few barcodes (**Figure 3d**). Of note, the above simulations included substitutions, insertions and deletions as well as read truncation. The former three error sources in principle still allow for reads covering the entirety of the barcode - a situation, in which errors can potentially be remedied by state-of-the-art algorithms. Read truncation however can completely delete a barcode - a situation that no algorithm could ever remedy. To understand the influence of read truncation, we performed simulation experiments without read truncation. This left precision largely unchanged, but increased recall to 83.3% (score=11), 81.8% (score=12), 75.4% (score=13) and 70.9% (score=14). Thus, a large portion of the recall loss is due to read truncation (**Figure 3e**). In the simulation scenario with truncation, the largest numbers of reads were lost when asking for the presence of the poly(A) tail and/or the linker in between the two barcodes. This loss did not occur in the simulation without truncation (**Figure 3f**). In summary, Spl-IsoQuant enables highly precise barcode recognition and pinpointing of problematic barcodes. The strongest contributor to not finding barcodes is read truncation - a chemical occurrence that cannot be remedied by algorithms.

**Figure 3.**
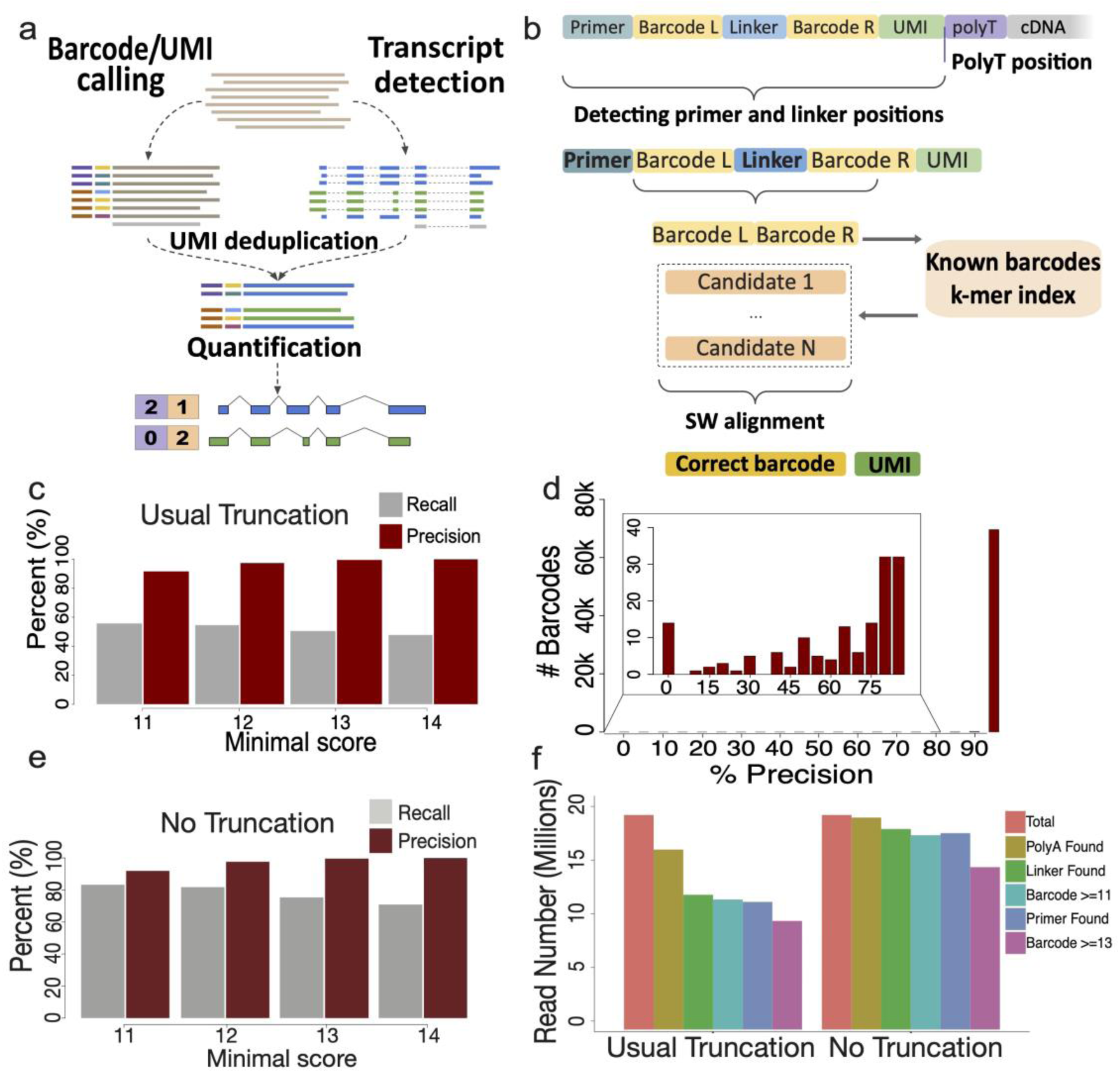
**a)** Outline of Spl-IsoQuant algorithm. **b**) Barcode calling and UMI determination in Spl-IsoQuant. **c**) Precision and recall as a function of minimal scores overall in Spl-IsoQuant with truncation which is representative of ONT data. **d**) Precision for individual barcodes. **e**) Precision and recall as a function of minimal scores overall in Spl-IsoQuant with no truncation. **f**) Number of reads found by Spl-IsoQuant to meet criteria cutoffs separated by reads modeled with usual truncation and no truncation.

### Developmental changes in splicing affect cortical layers more than white matter

The architecture and function of the visual system and V1 are well studied in the neuroscience field^38–40^. The visual cortical layers are distinguishable by key attributes including cell density, layer thickness, and layer-specific markers. We used H&E stains (**Supplemental Figure S2a-h**) and short-read deconvolution (**Supplemental Figure S18a-h**) to identify cell densities and layer thickness and examined the location of layer-specific marker expression. Based on these measurements, we divided each slide into layers 1-3 (“L1-3”), layer 4 (“L4”), layers 5-6 (“L5-6”) and white matter (“WM”). In addition to the previously mentioned two PFC samples, we long-read sequenced eight visual-cortex samples to yield four male children individuals (8-11yrs) and four male young adults (16-19 yrs) with ONT (**Supplemental Figure S19a-f; Supplemental Table 3**). Intron lengths were in-line with our previous results, in that close to no expressed introns were found below 67 bases. Internal-exon length (average=143 bases) and terminal-exon length (average=642 bases) were both in-line with recent literature^55^ (**Supplemental Figure S20a-b**). As we established previously^56,57^, extremely few introns are <70 bases and few introns extended past 20kb (**Supplemental Figure S20c-d).** Overall, barcoded molecules assigned to a transcript identified 19,686 annotated genes with >=10 UMIs. Downsampling experiments showed some signs of saturation although perfect saturation was not attained (**Supplemental Figure S20e**). Similar trends are found when repeating experiments with >=3 UMIs (**Supplemental Figure S20f**). We first tested individual exons for distinct exon inclusion between children and young adults (Methods). An alternative exon is included or excluded in a cDNA molecule and here we consider two age groups – leading to a 2×2 contingency table^3,27,36^. Of note, we found many exons with deltaPSI values close to -1 or +1 in the cortical layers, implying complete developmental shifts in exon inclusion (**Figure 4a**). In white matter, we observed much fewer of such extreme shifts (**Figure 4b**). In the cortical layers, 18.54% (95%-confidence interval [18.03,19.05]) of tested exons changed inclusion during puberty, while only 6.14% (95%-confidence interval [4.96,7.31]) did so in white matter (**Figure 4c, Supplemental Figure S21**). Of note, performing the same analyses with spliced short reads resulted in a 93.3-fold decrease in cortical tested exons, where not a single exon was identified to be regulated across age after FDR correction (**Supplemental Figure S22**). Considering deltaPSI values of significant exons in both compartments, we found those in the white matter to exhibit negative (that is decreasing inclusion with developmental time) changes more often than those in the cortical layers – the latter of which showed positive and negative changes at a more balanced ratio (**Figure 4d**). Importantly, larger spot numbers and read numbers per spot can lead to higher significance numbers in one of the areas. To test whether this affected cortex and white matter differentially, we performed downsampling experiments (Methods) that equalized data imbalances. These downsampling experiments confirmed that indeed cortical layers had a higher rate of developmentally-regulated exon inclusion than the white matter did (corrected 2-sided Wilcoxson rank sum test; FDR= 1.1×10^-11^, **Figure 4e**; Cortex: Mean=0.55, SD=0.80; WM: Mean=0.0, SD=0.0). We additionally performed simulations to examine how read number and |deltaPSI| contributed to false positive and negative exons detected (**Supplemental Figure S23a**). This analysis determined that 10-19 reads per comparison is necessary to detect a change in exon inclusion, which aligns with the results (**Supplemental Figure S23b**). An example of alternative-splicing regulation specific to the cortical layers is the *SLC12A2* gene, in which an exon increases inclusion from 19.3% to 70.2% in cortex across age. In white matter however, we found no significant change (**Figure 4f**). This exon keeps the reading frame and has been shown to impact sodium influx by the cell^58^. Searching for SynGO^59^ enrichment of genes with significant developmental exon inclusion changes in cortical layers yielded strong enrichment for “postsynaptic density”, “synaptic vesicle cycle” and “receptor localization” in location or function mode of SynGO (**Figure 4g,h**). However, similar analysis for genes with altered exons in the white matter gave no such enrichment (**Supplemental Figure S24**). Thus, developmentally regulated splicing in cortical layers is heavily connected to the postsynaptic density. Next, we identified exons which contain start codons and coding sequences based on published annotations. Cortical developmentally regulated exons (FDR < .05 & |deltaPSI| > 0.2) contained start codons less often than exons that were not developmentally regulated (FDR > .05 & |deltaPSI| < 0.05; corrected 2-sided Fisher test, FDR = 1.19×10^-8^). Thus, there is a tendency not to alter translation initiation in postnatal development of the cortical layers. In the white matter however, we did not observe such a difference between developmentally-regulated and non-regulated exons (corrected 2-sided Fisher test, FDR=0.39; **Figure 4i**). On the other hand, developmentally regulated exons in the cortical layers contained non-coding sequence significantly more often than non-regulated exons – an observation that we could again not make in the white matter (corrected 2-sided Fisher test, cortex FDR=2.40 x10^-22^; **Figure 4j**). We next considered exons which are developmentally regulated in the cortex. Exploiting the near-single-cell nature of this spatial technology and short-read gene expression, we classified each physical spot as either excitatory neuron, inhibitory neuron, oligodendrocyte, astrocyte, microglia, vascular cell, fibroblast or unclassifiable **(Supplemental Figure S18)**. Unclassifiable spots usually have low RNA content, either because of biological low expression, such as a cell-free area or because of technical limitations. Separating the most abundant cell type, excitatory neurons, from other spots, we found developmentally regulated exons in excitatory neurons to show a strong trend for decreasing their inclusion with increases in age, while exons from other spots showed more balanced proportions of increasing and decreasing proportions (2-sided Wilcoxon rank sum test, p=9.44 x10^-9^; Excitatory Neuron (ExN): Mean=-0.04, SD=0.43; Non-ExN: Mean=0.03, SD=0.41; **Figure 4k**). Considering the second most abundant cell type – oligodendrocytes – we asked whether WM oligodendrocytes had splicing changes more frequently than cortical oligodendrocytes. In opposition to the trends seen in all cortical and white matter spots, we found WM oligodendrocytes to show splicing changes more frequently than cortical oligodendrocytes (**Figure 4l**; Cortex Standard Error (S.E.) [0,0]; WM S.E. [15.60,24.40]). Sufficient data was not available to down-sample this result. However, WM overall has a smaller number of assigned oligodendrocytes than in the combined cortical layers, suggesting that this result is not due to power imbalances. We then searched for protein domains that are affected by cortical developmentally regulated exons and considered the percentage of exons affected by age. Exons were grouped based on which age group they were most included (higher in children or higher in Y.A.). As a background set, we considered exons without any apparent regulation (Methods). After a Fisher test of higher-young-adult inclusion and higher-children inclusion exons and a Bonferroni correction, 14 protein domains showed differences. Some of these groups affected few exons overall, but others had higher differences. For example, voltage-gated potassium channels were present in 13.6% of higher-adult-inclusion exons but only in 0.64% of higher-children-inclusion exons. Thus, voltage-gated potassium channels are increasingly used in early adulthood (as compared to younger children) and a similar observation was made for bromodomains. Conversely, Armadillo repeats were frequently observed in higher-children-inclusion exons (**Figure 4m**). Armadillo repeat (ARM) domains is an example of a ubiquitous structural fold of alpha solenoid consisting of a varying number of stacked short helices^60^. *UBR4* is a gene that contains two ARM domains and has been identified as one of the key proteins in the mammalian brain associated with neurogenesis, neuronal migration, and neuronal signaling^61^. However, the role of AS in regulating its functional activity has not yet been studied. We found that *UBR4-204* has both ARM domains, where others either have both domains missing (*UBR4-203*), or a shortened copy of one ARM domain (*UBR4-202*), which could indicate the absence or substantial reduction of its function (**Figure 4n**). Interestingly, *UBR4-204* reads are increased in the child cortex compared to young adults. That trend is flipped in the *UBR2-202* where reads are increased in young adults rather than children (**Figure 4o**). Due to the modular nature of the armadillo repeat domains, consisting of short helices of the same length, it is expected that the shorter version of the ARM domain will fold into a stable structure even though shorter proteins are generally more unstable (Methods, **Figure 4p**). The observed stability can be explained by the previously identified property of ARM repeats to self-assemble even when they are non-covalently bound^62^. Furthermore, the modular stability of ARM, TRP, and other repeat proteins has increasingly been utilized in the modular design of synthetic repeat proteins^63^.

**Figure 4.**
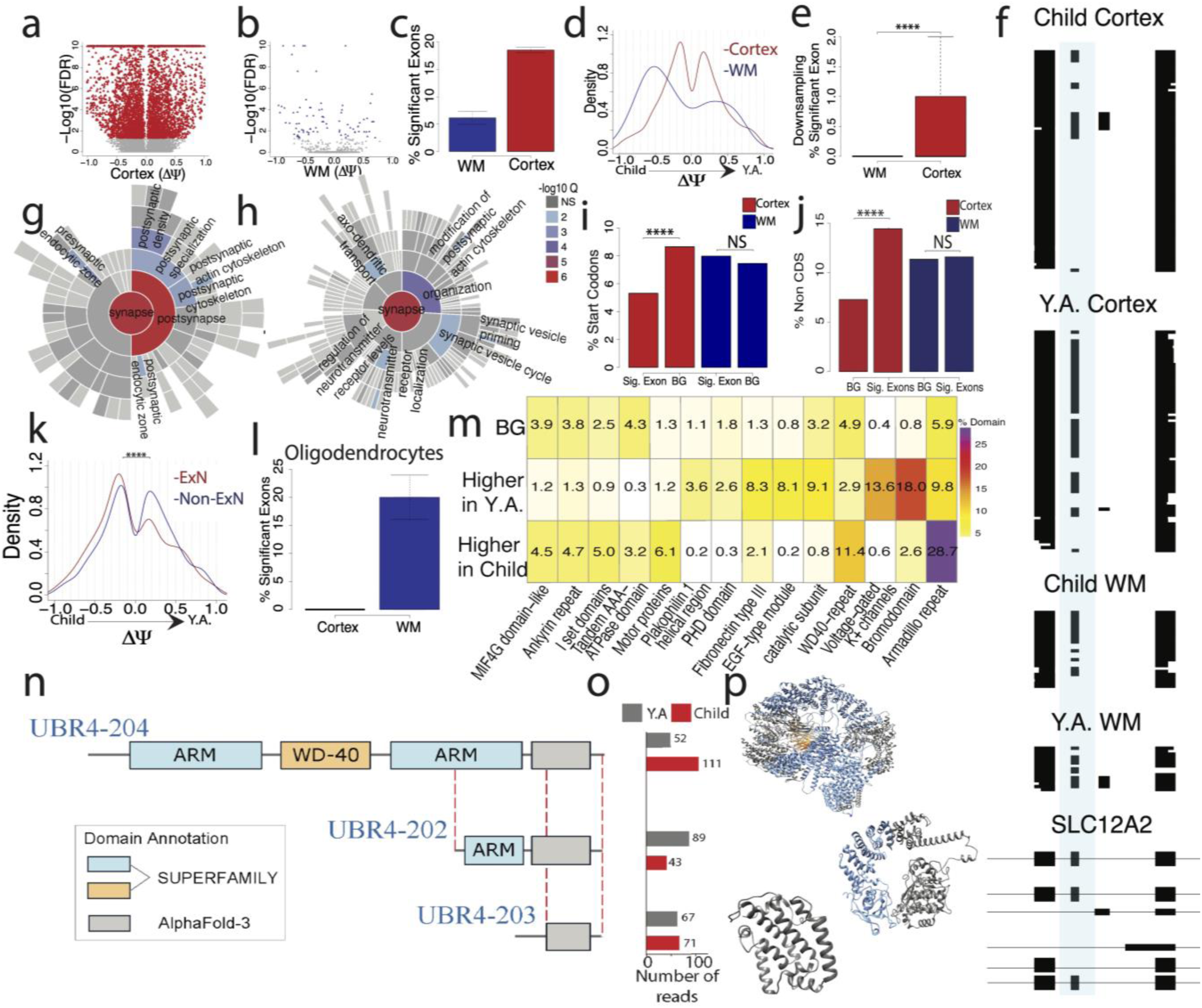
**a)** Exons tested between ages in cortex by deltaPSI (YA-Child) and -log10(FDR). Red points are significant exons and grey points are non-significant. **b)** Exons tested between ages in WM by deltaPSI and -log10(FDR). Blue points are significant exons and grey points are non-significant**. c)** Percent significant of tested exons and |deltaPSI| >=0.2 for cortex and WM. **d)** deltaPSI density of significant exons for cortex and WM. **e)** Percent significant exons from downsampling experiments. Exon selection and percent significant calculations are repeated 100x and plotted. **f)** ScisorWiz^48^ plot of exon chr5_128177105_128177152_+ in *SLC12A2* gene. **g)** SynGO location enrichment of significant cortex genes. **h)** SynGO function enrichment of significant cortex genes. **i)** Percent of exons associated with start codons between highly significant and a background set of exons. (**j)** Percent of non-CDS exons between highly significant and a background set of exons. **k)** deltaPSI density of significant exons of excitatory neurons and non-excitatory neurons. **l)** Percent significant of tested exons and |deltaPSI| >=0.2 for cortex and WM in Oligodendrocytes. **m)** Enrichment of protein domains in cortex highly significant alternative exons (|deltaPSI| > .5 and FDR <.05). **n)** Diagram of 3 transcripts of the multidomain protein UBR4 which affect its domain architecture. Transcripts contain 2 Armadillo repeats (ARM) and 1 WD-40 domain (UBR4-204), 1 shorter ARM repeat only (UBR4-202), or neither domain types (UBR4-203). **o)** Number of spliced and barcoded reads assigned to each transcript separated by Y.A. and Child. **p)** AlphaFold3 predictions of protein structure; blue = ARM repeats and yellow = WD-40.

### Developmental polyadenylation regulation equally affects cortical layers more than the white matter

Our assay equally reveals changes in poly(A) sites. We tested whether poly(A) sites would follow similar trends as alternative exons using our concept of poly(A)-site tests^27,31^. In contrast to alternative exons with two possible states, a gene can have many poly(A) sites. We therefore employ 2xN tests for poly(A)-site testing^27^. Similarly, as alternative exons, we found a higher fraction of genes (estimate=24.67; 95%-confidence interval [23.52,25.81])) in cortical layers as compared to white matter spots (estimate=3.70; 95% confidence interval [2.55,4.85]). However, while we had observed a three-fold higher value for alternative exons between cortical layers and white matter, for poly(A)-site choice we observed a >6-fold higher level (**Figure 5a**). In contrast to deltaPSI values for alternative exons, the deltaPI values for poly(A) sites are, by definition, positive. For genes with significant poly(A)-site regulation these values were stronger for the white matter, which is likely a consequence of lower read numbers requiring stronger effect sizes to achieve significance (**Figure 5b**; Cortex: Mean=0.25, SD=0.20; WM: Mean=0.29, SD=0.17). Downsampling experiments again showed that the observation of more poly(A)-site regulation in the cortex than in the white matter remains true when equalized statistical power is enforced (2-sided Wilcoxon rank sum test; p=8.12×10^-38^; Cortex: Mean=7.92, SD=4.48; WM: Mean=0, SD=0; **Figure 5c**). An example of this is the *VBP1* gene, in which the usage of a downstream poly(A) site is dramatically increased with age in the cortex, while in white matter the downstream site is exclusively used regardless of age (**Figure 5d**). Poly(A)-site choice defines the length of the transcript’s last exon. We separated genes into 2 groups based on those which lack developmentally regulated Poly(A)-sites (|deltaPI| <0.2, “non-changing”) and those which undergo developmentally regulated Poly(A)-sites (|deltaPI| <0.2, “changing”). We defined these groups based on |deltaPI| criteria rather than significance in order to limit the number of false positives while also capturing substantial changes which may lack sufficient power to reach significance. We first measured last-exon length in non-changing tested genes (|deltaPI| <0.2; Methods) both in children’s cortices young adults’ – and similarly in the white matter. No significant differences in length were detected between regions and age groups (**Figure 5e**; WM Child: Mean=443.34, SD=303.42; WM Y.A.: Mean=404.67, SD=278.49; Cortex Child: Mean=431.52, SD=291.08; Cortex Y.A.: Mean=444.89, SD=316.33). However, genes with poly(A)-site regulation (|deltaPI| > 0.2) in development had shorter last exons in the white matter than in cortex – regardless of age group (corrected 2-sided Wilcoxon rank sum tests; cortex Y.A. v WM Y.A. FDR=4.96×10^-5^; cortex Y.A. v WM child FDR=1.36×10^-3^; WM Y.A. v cortex child FDR=1.31×10^-5^; WM child v cortex FDR=4.52×10^-4^; WM Child: Mean=326.57, SD=198.45; WM Y.A.: Mean=315.09, SD=197.41; Cortex Child: Mean=426.70; SD=278.65; Cortex Y.A.: Mean=428.03, SD=294.29). Thus, white-matter poly(A)-site regulation affects genes with short last exons (**Figure 5f**). An example of this length observation can be seen in the *KAT8* gene (**Supplemental Figure S25**). Additionally, a similarly to the length of last exons, the UTR length of genes with regulated poly(A) sites is shorter in WM compared to cortex (**Figure 5g**). We also found that the genes which exhibit developmentally regulated exon inclusion (|deltaPSI| > 0.2) also tend to show developmentally regulated Poly(A) site usage in the cortex (2-sided Fisher test p=1.49×10^-16^; **Figure 5h**), indicating potential coordination between these processes.

**Figure 5.**
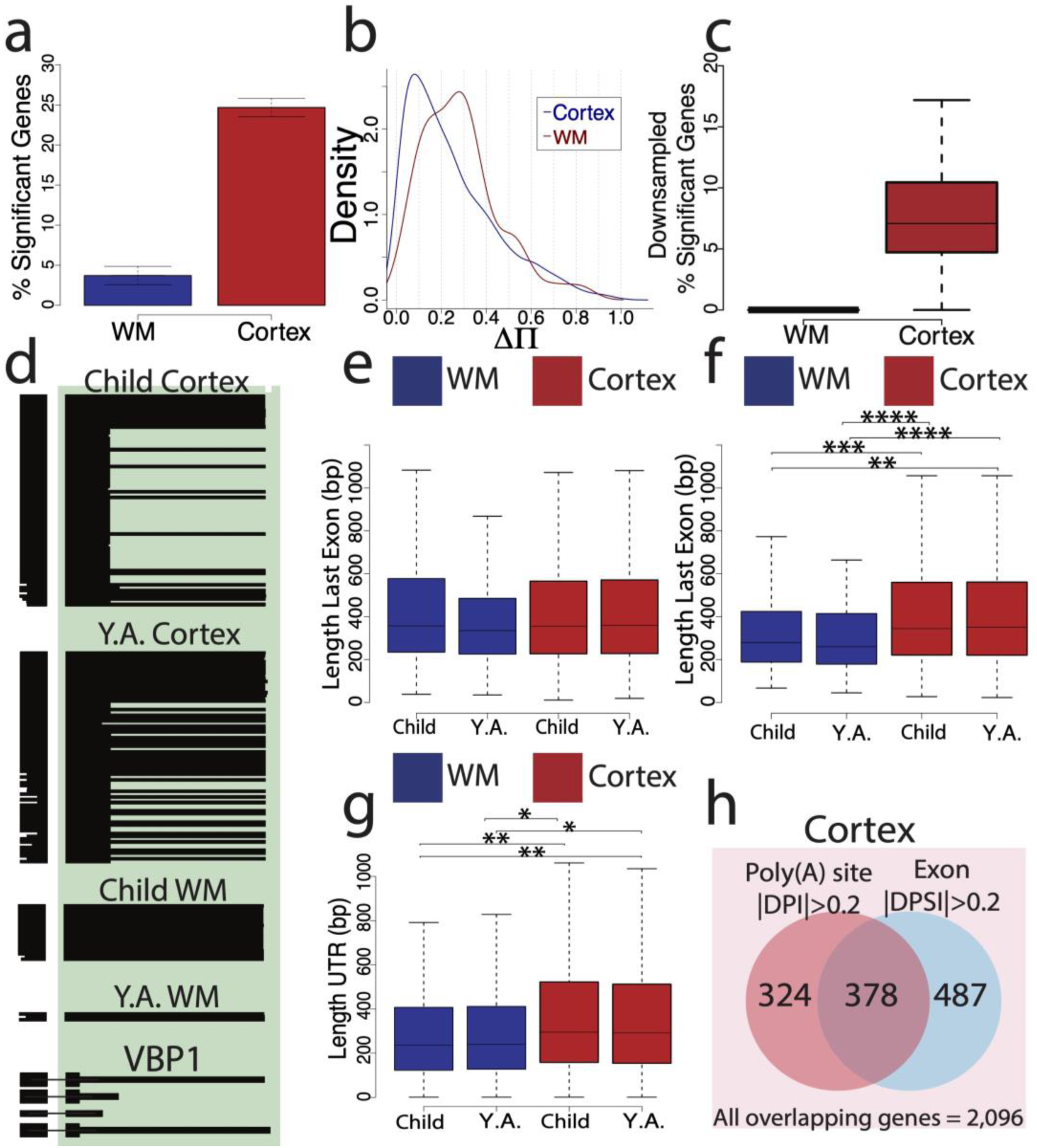
**a)** Percent significant genes with differential polyA sites. **b)** PolyA deltaPI density in cortex and WM. **c)** Downsampling experiments which equally sampled from individuals equally and selected 20 reads and 50 genes randomly. Genes were resampled 100x and calculated percent significant per iteration. **d)** ScisorWiz^48^ example of alternative polyA site in the VBP1 gene. **e)** Average length of last exons per tested genes with |deltaPI| <0.2 compared across regions and age groups. **f)** Average length of last exons per genes with |deltaPI| >0.2 compared across regions age groups. WM is colored in blue and Cortex is colored in red. **g)** Average length of UTR per genes with |deltaPI| >0.2 compared across regions age groups. **h)** Overview of all cortical overlapping genes that were tested in both Poly(A) and exon tests (n=2,096). Genes were classified as only having |deltPI| > 0.2, only having a |deltaPSI|>0.2, or both.

Of note, although many of the total barcoded, spliced reads have an annotated poly(A) site, only 6.35% have a TSS, making the ability to do full-isoform tests challenging. By employing our long-molecule capture we have increased the length and % transcript covered but still lack a majority of full-length cDNAs. Even so, we performed full-length isoform tests on the full-length reads and confirmed that the number of significant genes with alternative-isoform usage per comparison is lower than on an individual exon basis (**Supplemental Table 4**). When comparing the child and young adult cortex, we identify 266 genes which exhibit alternative isoform usage between age groups. Of these 266 genes, 90 contain significantly altered exons from our primary analysis, where 30 genes are associated with two or more. Additionally, we find similar trends to alternative exons and Poly(A) sites demonstrating that cortex contains more significant isoform alternative usage than WM across age (Cortex: 25.32%, 95%-confidence intervals [13.40,37.24]; WM: 2.78%, 95%-confidence intervals [1.70,3.85]; **Supplemental Figure S26a**), although we note that these values are not downsampled. An example of alternative-isoform usage in cortex can be found in M-Phase Phosphoprotein 6 (*MPHOSPH6*), a gene which is involved in the RNA exosome process^64^. In this example, one isoform is primarily used in the child cortex, whereas another isoform is primarily used in the young adult cortex (**Supplemental Figure S26b**). Thus, Spl-ISO-Seq enables the capture of full-length isoforms and the identification of differences in isoform usage, albeit at a sparser rate than at the individual-exon level.

### Among cortical layers, layer 4 shows the strongest splicing alterations

We divided the cortex into three well-distinguishable compartments: Layers 1-3 (“L1-3”), layer 4 (“L4”) and layers 5-6 (“L5-6”). We found many altered exon-inclusion events in each compartment between age groups (**Figure 6a-c**). Testing for differentially spliced exons, L4 had the highest fraction of developmentally regulated exons: 13.57% of tests were significant for L1-3 (95%-confidence interval [12.84,14.30]), 16.12% for L4 (95%-confidence interval [15.44,16.77]), and 12.45% for L5-6 (95%-confidence interval [11.85,13.02]) (**Figure 6d**). L1-3 and L4 showed approximately similar numbers of up- and downregulated exons. Surprisingly however, L5-6 revealed a markedly different trend: alternative exons were frequently downregulated after puberty (**Figure 6e**; L1-3: Mean=0.03; SD=0.44; L4: Mean=0.03; SD=0.42; L5-6: Mean=-0.04, SD=0.43). Downsampling experiments also enforcing that no single or pair of individuals can dominate the data for a given exon, showed that this observation is robust to equalizing statistical power (corrected 2-sided Wilcoxon rank sum test; L1-3 V L4 FDR=0.0024; L4 V L5-6 FDR=0.0037; L1-3: Mean=0.33, SD=0.53, L4: Mean=0.7, SD=0.82; L5-6: Mean=0.39, SD=0.68; **Figure 6f**). As an example, three children show strong exon inclusion, while three young adults exhibit mostly exon exclusion in layer 4 of the cortex in the *TSPAN19* gene (**Figure 6g,h**). SynGO analysis revealed no significant enrichment of any synaptic compartments in altered exon inclusion in L1-3 (**Supplemental Figure S27a**). However, in L4 developmentally regulated splicing, we found enrichment of the postsynaptic compartment and to a lesser degree as well in L5-6 (**Figure 6i, Supplemental Figure S27b).** Similarly to observations on alternative exons (see Figure 5 and prior panels of Figure 6), we found L4 to have higher percentages of significant alternative-acceptor (**Supplemental Figure S28a;** L1-3 95%-confidence interval [1.86, 3.21]; L4 95%-confidence interval [3.46, 5.12]; L5-6 95%-confidence interval [1.68, 2.91]) as well as poly(A)-site regulation in the considered developmental period (**Figure 6j**; L1-3 95%-confidence interval [15.88,18.48]); L4 95% confidence interval [20.01, 22.63]; L5-6 95%-confidence interval [14.35, 16.65]; **Supplemental Figure S29**). However, we note that these values were not down sampled due to a lack of sufficient data amounts. For alternative donors, we found a visually similar trend, however with overlapping 95%-confidence intervals (**Supplemental Figure S28b**; L1-3 95%-confidence interval [2.36, 3.93]; L4 95%-confidence interval [3.52, 5.31]; L5-6 95%-confidence interval [2.86, 4.47]). Layer 4 similarly showed increased changes to Layer 5-6 in full-length isoform usage (**Supplemental Figure S26c**). We had the largest amount of overall read numbers for excitatory neurons, which enabled downsampling experiments using exclusively spots that are representing excitatory neurons only. These downsampling experiments revealed that the higher ratio of splicing regulation in layer 4 is confirmed when only considering excitatory neurons (L1-3: Mean=0, SD=0; L4: Mean=0.26; SD=0.44; L5-6: Mean=0, SD=0; corrected 2-sided Wilcoxon rank sum test; L1-3 v L4 FDR=5.0×10^-8^; L4 V L5-6 FDR=5.0×10^-8^; **Figure 6k**). We then compared the lists of genes with altered alternative splicing during puberty to published lists with disease-associated splicing. For a published list of genes with autism-spectrum disorder-associated splicing changes^12,65,66^, we found a higher odds-ratio for L1-3 compared to L5-6, with non-overlapping 95% confidence intervals. The L4 odds ratio was in between and had overlapping 95% confidence intervals with both L1-3 and L5-6 (L1-3 95%-confidence interval [1.42, 2.30]); L4 95%-confidence interval [1.06, 1.58]; L5-6 95%-confidence interval [0.77, 1.20]; **Figure 6l**). Similarly investigating gene lists of amyotrophic lateral sclerosis (ALS)^67^, schizophrenia^68^ and Alzheimer’s disease (AD)^11^, we found non-significant associations for all diseases and layers, when correcting for the three tests performed per disease (**Supplemental Figure S30**). We next examined if any repetitive elements existed either upstream or downstream of developmentally-regulated exons. We found that while there was no increased abundance of repetitive elements upstream (2-sided Fisher test: p=0.334; **Supplemental Figure S31**), the regulated exons had significantly more repetitive elements downstream compared to a background set (2-sided Fisher test: p=0.004; **Figure 6m**; Methods). When investigating if this effect was driven by one or multiple types of repetitive elements, we found that LTR/ERV1 repeats out of the most abundant elements were increased downstream of developmentally-regulated exons (corrected 2-sided Fisher test: FDR=0.028). The three other types of repeats we monitored did not reach significance on their own but suggested a similar trend (**Figure 6n**).

**Figure 6.**
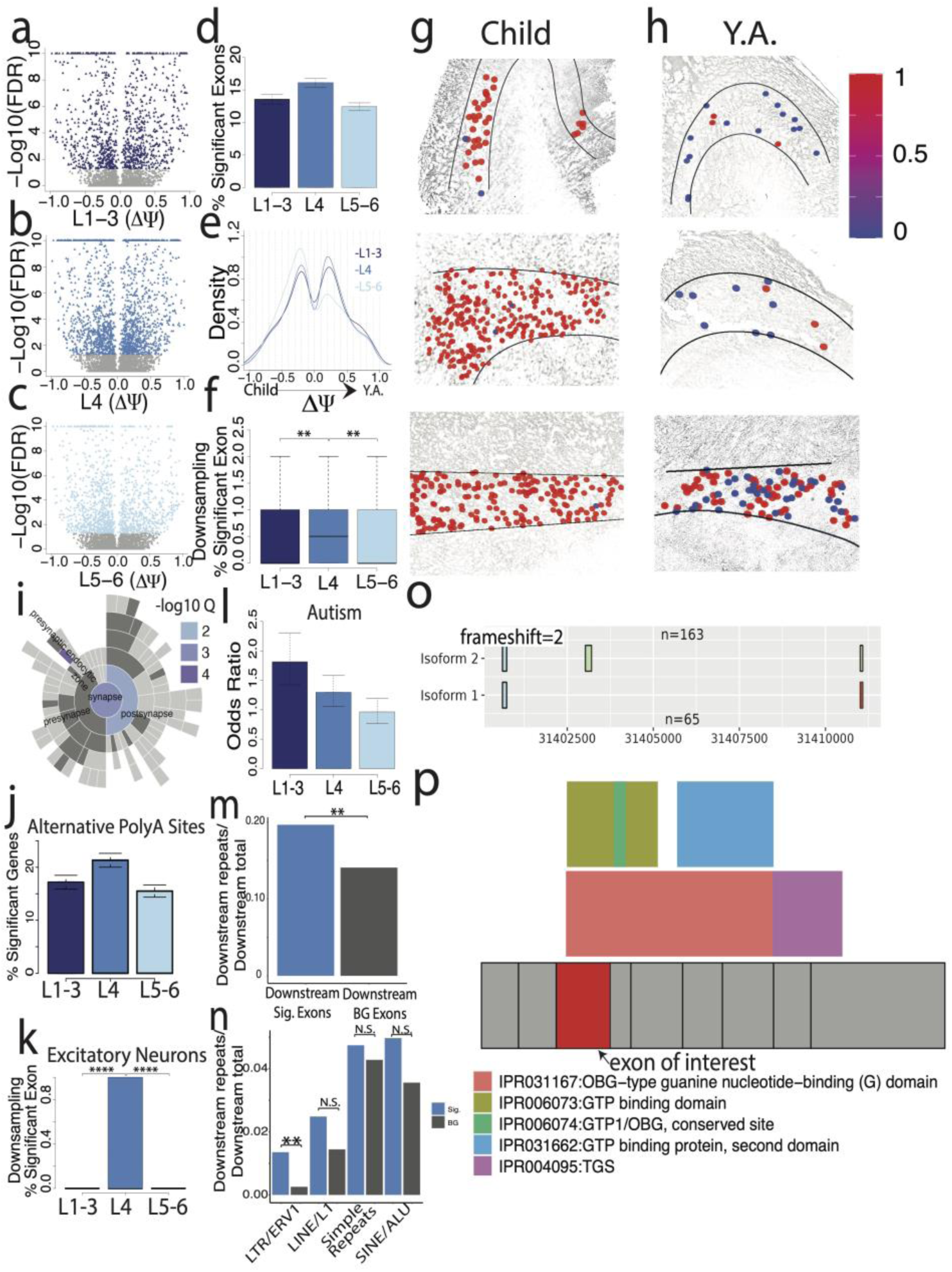
**a)** Exons tested between ages in L1-3 by deltaPSI (Y.A.-Child) and -log10(FDR). Colored points are significant exons and grey points are non-significant. **b)** Exons tested between ages in L4 by deltaPSI and -log10(FDR). Colored points are significant exons and grey points are non-significant. **c)** Exons tested between ages in L5-6 by deltaPSI and -log10(FDR). Colored points are significant exons and grey points are non-significant. **d)** Percent significant of tested exons and |deltaPSI| >=0.2 for L1-3, L4, and L5-6. **e)** deltaPSI density of significant exons for L1-3, L4 and L5-6. **f)** Percent significant exons from downsampling experiments. Exon selection and percent significant calculations are repeated 100x and plotted. **g)** Exon chr12_85027899_85028023_- in the *TSPAN19* gene plotted by area in 3 child samples. **h)** Exon chr12_85027899_85028023_- in the *TSPAN19* gene plotted by area in 3 Y.A. samples. **i)** SynGO location enrichment of significant L4 genes. **j)** Percent significant genes with alternative polyA sites by layer. **k)** Downsampling of excitatory neuron specific reads and genes. **l)** Odds ratio comparing significant group v background group of autism associated genes. **m)** Ratio of sequences with repetitive elements found to total number of sequences per group. Sig: Sequences found upstream of exons with |deltaPSI| > 0.5 and FDR < .05. BG: Sequences found upstream of exons with |deltaPSI| < 0.05 and FDR > .05. **n)** Ratio of sequences with repetitive elements found to total number of sequences per group, broken down by repetitive element. **o)** RiboSplitter plotted example of an exon skipping event in the DRG1 gene. Blue squares indicate constitutive exons, green square indicate alternative exon. “n” indicates number of reads in each isoform. **p)** DRG1 gene plotted where exons are denoted in grey squares. Colored boxes indicate protein domains and their locations relative to exons.

Additionally, RiboSplitter^69^ was used to identify protein domains associated with spliced L4 genes. RiboSplitter requires another alternative splicing analysis software, Spladder^70^, to identify alternative splicing events and their related protein domains. Spladder identified 94 exon-skipping events across development in Layer 4. Among these 94 exons, we considered those that are also tested in our data for correlation analysis. DeltaPSIs from the two sources correlate highly (Spearman’s R = 0.83, **Supplemental Figure S32a**). In addition to the 94 exon skipping events, RiboSplitter also identified alternatively included retained introns, 3’, 5’, mutually exclusive exons, and multiple skipped exons between age groups (**Supplemental Figure S32b**). Of these, 47.2% led to a frameshift mutation. The PDZ domain (IPR001478) was most affected by exon skipping and no domain was most prominently affected by intron retention (**Supplemental Table 5**). Of note, PDZ domains are often found in scaffolding proteins which help to organize and shape the postsynaptic landscape^71–73^. An example of an alternative exon which overlaps with a protein domain can be seen in the Developmentally Regulated GTP Binding Protein 1 (*DRG1*) gene. In this example, the highlighted exon is 59% more included the young adult age group compared to the child group, where a total of 163 reads contain the exon and 65 skip it (**Figure 6o**). Of note, the removal of this exon changes the reading frame of this protein, and the exon itself spans several protein domains, including OBG-type guanine nucleotide-binding (G) domain and GTP binding domain (**Figure 6p**). The *DRG1* gene has been implicated with microtubule dynamics^74^, ribosome regulation^75,76^, as well as several developmental neurological disorders including autism spectrum disorder^77^ and microcephaly^78^.

## DISCUSSION

Developmental changes happening around puberty are known to affect cortical plasticity. However, splicing and poly(A)-site changes in specific layers and cell types, and if these spatially defined changes contribute to neuronal plasticity equally, had not been mapped out completely. Here, we develop a new technology of spatial isoform sequencing at near-single-cell resolution (Spl-ISO-Seq) as well as a software suite (Spl-IsoQuant) enabling barcode deconvolution and isoform expression analysis. In general, the ability to capture single cells with most spatial technology may be limited to the size of the cells within the tissue being studied. Here, we use human brain tissue which contains an increase in neuronal cell size and density compared to other mammals, where neuronal soma volume can be greater than 2000 um^3 79–81^. The literature gives abundant examples of alternative splicing influencing synaptic function, transmission, and plasticity^82–86^. However, which splicing events are altered in puberty and in which cell types and which cortical layers is much less studied. Armed with the above technological advances we find that the cortical layers overall show higher developmental splicing changes than the white matter and that these splicing changes harbor enrichment for different types of protein domains. We also noticed that cortical genes containing developmentally regulated exons are enriched for developmentally regulated poly(A)-site usage as well. However, for oligodendrocytes, we find the opposite trend: Higher splicing regulation occurs in the white matter than in the cortex. This raises the compelling question of which variable may be influencing oligodendrocyte splicing in this region-specific way, which could include microenvironmental factors or unique cell-type interactions.

Comparing distinct layers, L4 stands out in developmental regulation of alternative-exon inclusion, alternative-acceptor usage, and Poly(A)-site usage. Results for alternative donors suggest a similar observation, which is however non-significant. This emphasizes the need for the exciting future direction of monitoring all RNA variables, including splice sites, Poly(A) sites, TSS sites, and RNA modifications during this developmental period. For alternative exons, the importance of L4 in developmental regulation can be traced to excitatory neurons – the most abundant data source in this spatial experiment. We also find in layer 4 that the regions downstream of developmentally regulated exons are enriched with repetitive elements, an observation which is driven by LTR/ERV1 repeats. Literature suggests that these repeats could be influencing splicing themselves, or perhaps act as a binding site to RNA binding proteins or other regulatory elements, as it’s known that many RNA binding proteins bind to repeats^87–91^. Interestingly, these repetitive elements have been shown to be dynamically expressed throughout development in the brain^92,93^. Overall, these findings show that LTR/ERV1s co-localize with developmentally-regulated exons. This observation suggests the exciting hypothesis that these repeats may play an important role in alternative splicing in the context of human cortical development, suggesting multiple future research directions. Importantly, L4 is distinct from other cortical layers as it receives signals from the thalamus, whereas other layers generally assist in further sensory processing. Additionally, L4 has a unique structure consisting of ocular dominance columns. Our results align with previous research demonstrating that L4 in the visual cortex exhibits increased synaptic plasticity compared to other layers^94–96^ especially during the critical period of sensory development^97–99^. Overall, we find that the splicing changes detected during puberty are linked to synaptic and especially post-synaptic biology as well as autism spectrum disorder. For this disorder L1-3 have stronger association than L5-6, with L4 showing an in-between result, suggesting layer-specific effects of splicing in ASD. We do not find similar associations to AD and ALS – which could be due to these diseases being associated to older ages or different cortical regions and not the period or anatomical area we investigate here – or because of differences in how distinct prior publications derived these gene lists. Of note, we also do not find an association to schizophrenia. A biological explanation may be that schizophrenia arises usually after the age of 20, which is past the age-range we investigated here, while autism arises earlier in childhood. However, technical differences in how the schizophrenia gene list was derived cannot be ruled out. Of note, the autism association of isoform usage is at this point not yet rooted in a specific cell type. The literature suggests that autism is linked to an imbalance of excitation and inhibition, perhaps driven by inhibitory neuron dysfunction^100–102^. Our data is dominated by excitatory neurons – with little conclusions to be drawn about inhibitory neurons. There are two possible, non-mutually exclusive, explanations for our observations. First, inhibitory neuron dysfunction could affect excitatory neurons and their splicing program due to changes in activation patterns. Second, isoforms could be altered in similar ways in both excitatory and inhibitory neurons but have stronger functional effects in inhibitory neurons. At the moment, this technology requires sufficient amounts of data between comparisons which limits the ability to investigate intra-layer variability, especially at the cell-type level or even between two individual cells. Enhancing the rate of assigned singlets and UMIs per barcode captured will enable this future analysis and greatly inform the fields of neuroscience and spatial transcriptomics. Of note, cell-type information was determined with short-read sequencing of fragmented cDNAs due to enhanced read depth and gene expression profiles using short reads. Long-read sequencing (here of unfragmented cDNAs), although necessary to describe transcript information spanning multiple exons, generally is not sufficient for gene-expression based analyses (unless multiple flow cells are used to match short-read depth per sample). Vice versa, short-read sequencing is not sufficient to identify alternative exon inclusion nor characterize fully synthesized molecules. Thus, both sequencing modalities are required in tandem for Spl-ISO-seq.

Additionally, while this technology increases the average length and transcript coverage per read compared to the standard, most reads are not full-length. This drawback limits the extent to which we can describe 5’ TSSs and full-length isoform changes across development. Other technologies, including but not limited to 10X Genomic’s Visium, DBiT-Seq, and StereoSeq, use a transcript switch oligo (TSO) rather than a universal priming site (UPS) during 2^nd^ strand synthesis which would enable full-length molecule capture. However, other variables including spatial resolution (spot size), molecule diffusion, and capture efficiency should also be considered when choosing a spatial transcriptomic technology. Future work should focus on optimizing high-resolution and low diffusion spatial technology for mapping not only of full-length isoforms across a region, but also other modalities at the same time. This would propel the field forward and enable the investigation of multi-modal networks in relation to their spatial location and microenvironments.

Overall, we demonstrate a near-single cell spatial transcriptomic technology which maximizes transcript coverage and employ it to better understand alternative splicing in the developing visual cortex. Our new technologies enable a view of splicing changes that can at once be rooted in specific cell types as well as specific areas or structures within anatomical regions. Thus, there are wide applications for this technology in cancer, neurodegeneration, neuropsychiatric disorders and many other diseases.

## DATA AVAILABILITY

All data used for this study are uploaded to the National Institute of Health’s Sequence Reads Archive (SRA) under submission PRJNA1116561. A reviewer link can be found at: https://dataview.ncbi.nlm.nih.gov/object/PRJNA1116561?reviewer=uc7v6e10toqp4lllatmu6vvf53

## CODE AVAILABILITY

The package Spl-Isoquant is available at https://github.com/algbio/spl-IsoQuant.

## CONFLICTS OF INTEREST

H.U.T. has presented at user meetings of 10x Genomics, Oxford Nanopore Technologies, and Pacific Biosciences, which in some cases included payment for travel and accommodations. The other authors declare no competing interests.

## ACKNOWLEDGEMENTS

We thank Olivia Spicer and the NIH NeuroBioBank for human tissues. We thank Adrian Tan, Chendong Pan, Aihong Liu, Seongeun Oh and Jenny Xiang from the Genomics Resources Core Facility at Weill Cornell Medicine for performing RNA sequencing. We thank Dr. Christopher Mason for use of his PromethION machine. We also thank Weill Cornell Medicine Scientific Computing Unit (SCU) for use of their computational resources.

Supported by: NIGMS 1R01GM135247-01 (H.U.T), MIRA R35 GM152101-01 (H.U.T), Brain Initiative grant 1RF1MH121267-01 (H.U.T.), NIDA U01 DA053625-01 (H.U.T among others), NIDA grant 2T32DA039080 (J.H., N.B.), NSF GRFP # 2139291 (C.F.), the Feil Family Foundation (H.U.T.), European Research Council (ERC) European Union’s Horizon 2020 research and innovation programme (grant agreement No. 851093, SAFEBIO) (A.D.P, A.I.T.), Research Council of Finland grants No. 322595, 346968, 358744 (A.D.P, A.I.T.), R01HD111089 (M.E.R.), NLM 1R01LM014017-01 (D.K.), ERC ASTRA_855923 (G.G.T.), National Center for Gene Therapy and Drugs based on RNA Technology (CN00000041), financed by NextGenerationEU PNRR MUR – M4C2 – Action 1.4-Call “Potenziamento strutture di ricerca e di campioni nazionali di R&S” (CUP J33C22001130001) (G.G.T., A.V.), MIRA R35 GM138152 (I.H.).

## Spl-ISO-seq Methods

### Human Brain Tissue Acquisition

The fresh frozen human brain tissues of 4 children (ages 8-11) and 4 young adults (ages 16-19) were obtained from the NIH Neurobiobank.

### Spatial Barcoding and cDNA Synthesis

Tissue was embedded in Tissue-Tek OCT Compound (Sakura, catalog no. 4583) and stored in -80C overnight. Tissue blocks and CryoCube (Curio Bioscience part no. JW001) were placed in a -20C Leica CM3050 S cryostat for 20 minutes prior to sectioning. Tissue was sliced at 10 uM thickness. Tissue was positioned and melted over a Curio Seeker 3×3 mm slide (Curio Bioscience part no. SK004). CryoCube was sliced at 30 uM thickness and positioned and melted on top of tissue on seeker slide. Tissue hybridization, reverse transcription, tissue clearing and bead resuspension, 2^nd^ strand synthesis, and cDNA amplification were performed according to the Curio Seeker 3×3 spatial mapping kit v 2.1. Glass slides with tissue were immediately placed into 1.5 mL tubes with 200 uL hybridization mix (190 uL hybridization buffer [Curio Bioscience part no. B005] & 10 uL RNAse Inhibitor [Curio Bioscience part no. E001]) for 15 minutes at room temperature. Following hybridization, slides were dipped into RT wash buffer (40 uL RT/SS buffer [Curio Bioscience part no B006] & 160 uL nuclease free water [Thermofisher catalog no. AAM9937]) for 3 seconds and transferred to tubes with 200 uL RT reaction mix (40 uL RT/SS buffer, 20 uL dNTPs [Curio Bioscience part no. N001], 5 uL RNase inhibitor, 10 uL RT Enzyme [Curio Bioscience part no. N001 E003], 125 uL Nuclease-free water) for 10 min at room temperature followed by 30 minute incubation at 52 C. 200 uL of Tissue clearing reaction mix (196 uL TC buffer[Curio Bioscience part no. B004], 4 uL TC Enzyme[Curio Bioscience part no. E002])) was added to tubes and incubated for 30 minutes at 37C. 200 uL bead wash buffer (Curio Bioscience part no. B003) was added to tubes and barcoded beads were dissociated from the glass slide by gently pipetting the mix over the area. Once beads were fully dissociated, the glass slide was removed from the tube. Beads were washed by centrifuging for 3000g for 2 minutes, supernatant removed, and resuspended in 200 uL bead wash buffer. This washing cycle was repeated once more prior to second strand synthesis. Beads were then incubated at 95 C for 5 minutes and washed. 200 uL second strand mix (40 uL RT/SS buffer, 20 uL dNTP, 2 uL SS Primer [Curio Bioscience part no. P002], 5 uL SS Enzyme [Curio Bioscience part no. E007], 133 uL Nuclease-free water) was added and incubated at 37C for 1 hour. Beads were then washed 2x and resuspended in cDNA amplification mix (100 uL cDNA amp buffer [Curio Bioscience part no. B007], 8 uL cDNA amp primer mix [Curio Bioscience part no. P003], 4 uL cDNA amp enzyme [Curio Bioscience part no. E006], 88 uL Nuclease free water). A PCR with the bead mix was conducted as follows: 98C for 2 minutes, followed by 4 cycles of 98 C for 20 seconds, 65 C for 45 seconds, and 72C for 3 minutes, an additional 14 cycles of 98C for 20 seconds, 67 C for 20 seconds, 72 C for 3 minutes, followed by 72 C for 5 minutes. cDNA was cleaned up with 0.6X SPRIselect beads, incubated for 5 min at room temperature, and washed 2x with 80% ethanol. SPRIselect beads were eluted in 50 uL nuclease free water and cleaned up with another round of 0.6X SPRIselect beads, ethanol washes, and elution. cDNA concentration was measured with Qubit and cDNA length was measured with Tapestation.

### Tagmentation and Illumina Library Preparation

5 ng of cDNA from the Curio seeker protocol was eluted to 5 ul with nuclease free water. Illumina libraries were prepared using the Nextera XT Library Kit (Illumina, part no. FC-131-1024) and followed protocol as provided from Curio Biosciences. 10 uL of TD and 5 uL ATM were added to the tube and incubated at 55 C for 5 minutes. Immediately after 5 uL of NT was added to the mix and incubated at room temperature for 5 minutes. 15 ul of NPM and 5 uL of a unique F and R primer pair were added per sample. A PCR with the mix conducted as follows: 72 C for 3 minutes, 95 C for 30 seconds, followed by 12 cycles of 95 C for 10 seconds, 55 C for 30 seconds, and 72 C for 30 seconds, followed by 72 C for 5 minutes. Library was cleaned up with 0.6X SPRI beads, washed 2x with 80% ethanol, and eluted in 50 ul TE. 0.8x SPRI beads were added to the elution and washed with ethanol 2x and eluted in nuclease free water. cDNA concentration was measured with Qubit and cDNA length was measured with a tapestation. Nextera XT libraries were sequenced with the Illumina Nova Seq X Plus.

### Exome Enrichment of Spatial cDNA

The following steps were performed during exome capture. Exome capture was performed as previously described to selectively amplify spliced molecules^1^. Agilent probe kit SSELXT Human All Exon V8 (Agilent, 5191-6879) and the reagent kit SureSelectXT HSQ (Agilent, G9611A) were used according to the manufacturer’s manual. First, the block oligonucleotide mix was made by combining 1 ul of PCR handle primer (5’-CTACACGACGCTCTTCCGATCT-3’) and 1 ul of UPS primer (5’-AAGCAGTGGTATCAACGCAGAGT-3’) at 200 ng/ul. Next, 5 ul of 50-100 ng/ul spatial untagmented cDNA diluted from the first step was diluted with 2 ul blockmix and 2 ul nuclease free water (NEB, AM9937). The mix was incubated on a thermocycler under the following conditions: 95 °C for 5 min, 65 °C for 5 min and 65 °C on hold. Next, hybridization mix was made by combining 20 ul of SureSelect Hyb1, 0.8 ul of SureSelect Hyb2, 8.0 ul of SureSelect Hyb3 and 10.4 ul of SureSelect Hyb4 at room temperature. Once the cDNA block mix reached to 65 °C on hold, 5 µl of probe mix, 1.5 µl of nuclease-free water, 0.5 µl of 1:4 diluted RNase Block and 13 µl of the hybridization mix were added to the cDNA block oligo mix and incubated for 24 h at 65 °C. M-270 Streptavidin Dynabeads (Thermo Fisher Scientific, 65305) were prepared by washing three times and resuspended with 200 µl of binding buffer. Once the incubation period ended, cDNA block oligo mix was let sit at room temperature and then mixed with M-270 Dynabeads. The bead-cDNA mix was placed on a Hula mixer for 30 minutes at room temperature. During the incubation, 600 µl of wash buffer 2 (WB2) was transferred to three wells (200 ul each) of a 0.2-ml PCR tube and incubated in a thermocycler on hold at 65 °C. After the 30-min incubation, the bead buffer was replaced with 200 µl of wash buffer 1 (WB1). Then, the bead-cDNA mix was put back into the Hula mixer for another 15-min incubation with low speed. Next, the WB1 was replaced with WB2, and the tube was transferred to the thermocycler for the next round of incubation (10 minutes). WB2 was replaced 2 more times at 10 minute intervals with the pre-heated WB2. Following incubation, beads were resuspended in 18 ul nuclease free H20 and then the spliced cDNA, which is attached to the beads, was amplified as described below.

### Long-cDNA Capture and Amplification

The spliced cDNA captured by exome probes, which bound with the M-270 Dynabeads, was amplified with primers (UPS: 5’-AAGCAGTGGTATCAACGCAGAGT-3’; PCR: 5’-CTACACGACGCTCTTCCGATCT-3’) and KAPA-HiFi enzyme by using the following PCR protocol: 95°C for 3 min, 8 cycles of 98°C for 20 s 64°C for 60 s and 72°C for 3 min. The amplified probe captured cDNA was isolated from M-270 beads as supernatant and followed by a size selection/purification with 0.48× SPRIselect beads (in 1.25M NaCl-20% PEG buffer) and then eluted in 25ul EB buffer. All 25 ul size-selected spliced cDNA was used as template for the second round amplification of 6-8 cycles with the same PCR conditions suggested above. The product of the second round PCR was size selected/purified with 0.48× SPRIselect beads (in 1.25M NaCl-20% PEG buffer) and then eluted in 50 ul EB buffer.

### Oxford Nanopore Library Preparation

For all samples, ∼75 fmol exome enriched and length optimized cDNA underwent ONT library construction by using the Ligation Sequencing Kit (ONT, SQK-LSK114), according to the manufacturer’s protocol (Nanopore Protocol, Amplicons by Ligation, version ACDE_9163_v114_revO_29Nov2022). The ONT library was loaded onto a PromethION sequencer by using PromethION Flow Cell (ONT, FLO-PR114M) and sequenced for 72 h. Base-calling was performed with Super-high accuracy settings on the Oxford Nanopore MinKNOWUI dorado basecaller.

### Hematoxylin and Eosin staining

The 10uM tissue following the experimental tissue slice for each sample was placed on a colorfrost plus slide (Epredia catalog no. 6776214) and fixed in 4% PFA (Electron Microscopy Sciences catalog no. 15710) for 30 minutes on ice. Slides were washed in 1x PBS 3x for 5 minutes each. H&E staining kit (abcam catalog no. ab245880) was used to stain slides. Incubation times for Hematoxylin and Eosin were optimized per sample using excess tissue. Slides were imaged on a Leica CTR 6500.

### Short Read Data Processing and deconvolution

Illumina data was analyzed using the Curio Seeker Bioinformatics pipeline (https://curiobioscience.com/bioinformatics-pipeline/). Resulting .rds files were inputted into Robust Cell Type Decomposition (V2.2.1) program^2^ with a reference file of including 7 human single nuclei PFC samples with annotated clusters/celltypes including excitatory neurons, inhibitory neurons, oligodendrocytes, oligodendrocyte precursor cells, astrocytes, microglia, vascular endothelial cells, T cells, fibroblasts, and unknown. Basic RCTD parameters were used in “doublet” mode following published tutorials (https://github.com/dmcable/spacexr/blob/master/vignettes/spatial-transcriptomics.Rmd). Spots designated as “singlets” were assigned as cell types per sample.

### Identifying Cortical Layers

Cortical layers were identified by data from combining H&E images, excitatory neuron density based on RCTD singlets, and known gene expression markers which are layer specific or white-matter specific. These were assigned on a slide-by-slide basis using established visual cortex architecture as a guide.

### Detecting differential expressed genes

For each age group and cell type, the set of differential expressed genes (DEG) detected from the comparison was obtained by running FindMarkers function of Seurat^3^. The GO enrichment analysis for the DEGs was performed by using enrichGO or enrichKEGG function of clusterProfiler^4^ 4.2.2.

### Long Read Data Processing

#### Spl-IsoQuant overview

To process spatial long-read sequencing data, generated from the previously mentioned methods, we developed a novel pipeline Spl-IsoQuant, which is based on IsoQuant — the software for bulk long-read data analysis previously developed in the lab^5^. Spl-IsoQuant starts with barcode calling to identify barcode and UMI sequences in every read. Further, reads are mapped to the reference genome with minimap2 in spliced mode^6^ and assigned to known genes and isoforms using existing IsoQuant algorithms. Further, PCR duplicates are removed using the obtained gene assignments and identified UMI sequences. The resulting collection of reads represent unique known mRNA molecules and their respective cells, which is further used for analysis of polyA sites, TSS and splicing patterns.

Spl-IsoQuant is an open source software and is openly available at https://github.com/algbio/spl-IsoQuant.

### Barcode detection

To identify linker, primer and barcodes positions in a sequenced read we implement a pattern matching algorithm based on short k-mer indexing and conventional Smith-Waterman local alignment. Given a set of known sequences (e.g. barcode whitelist, a linker or a primer), a k-mer index is a key-value hash table containing all possible k-mers as keys and a set of known sequences containing this k-mer as a respective value. Thus, to quickly detect a presence of a known sequence in the read, one needs to split the read (or its region) into k-mers and extract their corresponding values for the index. We construct 3 distinct k-mer indices: for the primer sequence (k=6), the linker (k=5) and the last one for the list of all known barcodes (k=6). Small k values are used to be able to detect exact k-mer matches even when the read has sequencing errors. However, short k-mer matches can be accidental, especially for a barcode whitelist containing tens of thousands of barcodes. Thus, a k-mer index can only be used to obtain a set of preliminary candidate sequences. Further, we use the classic Smith-Waterman (ssw-py library^7^) local alignment algorithm to check which k-mer matching derived candidates are actually present in the read and filter out arbitrary matches. Exploiting the k-mer index approach allows to shrink the search region when matching the primer and the linker, as well as greatly reduce the number of Smith-Waterman calls when detecting barcode sequence.

To detect barcodes and UMIs, reads are processed individually using the following procedure. First, a poly-T stretch is detected using a sliding window technique. Further, the primer and the linker positions are identified as described above in the region of the read preceding the polyT stretch. Primer and linker coordinates are used to extract a potential barcode sequence (Fig. 3b). This sequence is further split into k-mers and possible matching barcodes from the whitelist are detected using the barcode k-mer index. Among those, the candidate with the highest Smith-Waterman local alignment score is selected as the barcode for this read. If the best alignment score is below a certain threshold, the candidate barcode is considered to be unreliable and is not reported. We exploit the classic local alignment scoring system that adds 1 for a symbol match and -1 for mismatch or indel, the default alignment score cutoff is 13. Although this scheme does not allow errors in the middle of the barcode, it does consider the possibility of terminal indels/substitutions, including ones caused by potential inaccuracy in barcode positioning. UMI is simply extracted from the read between barcode end and polyT start. If no polyT stretch, linker, primer or barcode are found, the above procedure is repeated for the reverse-complemented read sequence. If no reliable barcode is detected for both forward and reverse sequence, no barcode and UMI are reported and the read is ignored in the further analysis.

The described algorithm is implemented in Python with the use of the SSW library^7^ and is as a part of the Spl-IsoQuant pipeline. An ONT dataset containing 85 million reads is processed in approximately 2 hours using 20 threads. In contrast, mapping the same dataset with minimap2 takes about 18 hours, and the remaining part of the pipeline (read-to-isoform assignment and UMI deduplication) takes roughly 4 hours on the same dataset.

#### Read processing and UMI deduplication

Once barcodes are detected, the reads are mapped to the reference genome using minimap2 with the default setting used in IsoQuant (minimap2 -x splice --junc-bed annotation.bed -a --MD -k 14 -Y --secondary=yes). Spl-IsoQuant then assigns all mapped reads to the reference genes and isoform using the algorithm described in (Prjibelski et al., 2024). Multimapped reads are resolved by selecting the alignment that is the most consistent with the reference annotation or discarded. Reads that were uniquely assigned to a known gene are further processed to remove PCR duplicates.

To filter out PCR duplicates, reads are grouped by the assigned gene and barcode. Thus, reads in the same group represent mRNAs originating from the same cell and the same gene. If a pair of reads in the group contain identical UMI sequences, it is highly likely they were sequenced from PCR duplicates of the same mRNA molecule. However, due to the presence of sequencing errors we consider reads to be PCR duplicates if their UMI sequences have a certain level of similarity, namely they are edit distance 2 apart or less (starting and trailing indels are not considered). To identify PCR duplicates within a set of reads having the same barcode and the same assigned gene, we iterate over these reads one by one and collect a set of “representative” UMIs (empty at the beginning of iteration). If a read contains a UMI that is similar to any of the collected representative UMIs, this read is assigned to the most similar representative UMI. Otherwise, the UMI is added to the representative set and the read is assigned to its own UMI. Although this algorithm might not be exact and requires quadratic number of UMI comparisons in the worst case, in practice it allows avoiding computing all-vs-all pairwise edit distances.

Further, among reads assigned to the same representative UMI we select the one covering the highest number of splice junctions. Other reads assigned to this representative UMI are deemed to be PCR duplicates and thus are further ignored. UMI-filtered reads are used in the downstream analysis, such as exon and transcript quantification and differential splicing analysis.

Indeed, it is possible that similar UMIs were attached to two distinct mRNA molecules and the described procedure will remove a read which does not have a PCR duplicate. However, due to randomness of UMI sequences (∼150K distinct possible UMIs of length 9 bp with last to non-T bases), such an event is rare and is unlikely to affect the downstream analysis.

### Benchmarking spl-IsoQuant

To assess the designed barcode calling algorithm we simulated Nanopore data with Trans-NanoSim^8^. We used a forked version of Trans-NanoSim (https://github.com/andrewprzh/lrgasp-simulation) with the improved read truncation procedure, which allowed us to simulate data with realistic ONT truncation probabilities, as well as reads with no sequence truncation. This version of Trans-NanoSim was previously used to benchmark IsoQuant^5^. The NanoSim error model was trained using actual sequencing data produced in this project (sample 329) exhibiting a sequencing error rate of 4.4% (1.4% mismatches, 2.0% insertions, 1.0% deletions). Therefore, recall and precision estimated with simulated data accurately represents a real life situation.

Molecule templates provided to Trans-NanoSim were generated by concatenating a reference transcript sequence with 30bp poly(T) stretch, a random UMI, an arbitrary barcode from the whitelist, the linker and the primer according to the molecule scheme (Fig. 2h). Reference transcripts were randomly selected according to the expression profile that was also derived from the same sample (329). 5 million distinct molecule templates were generated, featuring ∼70,000 distinct barcodes. In total, 20M Nanopore reads were simulated for the benchmarking.

We Spl-IsoQuant ran on the simulated dataset and detected barcodes were compared against true barcodes attached to mRNA sequence during the simulation. In this procedure, correctly called barcode is regarded as true positive, while wrongly reported barcode — as false positive. Reads, for which no barcode is detected due to, for example high number sequencing errors or read truncation, are considered as false negatives.

### Calling differentially included exons

After Spl-IsoQuant was run on the long read data and identified reads mapped, barcoded, and spliced reads with corrected UMIs, an AllInfo file was generated containing the following information: Read name, Gene, Group for comparison (ie. cell type or region), barcode, UMI, intron chain, TSS site, PolyA site, exon chain, known or novel transcript, and number of introns. More information regarding AllInfo files can be found at https://github.com/noush-joglekar/scisorseqr^9^. We separated AllInfo files by which comparisons were being performed. For example, if the comparison of child cortex vs. young adult cortex was being performed, reads which aligned with each comparison were identified and placed into 2 distinct AllInfo subfiles. These subfiles were run in the casesVcontrols() function of the scisorATAC R package with minimum reads required per exon set to 10, in order to identify which exons are differentially included between the 2 groups using basic parameters.

First, we identify alternative exons using reads from both groups that are being compared (e.g., layer4 barcodes from young individuals and layer 4 barcodes from old individuals). This uses only reads that

- Have introns
- Respect the splice site consensus at all of their introns

We identify exons as alternative ‘cassettes’ using the following approach:

For each exon, we collect the following numbers

- A: Number of reads that support the exon with both splice sites
- B: Number of reads that support the left splice site with the right end of the read ending on the exon
- C: Number of reads that support the right splice site with the left end of the read ending on the exon
- D: Number of reads skipping the exon (but being assigned to the same gene)
- E: Number of reads overlapping the exon

First, we exclude exons, for which A+B+C+D<10. For these exons there are simply too few reads to perform any cluster-specific analysis.

Second, we exclude exons, for which (A+B+C+D)/(A+B+C+D+E)<0.8. These are exons, for which less than 80% of overlapping reads clearly support inclusion or exclusion of the exon. In these cases a binary view of ‘inclusion vs. exclusion’ is not advisable.

Third, we calculate a PSI for the left part of the exon (“leftPSI’)) as (A+B)/(A+B+D) and a ‘rightPSI’ as (A+C)/(A+C+D). We keep the exon if and only if, both leftPSI and rightPSI are contained in the interval [0.05,0.95].

For the exons remaining after this step, we proceed to calculating cluster specific (i.e., layer 4 in ‘old’ individuals) exon inclusion and skipping reads as well as a cluster specific PSI as (A+B+C)/(A+B+C+D) if and only if there A+B+C+D>=10. Exclusion and inclusion reads from the two groups (i.e., layer 4 ‘old’ and layer4 ‘young’) are then used to populate a 2×2 table. A table is kept for statistical testing if and only if three of the four cells in the 2×2 table have expected counts of 5 or higher.

The remaining exons (represented each by one table) are testing for a significant association of cell type and exon inclusion using two-sided fisher test. Calculated p-values are then corrected for multiple testing using the Benjamini-Yekutieli correction.

### Calling Differentially included Isoforms

After files containing all barcoded and UMI corrected reads with corresponding gene and transcript information (AllInfo files) were generated as described above, files were run through the IsoQuant() and DiffSplicingAnalysis() functions from the ScisorSeqR package (Joglekar, et al., Nature Communications, 2021). The IsoQuant function quantifies unique CAGE and PolyA sites, as well as whole isoforms, per gene and barcode. The DiffSplicingAnalysis() function identifies differential isoform usage by spatial region and age comparisons. Each isoform is assigned an ID based on the TSS, introns, and polyA-site information, with isoform 1 being most abundant per gene. Isoform counts were then identified by which comparison they originated from (ie. child v. young adult). Genes were filtered out if they did not reach the minNumReads criteria (set to 10). Remaining genes were tested in an maximum of 11×2 matrix of counts, where a maximum of 10 isoforms were identified, with all remaining isoform counts summed in row 11. P-values were calculated with a chi-squared test and a Benjamini Hochberg correction. Delta Pi values per gene were calculated as the sum of change in percent isoform from the top two isoforms detected.

### Calling Differentially included PolyA sites

We determined the PolyAs per read as assigned in the Allinfo file. Files were run through the DiffSplicingAnalysis() function where typeOfTest = ‘PolyA’, which identifies alternative PolyA sites by group. PolyA sites per gene were assigned an ID, with PolyA 1 being most abundant. Counts were identified per testing condition and summarized in an 11×2 table as described above. Testing was performed as described above for isoforms.

### Down sampling experiments

To compare two comparisons (ie. Cortical differences between age groups with WM differences across age groups) with equal power, we performed down sampling experiments. We first selected all exons which have at least 17 reads coming from 3 or more samples per age group. We then randomly selected 20 reads among the total. These reads were then used to re-calculate the difference in percent isoform inclusion between the ages (ΔPSI). Next, we selected 50 exons randomly for this comparison, enforcing that there be at most exon per gene. We then repeated these steps for all cell types which were compared. This yields 50 2×2 tables for all comparisons, with exactly equal column sums and the same characteristics (table number) for correction multiple testing. We then proceeded to carry out fisher tests and corrections for multiple testing and recorded the number of significant events for all comparisons. The procedure was repeated 100 times giving a distribution of significant percentages for both comparisons. These two distributions were compared with a Wilcoxon rank-sum test. For excitatory neuron sampling in Figure 6n, we employed cutoffs of 7 reads per sample in at least 3 samples per age group and sampled 15 exons and 30 genes due to smaller sample sizes.

### Identifying last exon length for PolyA differences

Reads which contained an annotated Poly(A) site were selected. The length of the last exon prior to the Poly(A) tail was calculated based on exon coordinates, respective to the strand direction. The average poly(A) length per gene was calculated per Cortex and WM age group. These lengths were plotted in boxplot format as seen in figure 4e. Additionally, we analyzed poly(A) length in genes which have differential poly(A) expression per group. Thus, from the cortical reads which contained an annotated Poly(A) site, we subsetted those which were mapped to the set of altered poly(A) genes in cortex. We similarly subsetted poly(A) reads from WM which were mapped to significantly altered genes in WM. We again calculated the average length of Poly(A) exons per gene for each area and age group which is shown as a boxplot in figure 4f.

### Repetitive Elements Analysis

The analysis concerning repeat identification and their occurrence was conducted using the RepeatMasker webserver with default parameters (A.F.A. Smit, R. Hubley, & P. Green, unpublished data; Current Version: open-4.0.9 with Dfam 3.0 only)..The analysis was carried out on upstream (150-30 upstream bp) and downstream (10-150 downstream bp) regions independently. To evaluate the top 4 types of repeats identified in the signal sequences, their occurrence in a significant group of exons (FDR < 0.05 + |DPSI| >0.5) was compared to that in background sequences (FDR > 0.05 + |DPSI| < 0.05) using Fisher’s exact test. The analysis considered both the individual contributions of each repeat class and their combined effect.

### Getting protein superfamily annotations per exon

Cortex tested exons were split into a highly variable group (FDR < .05 and |DPSI| > .5) and a background group (FDR >.05 and |DPSI| < .1). For each exon per group, we used the genomeToProtein() function of the ensembldb package and extracted the Ensembl ID, coordinates, and residue sequence of the protein identified. We filtered the obtained protein identifiers based on their corresponding Ensembl transcript IDs and limited the search to the principal isoforms from the APPRIS database^10^. For each protein sequence, we ran the SUPERFAMILY^11,12^ tool that uses the hidden Markov model to identify the structural-defined SCOP protein domain families and the domain boundaries. The tool was implemented in InterProScan^13–15^. Protein regions not associated with domains were considered interdomain linkers. Only transcripts with domains which spanned exons in background and significant were selected.

### Modeling Domain Architecture

Domain architecture of each protein isoform were obtained using structure-driven SUPERFAMILY domain annotation^12^ extracted from Ensemble^16^. Structural characterizations of the proteins were done *de novo* using AlphaFold 3^17^ server no structural templates were detected for either protein, preventing a potentially more accurate template-based modeling approach. Some of the long interdomain linkers in the proteins were modeled with low or very low confidence scores (70 > plDDT > 50 and plDDT < 50), correspondingly and were excluded from the final models.

### Ribosplitter Protein Domain Analysis

Spladder^18^ was performed with .bam files with isolated L4 reads per individual and basic inputs as described in the online tutorial (https://github.com/ratschlab/spladder?tab=readme-ov-file), using commands *spladder build* and *spladder test*. The output folder created from *spladder test* was used as the input for RiboSplitter^19^, using functions read_details(), read_isoforms() with basic inputs as documented in the online tutorial (https://github.com/R-Najjar/RiboSplitter). Basic tutorial commands were followed in order to generate a table containing differentially expressed events between age groups. Examples were plotted with commands *splicing_figure*() and *domain_fig*() using basic inputs.

## Supplemental Figures

**Figure S1.**
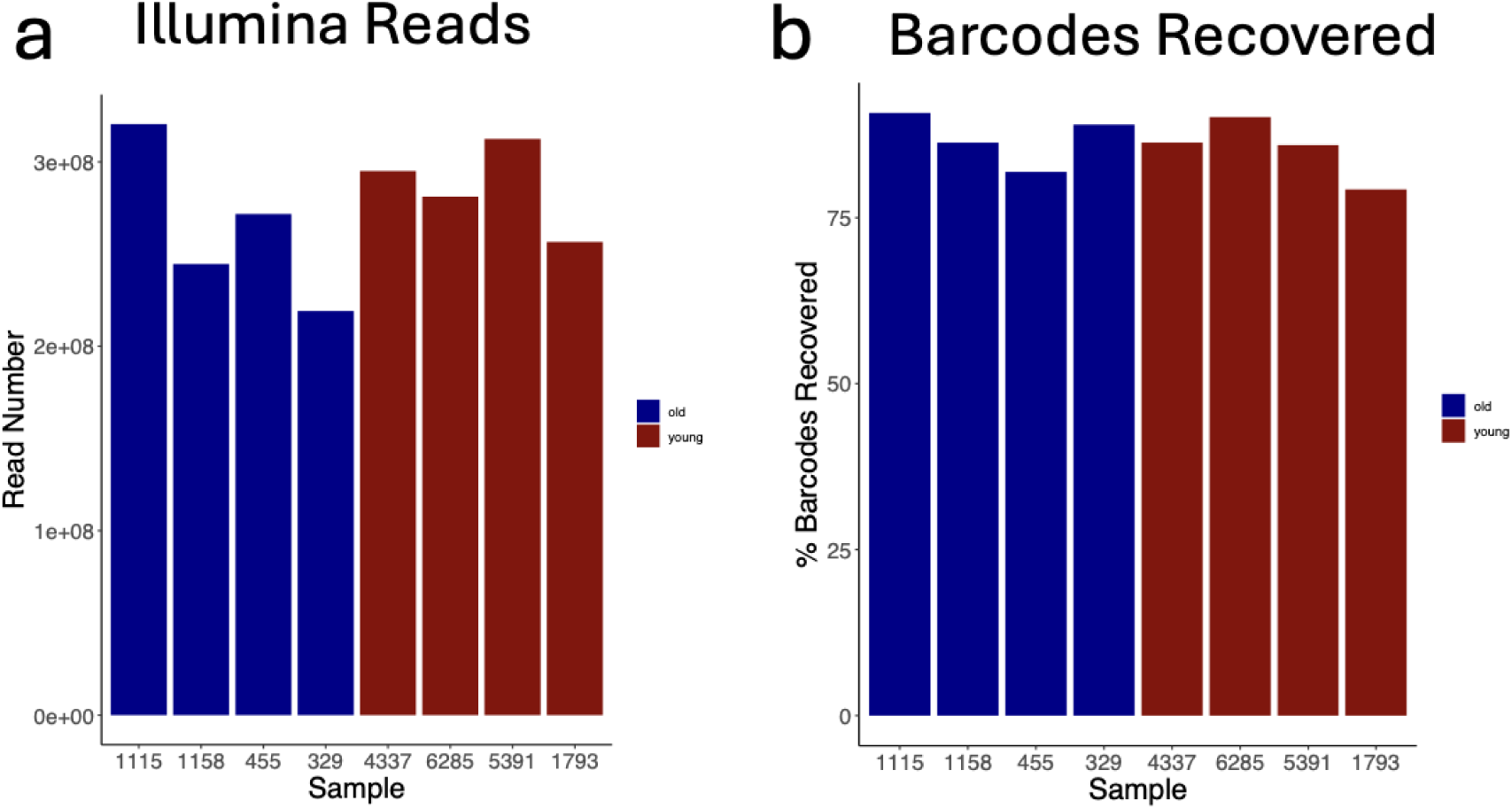
**a)** Number of sequenced Illumina reads by sample. **b)** Percent of barcodes recovered from Illumina data.

**Figure S2.**
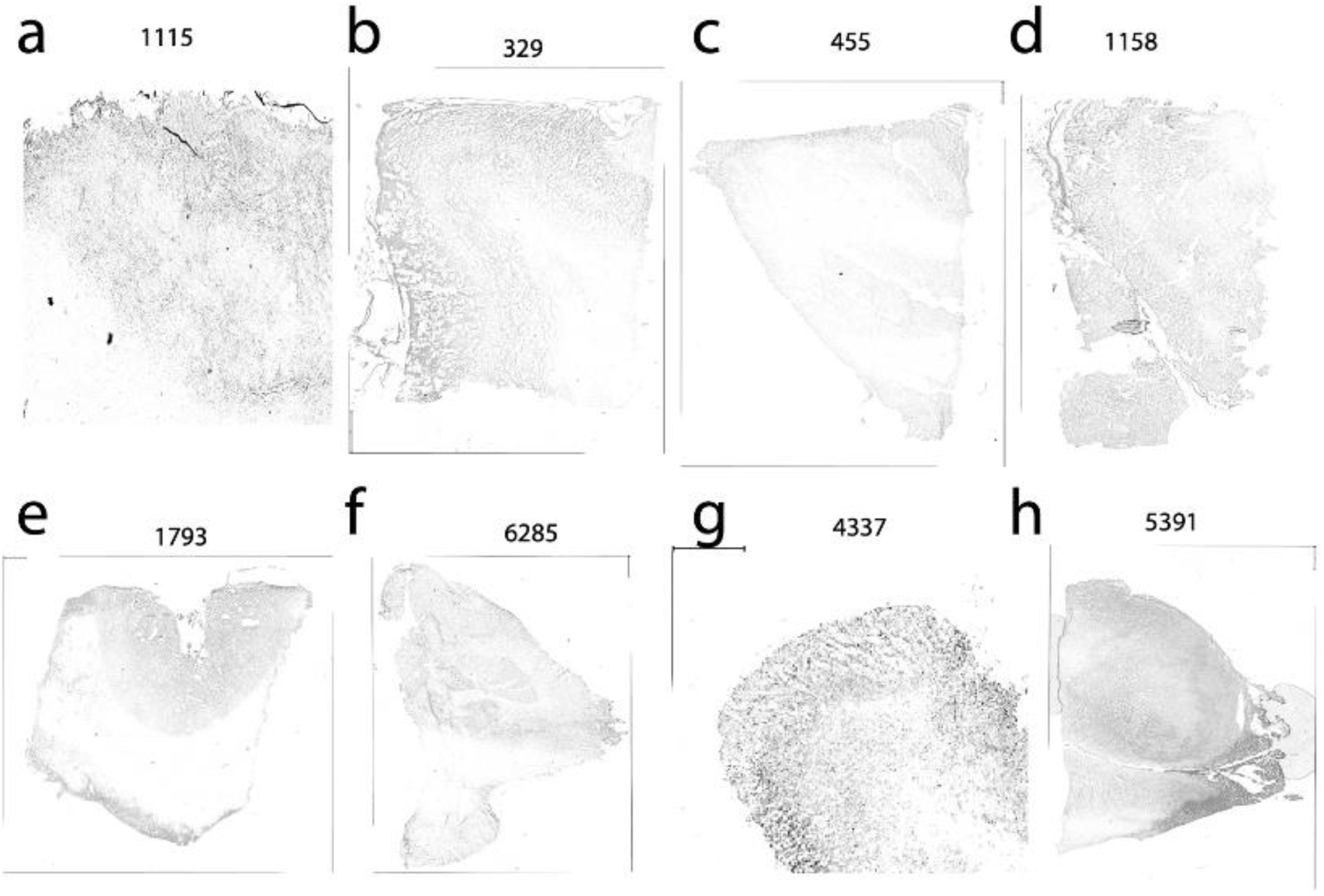
**a-h)** Hematoxylin and Eosin staining of 10 uM tissue slice following experimental slice for each sample. Panels a-d are in the young adult group. Panels e-h are in the child group.

**Figure S3:**
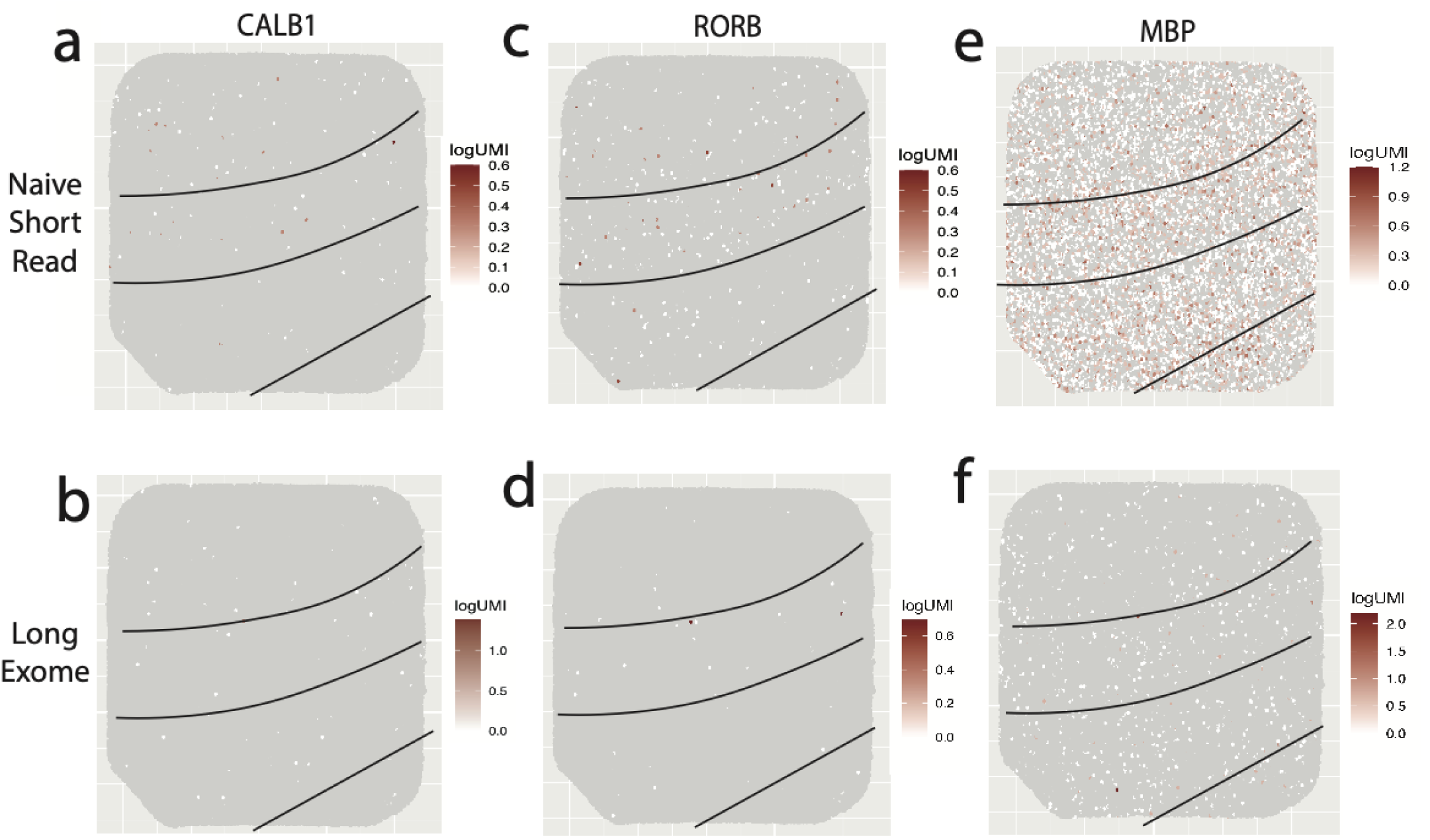
**a)** Log10(UMI) counts of Naïve Short-Read data per spot of gene CALB1 plotted by spatial location. **b)** Log10(UMI) counts of Long-Exome LR data per spot of gene CALB1 plotted by spatial location. **c)** Log10(UMI) counts of Naïve Short-Read data per spot of gene RORB plotted by spatial location. **d)** Log10(UMI) counts of Long-Exome LR data per spot of gene RORB plotted by spatial location. **e)** Log10(UMI) counts of Naïve Short-Read data per spot of gene MBP plotted by spatial location. **f)** Log10(UMI) counts of Long-Exome LR data per spot of gene MBP plotted by spatial location.

**Figure S4.**
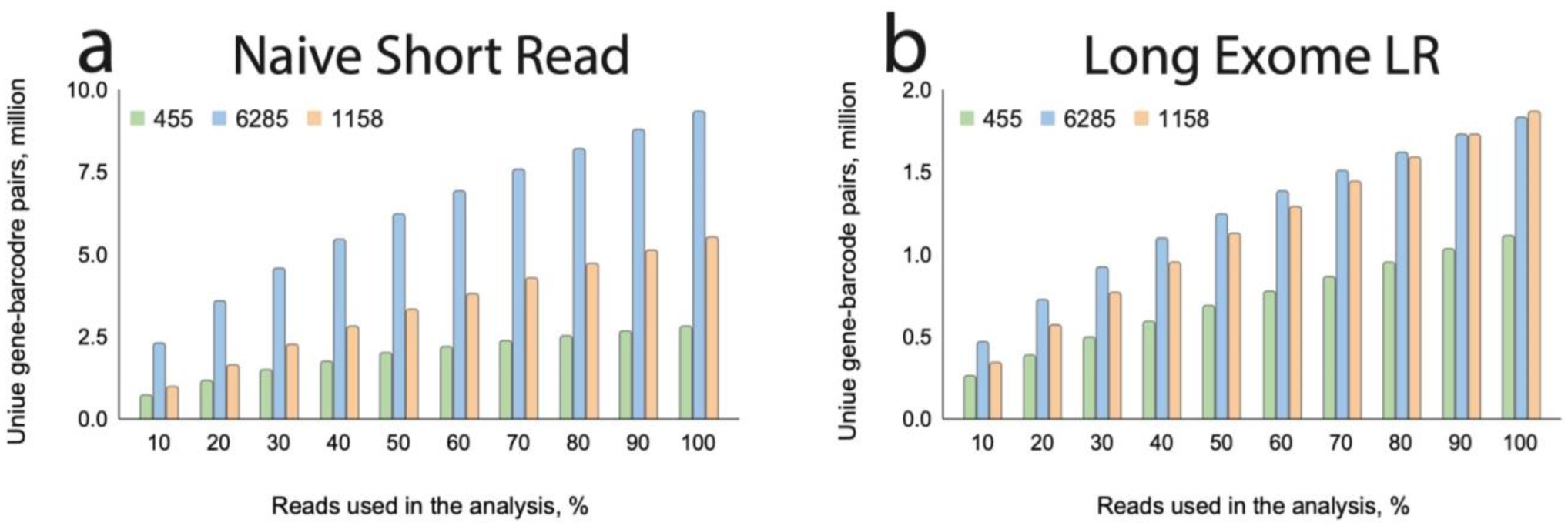
**a)** Unique gene-barcode pairs in Naive SR data plotted in bins per fraction of data processed. **b)** Unique gene-barcode pairs in Long Exome LR data plotted in bins per fraction of data processed. Color indicates 3 individual samples.

**Figure S5:**
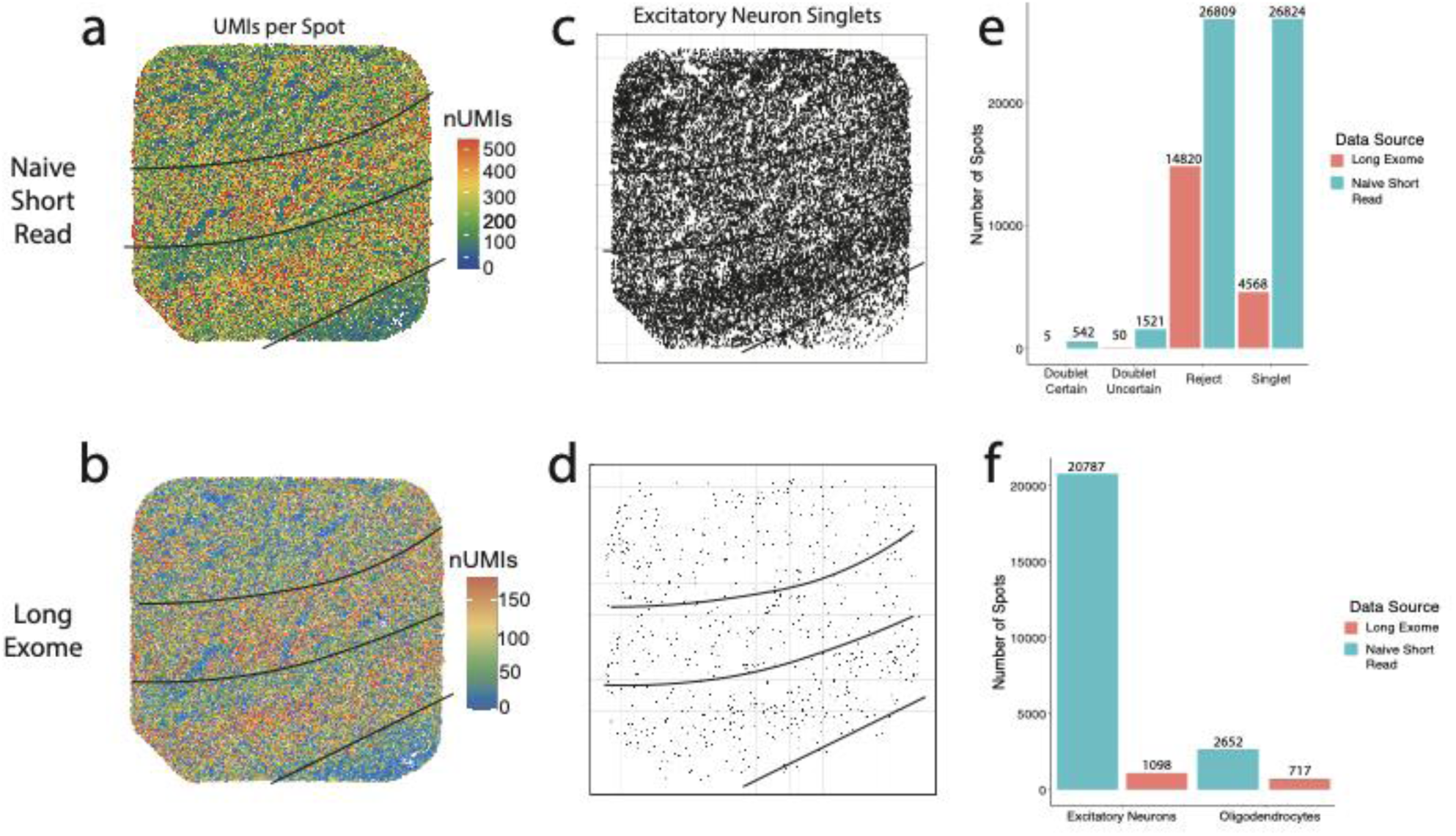
**a)** UMIs per spot from Naive short-read data plotted by spatial location. **b)** UMIs per spot from Long-Exome data plotted by spatial location. **c)** Excitatory-neuron singlets and their positions identified from inputting Naive Short-Read data into a deconvolution program. **d)** Excitatory-neuron singlets and their positions identified from inputting Long-Exome data into a deconvolution program. **e)** Number of identified “Doublet Certain”, “Doublet Uncertain”, “Reject”, or “Singlet” spots by data input. **f)** Number of identified excitatory-neuron and oligodendrocyte singlets by data input.

**Figure S6.**
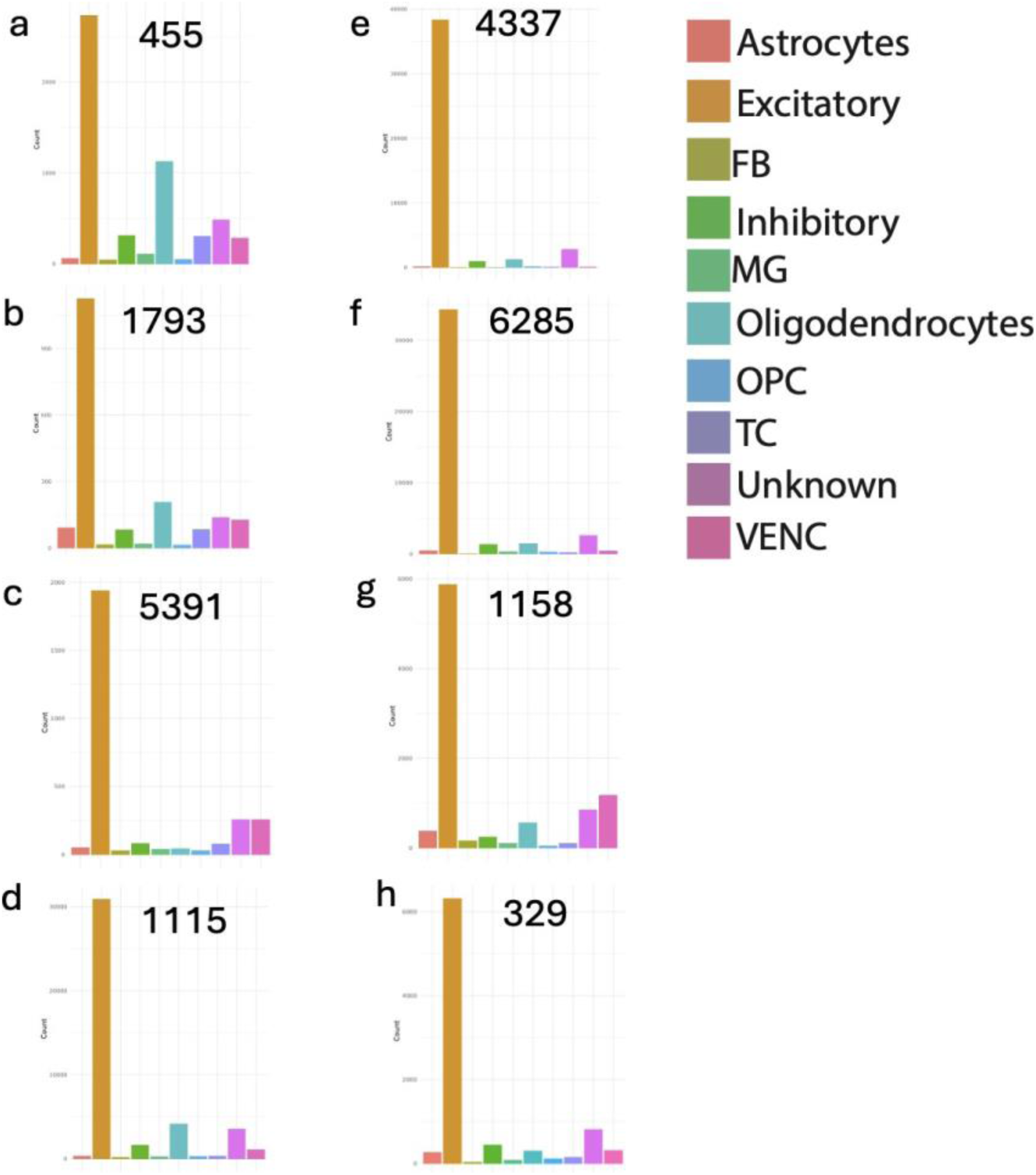
**a-h)** Robust Cell Type Decomposition (RCTD) program results showing total number of singlet spots per cell type by sample, including astrocytes, excitatory neurons, fibroblasts (FB), inhibitory neurons, microglia (MG), oligodendrocytes, oligodendrocyte precursor cells (OPC), T cells (TC), unknown, and vascular endothelial cells (VENC). Young Adult age group are panels a,d,g, and h. Child age group are panels b,c, e and f.

**Figure S7.**
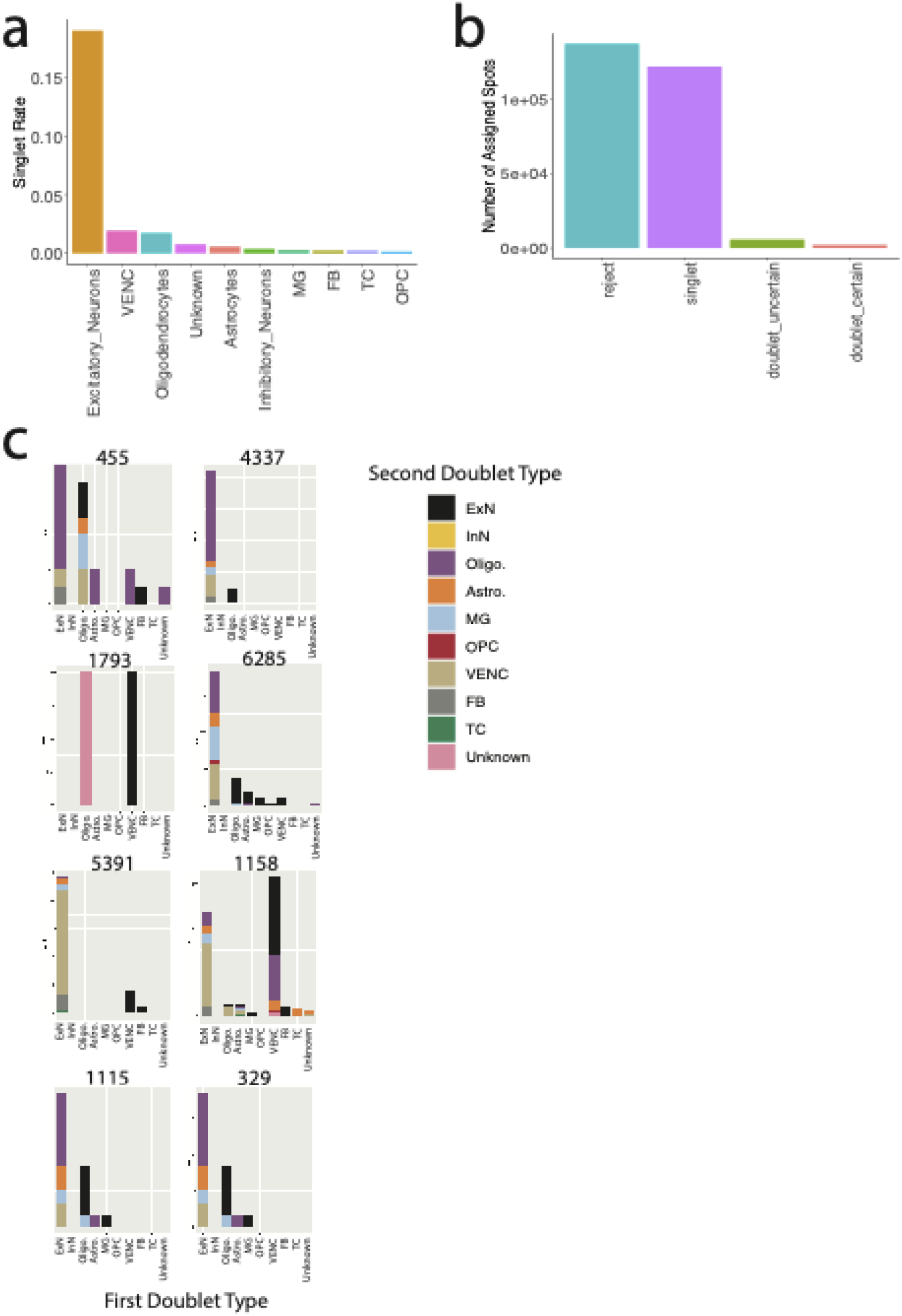
**a)** Number of assigned “singlets” from RCTD from all samples combined per cell type divided by the total number of detected barcodes in Illumina data (n=482,777). **b)** Number of assigned spot classes from all samples combined. **c)** Stacked bar plot by sample denoting the cell-type compositions of certain doublets. X axis lists the first identified cell type, and stacked color indicates the second cell identified type. Cell types include ExN=Excitatory Neurons, InN=Inhibitory Neurons, Oligo.= Oligodendrocytes, Astro.=Astrocytes, MG=Microglia, OPC=Oligodendrocyte Precursor Cells, VENC=Vascular Endothelial, FB=Fibroblasts, TC=T Cells, and Unknown.

**Figure S8.**
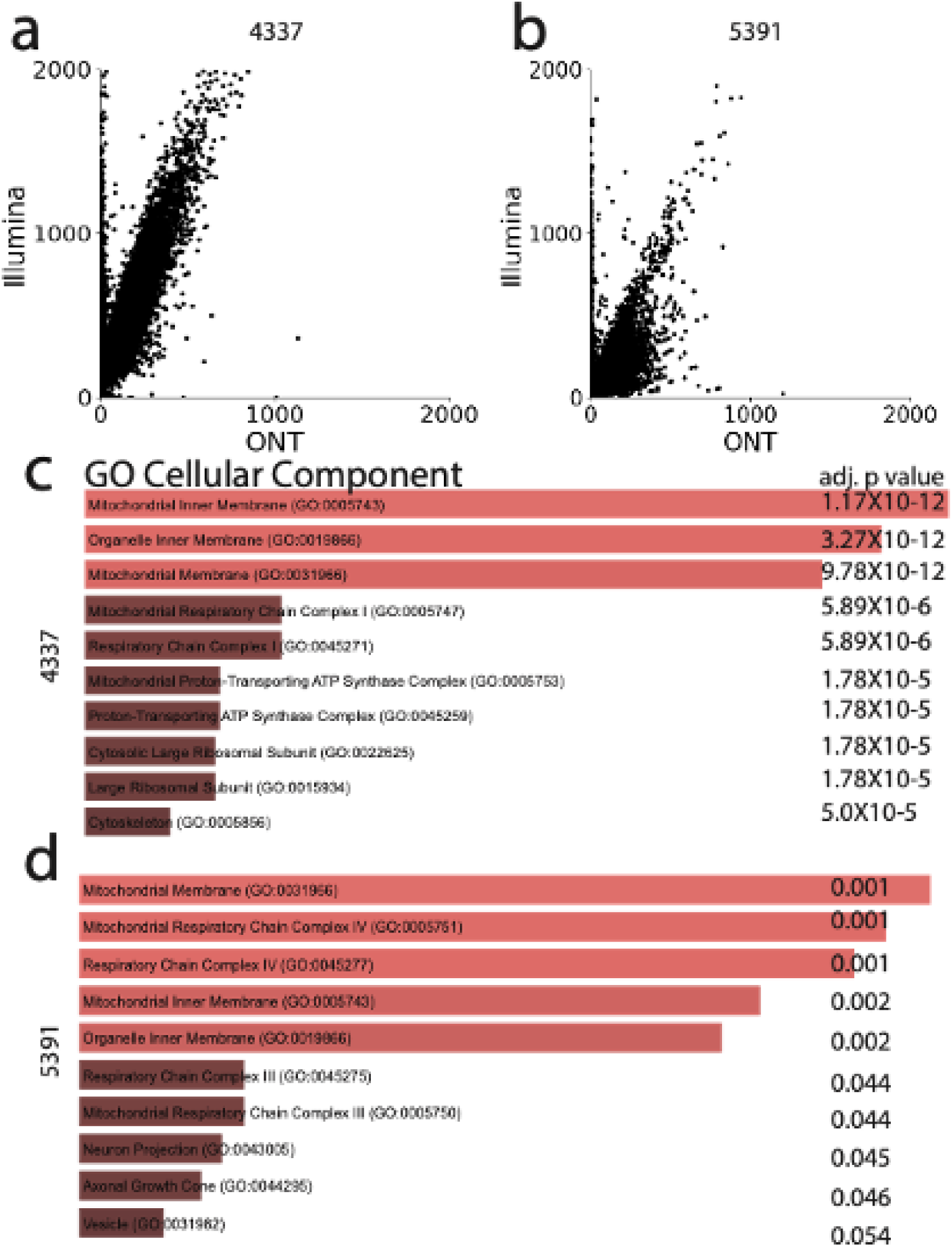
**a)** Dot plot of number of UMIs per barcode from a Long Exome dataset and Illumina dataset from the sample 4337. **b)** Dot plot of number of UMIs per barcode from a Long Exome ONT dataset and Illumina dataset from the sample 5391. **c)** EnrichR Gene Ontology analysis of the top 100 genes differentially regulated between barcodes which have high UMIs in Illumina data and low UMIs in ONT data for the sample 4337. **d)** EnrichR Gene Ontology analysis of the top 100 genes differentially regulated between barcodes which have high UMIs in Illumina data and low UMIs in ONT data for the sample 5391.

**Figure S9.**
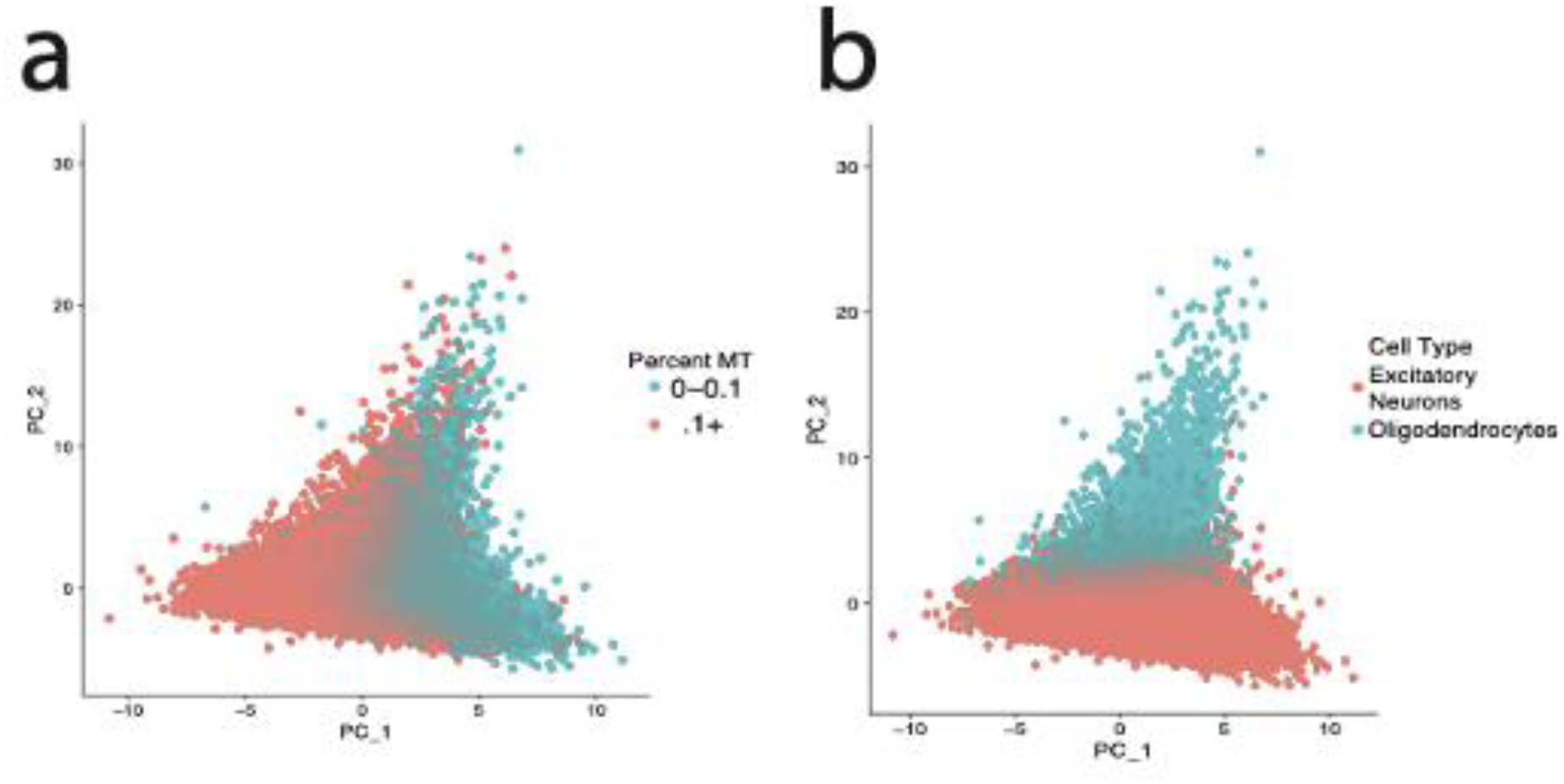
**a)** Spatial barcodes of singlets plotted by PC 1 and PC2. Color indicates percent of reads coming from mitochondrial (MT) genes. Blue=Percent MT from 0 - 0.1; Red= Percent MT 0.1+. b) Spatial barcodes of singlets plotted by PC 1 and PC2. Color indicates RCTD determined singlet cell type. Blue=Oligodendrocytes; Red= Excitatory Neurons.

**Figure S10.**
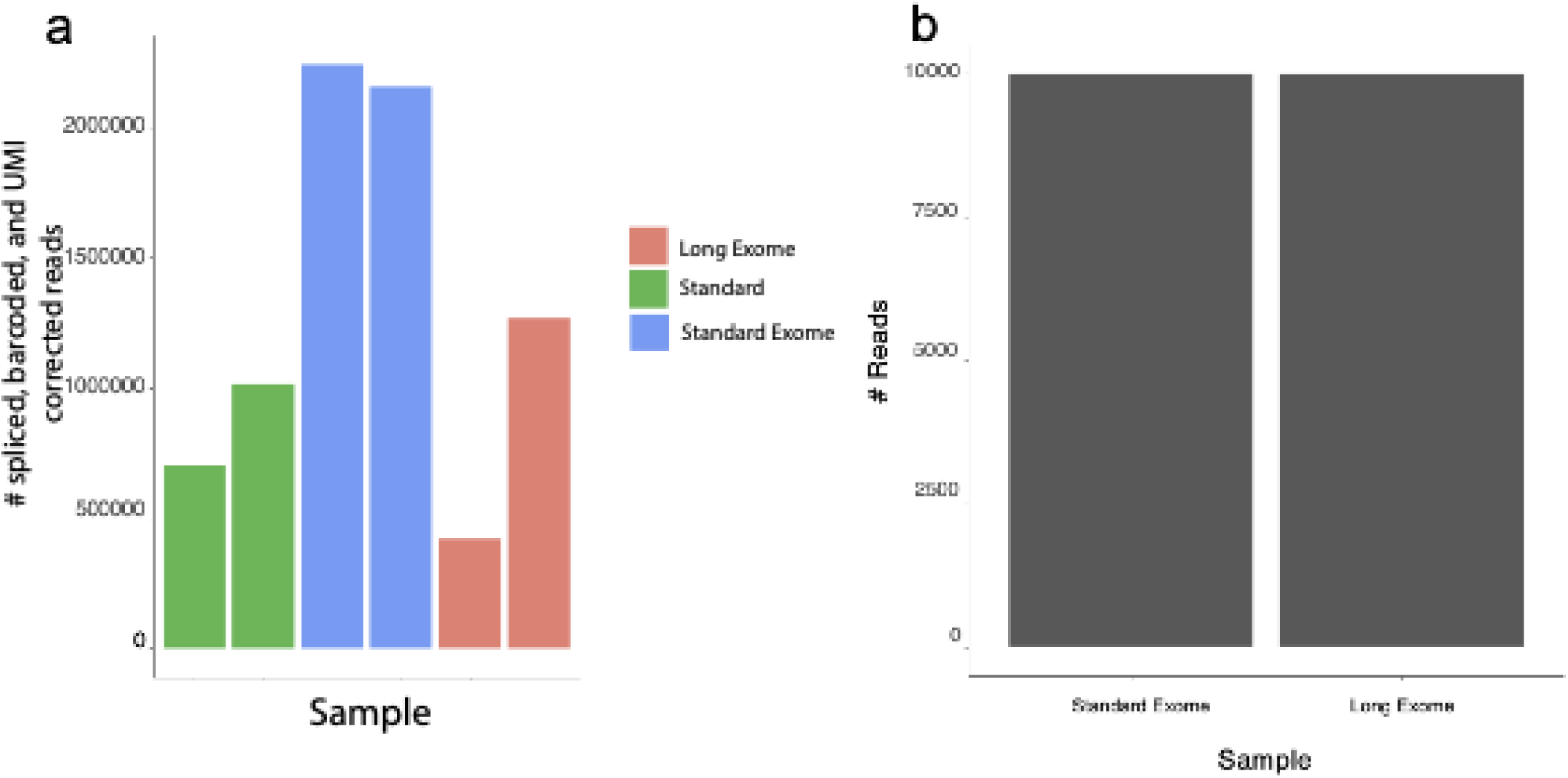
**a)** Number of reads from the 2 spatial samples shown in figure 2b and 2c. Standard: Untagmented cDNA acquired from the curio pipeline; Standard Exome: Untagmented cDNA acquired from the curio pipeline followed by an enrichment of molecules containing exons; Long Exome: Untagmented cDNA acquired from the curio pipeline followed by an enrichment of molecules containing exons and are optimally size selected. **b)** Number of reads from each single cell sample shown in figure 2d-f. Standard Exome: Untagmented cDNA acquired from 10X Genomics’ single cell pipeline followed by an enrichment of molecules containing exons; Long Exome: Untagmented cDNA acquired from 10X Genomics’ single cell pipeline followed by an enrichment of molecules containing exons and are optimally size selected.

**Figure S11.**
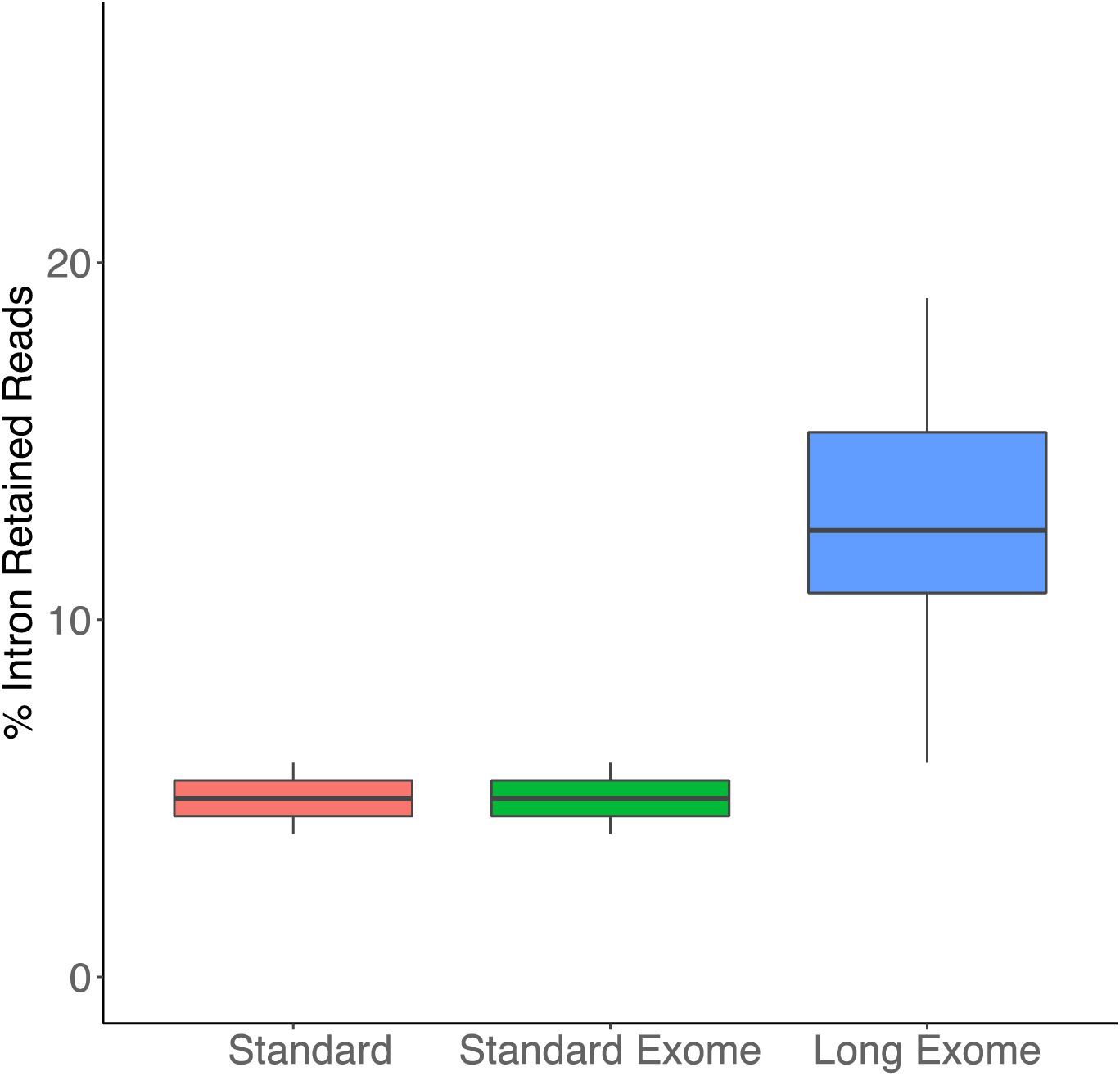
Percent of all reads which contain retained introns across long read sequencing datasets.

**Figure S12.**
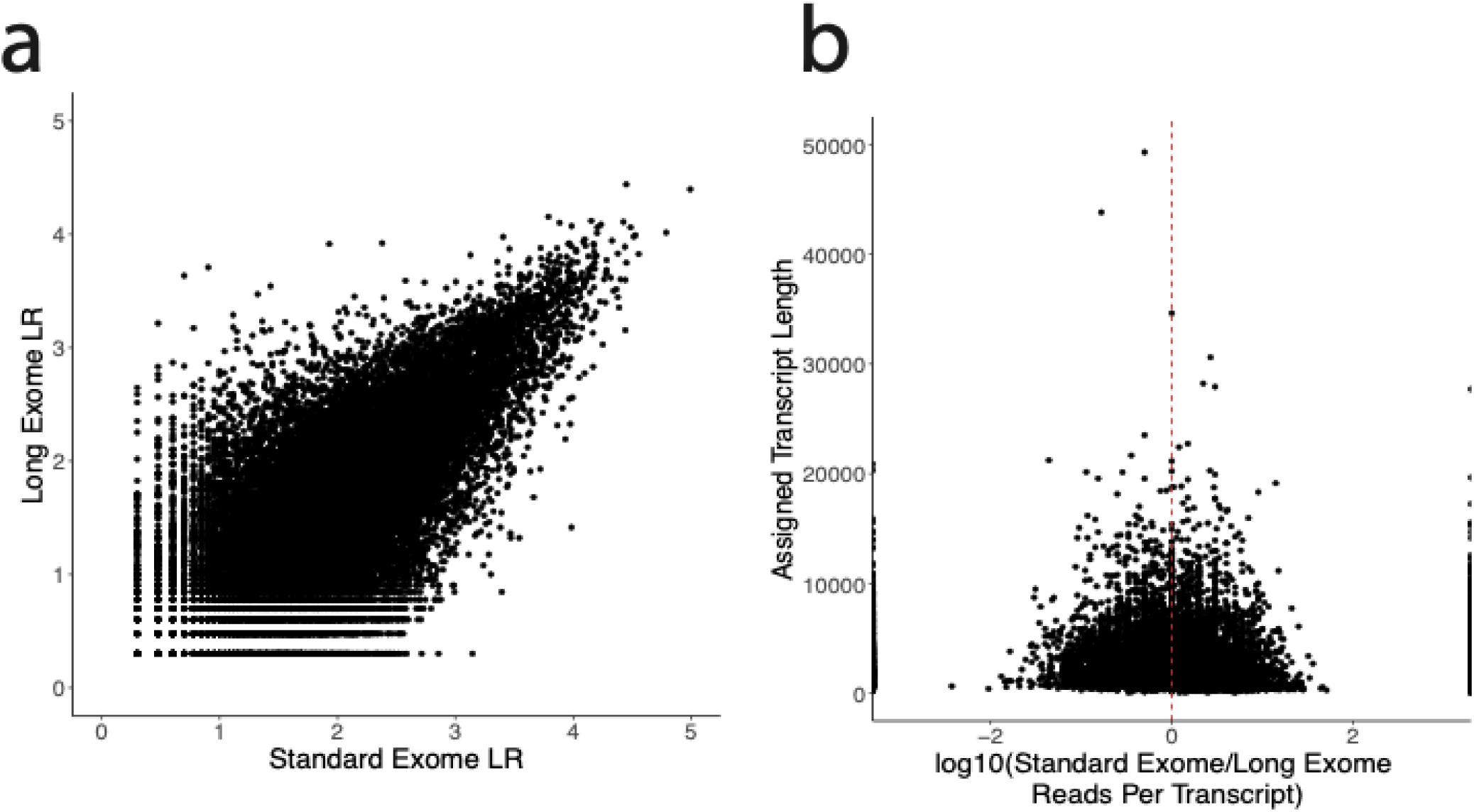
**a)** Correlation of transcript expression between equally downsampled Standard Exome LR (n=2) and Long Exome LR datasets (n=2). **b)** Log10 of ratio reads per transcript (Standard Exome LR/Long Exome LR) by assigned transcript length. Red line indicates a value of 0.

**Figure S13.**
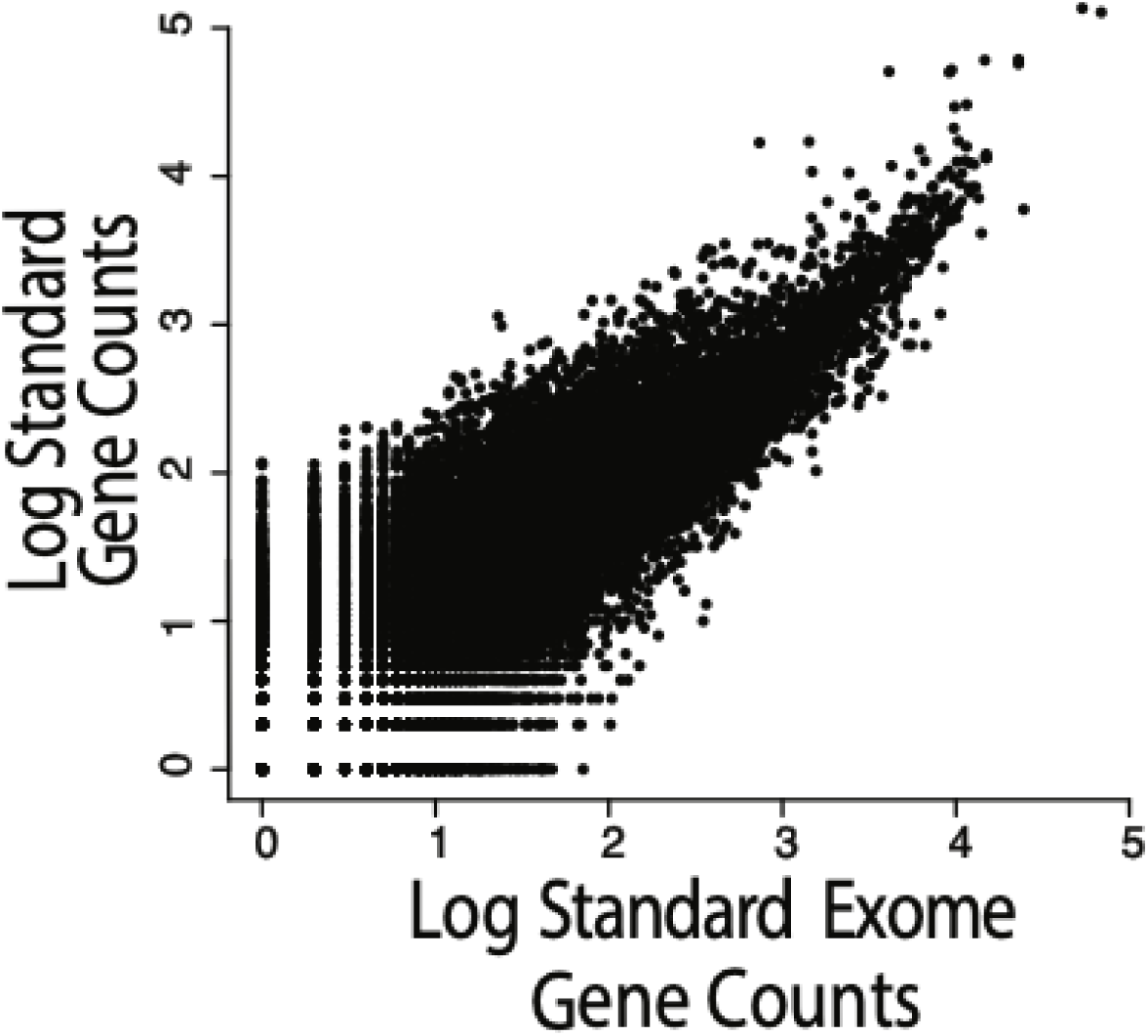
Log10 of the number of reads per gene between Standard Exome and Control No Exome datasets.

**Figure S14.**
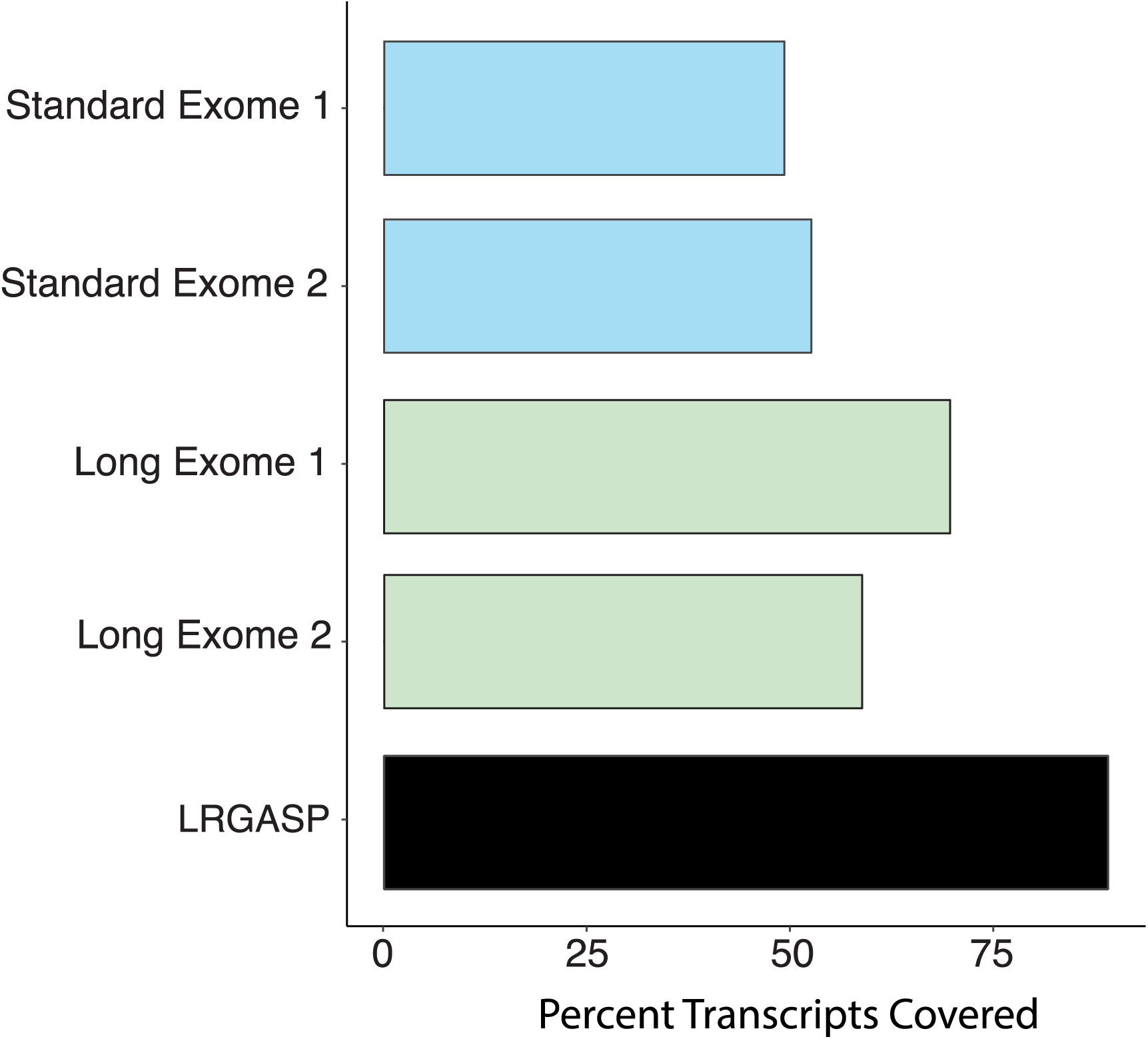
Percent of the transcript per base which is spanned by reads in each dataset. Dataset 1=455; Dataset 2=5391; LRGASP data is bulk, full length RNA seq data from the LRGASP consortium.

**Figure S15.**
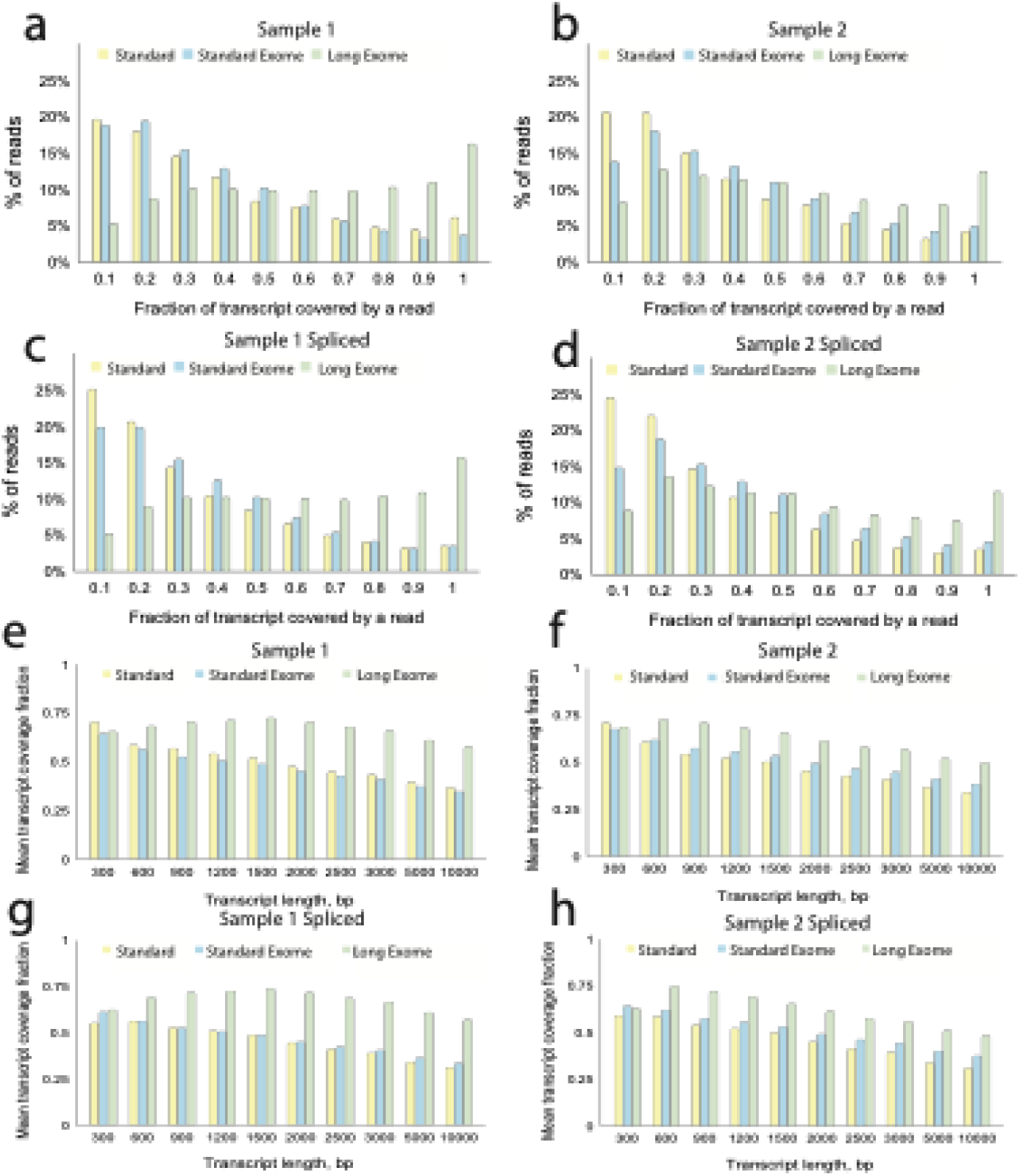
**a)** % of reads which fall into bins of fraction of transcript covered in sample 1 across standard, standard exome, and long exome datasets. **b)** Same as (a) but for sample 2 across standard, standard exome, and long exome datasets. **c)** % of spliced reads which fall into bins of fraction of transcript covered in sample 1 across standard, standard exome, and long exome datasets. **d)** Same as (c) but for sample 2 across standard, standard exome, and long exome datasets. **e)** Average fraction of transcript covered binned by reference transcript length covered in sample 1 across standard, standard exome, and long exome datasets. **f)** Same as (e) but for sample 2 across standard, standard exome, and long exome datasets. **g)** Fraction of average spliced transcript covered binned by transcript length covered in sample 1 across standard, standard exome, and long exome datasets**. h)** Same as (g) but for sample 2 across standard, standard exome, and long exome datasets. Sample 1 indicates sample 455 and sample 2 indicates sample 5391.

**Figure S16.**
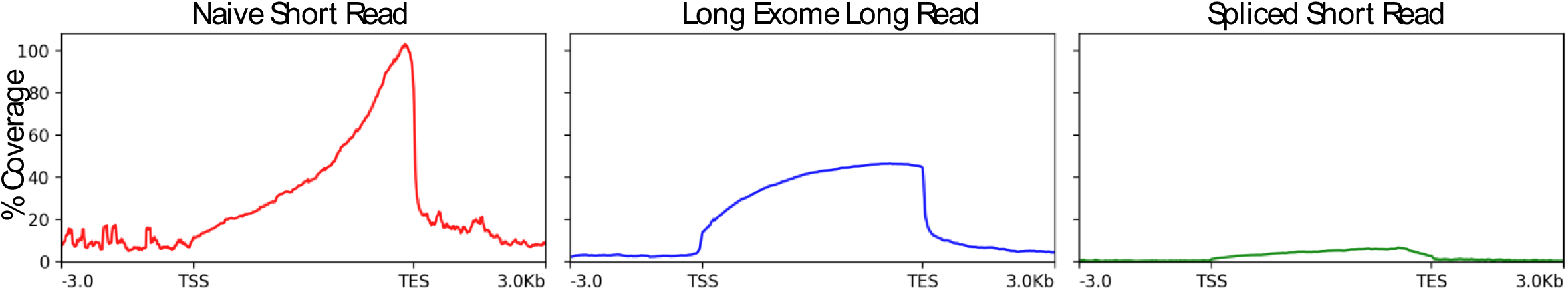
Metagene plot showing exonic coverage of annotated genes with normalized length. Red shows coverage from the Naïve Short Read data, blue trace shows coverage from the Long Exome LR data, and green trace shows coverage from the Spliced Short read data. TSS indicates transcription start site and TES indicates transcription end site.

**Figure S17.**
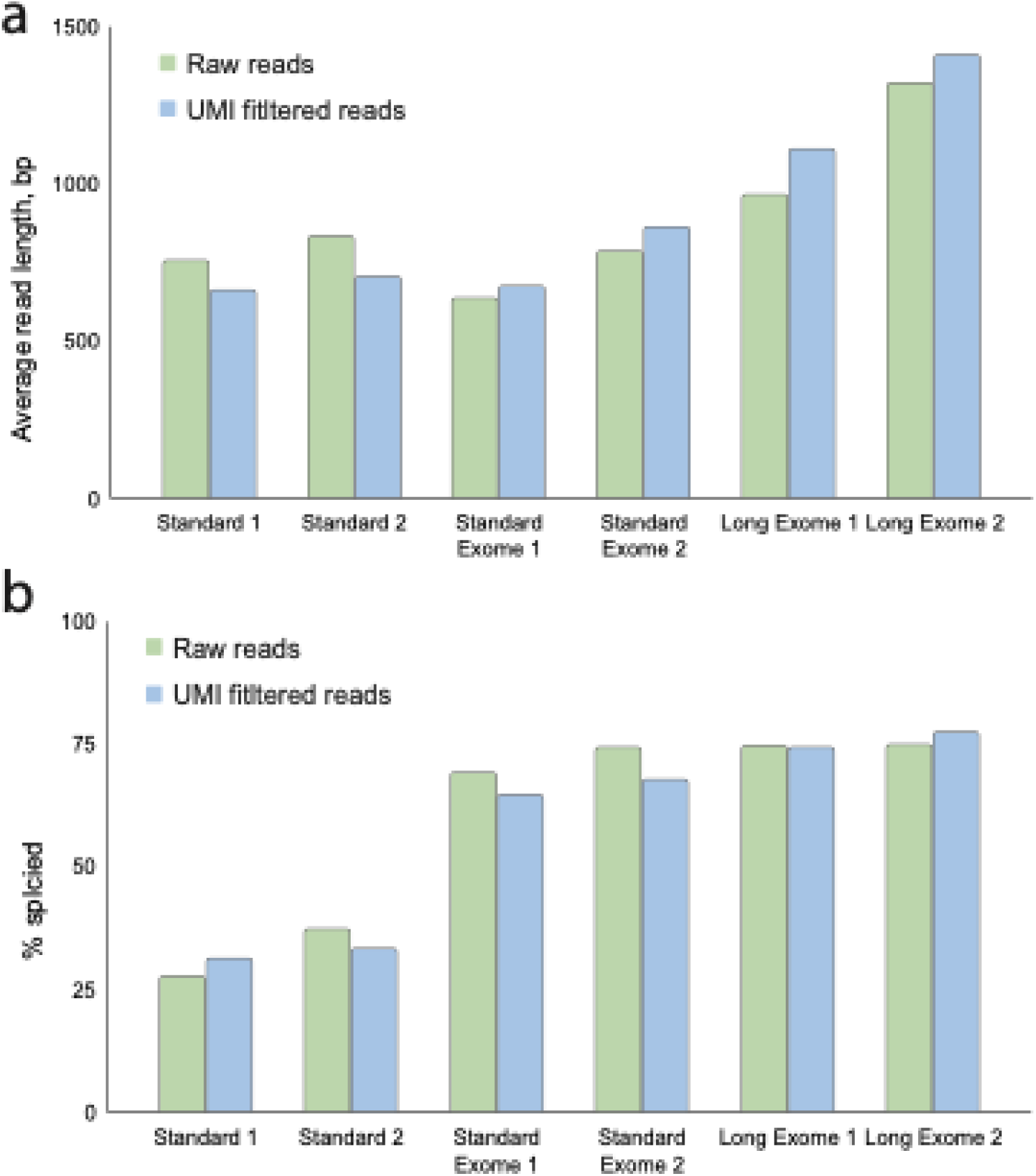
**a)** Average read length per sample separated by all reads (raw) and UMI filtered. **b)** % of spliced reads per sample separated by all reads (raw) and UMI filtered.

**Figure S18.**
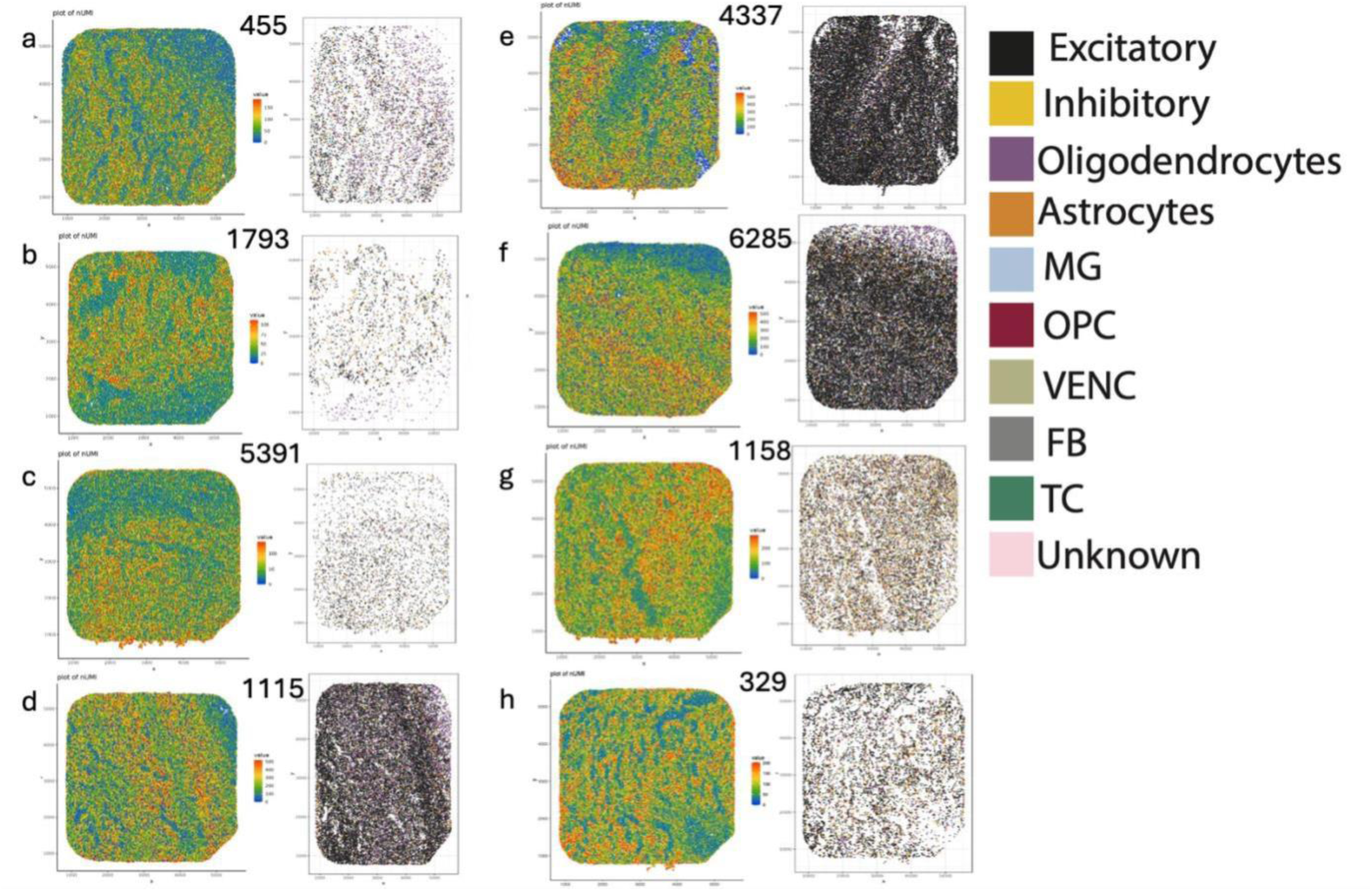
**a-h)** Left panel: Number of UMIs per spot plotted by spatial location for each sample. Right panel: RCTD predicted singlets plotted by cell type and spatial location for each sample, including astrocytes, excitatory neurons, fibroblasts (FB), inhibitory neurons, microglia (MG), oligodendrocytes, oligodendrocyte precursor cells (OPC), T cells (TC), unknown, and vascular endothelial cells (VENC).

**Figure S19.**
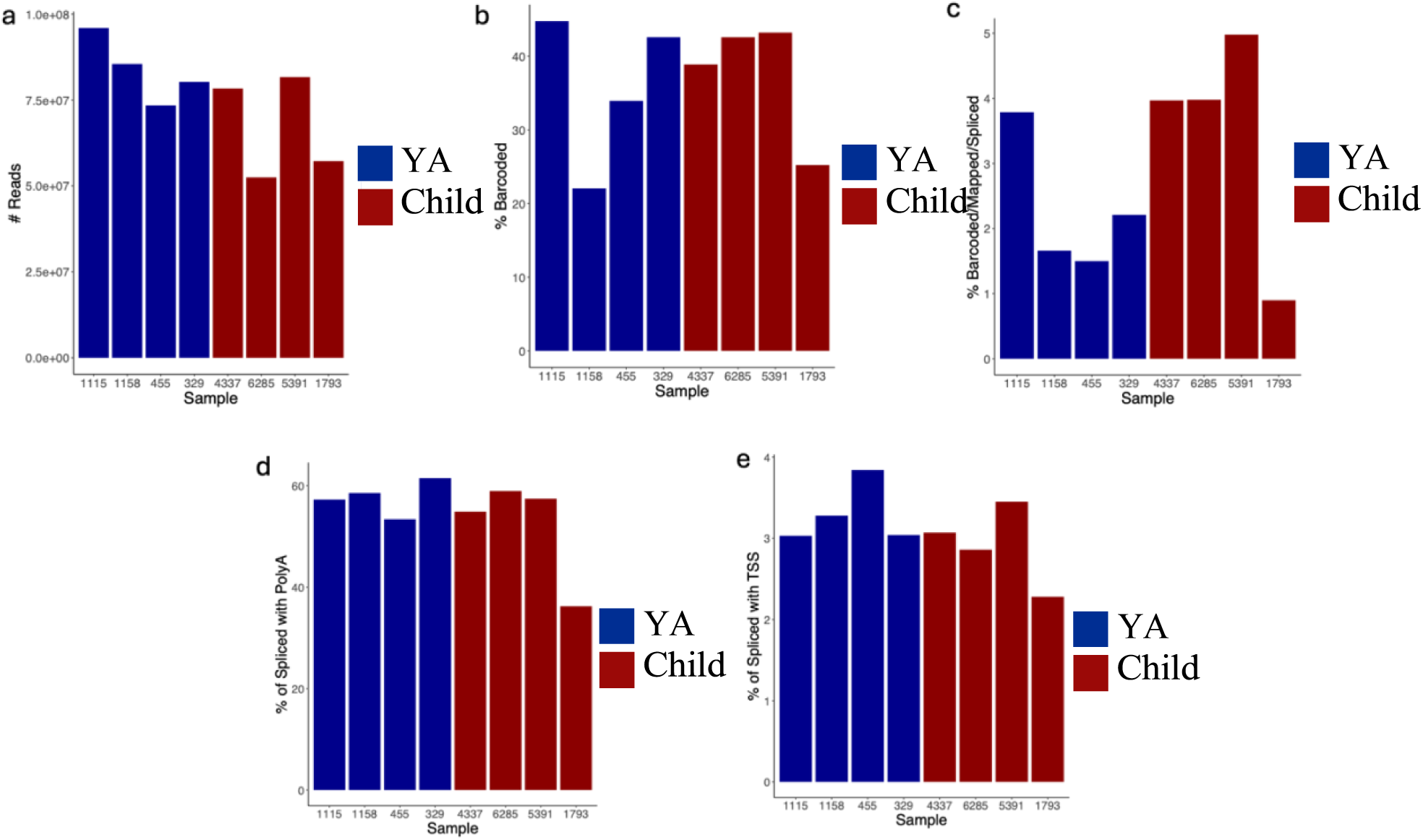
**a)** Number of Oxford Nanopore Technology (ONT) reads sequenced per sample. **b)** Percent of ONT reads containing proper barcode with a barcode score of at least 13 per sample. **c)** Percent of ONT reads which are barcoded (score >=13), mapped to hg38, and spliced as defined by Spl-isoquant. **d)** Percent of the barcoded, mapped, and spliced reads with a defined PolyA site. **e)** Percent of the barcoded, mapped, and spliced reads with a defined TSS site.

**Figure S20.**
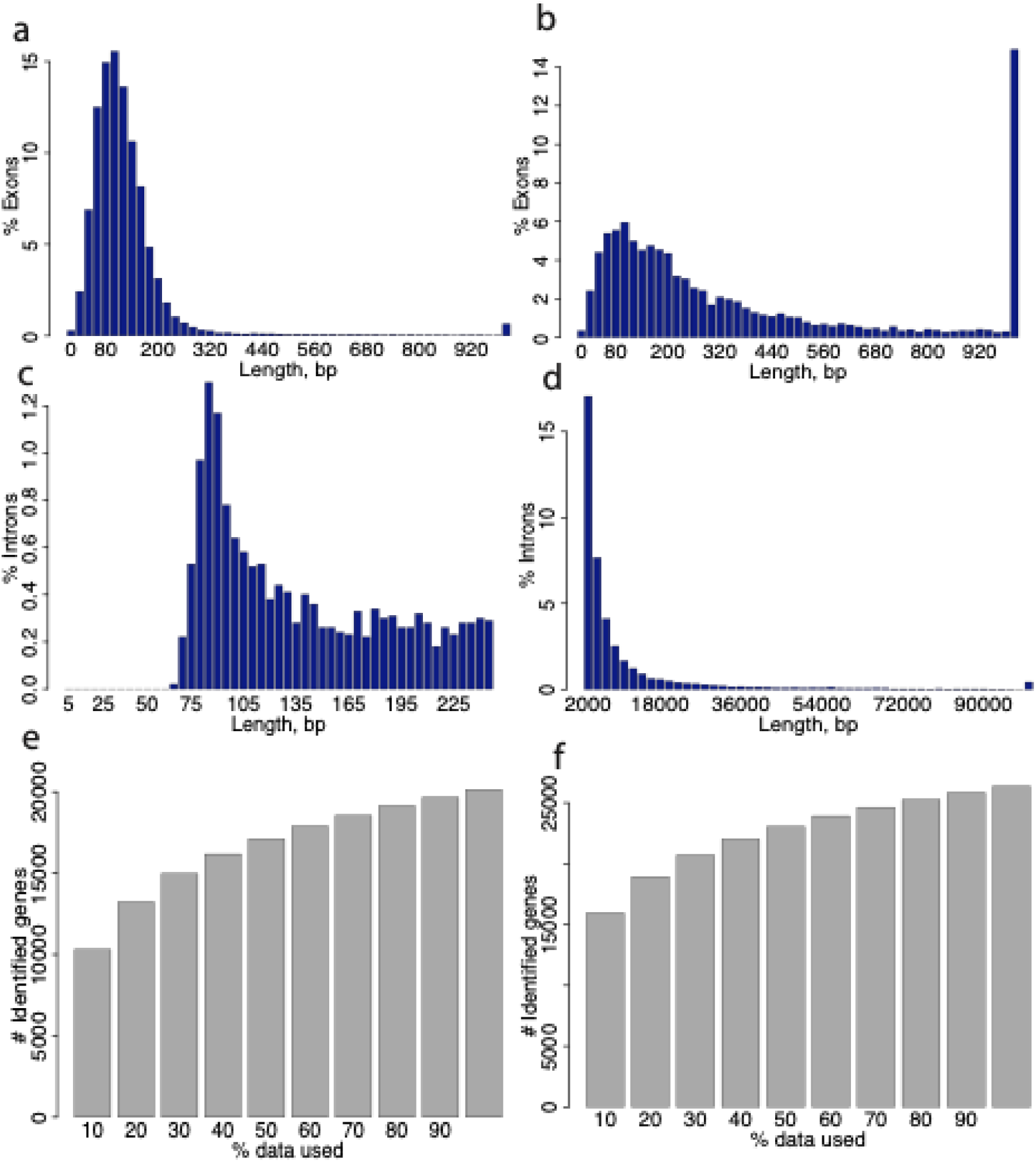
**a)** Length distribution of internal exons where last bar corresponds to all exons >= 1000 bp. **b)** Length distribution of terminal exons where last bar corresponds to all exons >= 1000 bp. **c)** Intron length distribution between sizes 0-250 bp. **d)** Intron length distribution where last bar corresponds to all exons >= 100k bp. **e)** Number of identified genes with >=10 UMIs when sampling decreasing percentages of spliced data. **f)** Number of identified genes with >=3 UMIs when sampling decreasing percentages of spliced data.

**Figure S21.**
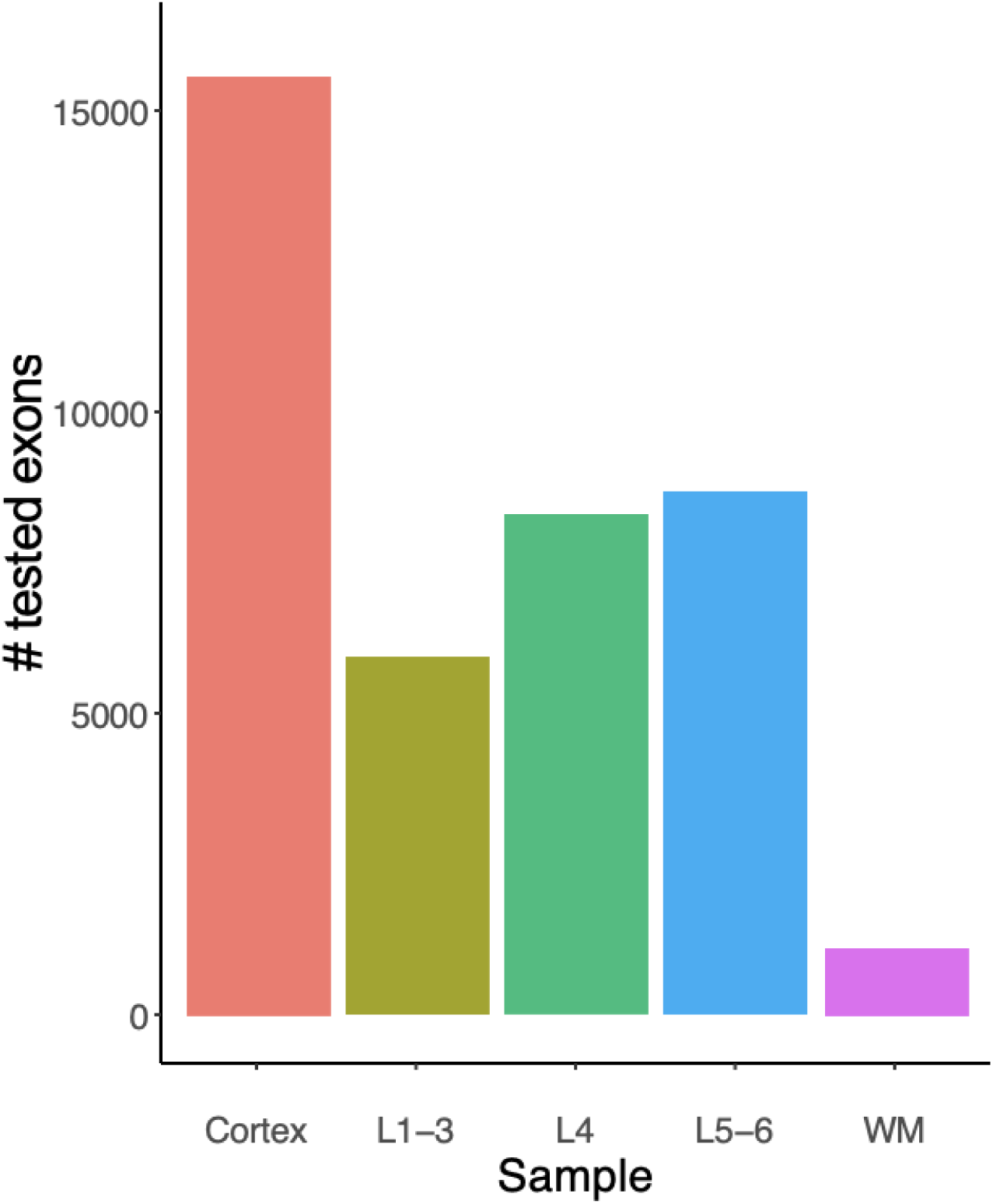
Number of tested exons for each area-specific comparison by age. Areas include Cortex, Layers 1-3 (L1-3), Layer 4 (L4), Layers 5-6 (L5-6), and white matter (WM).

**Figure S22.**
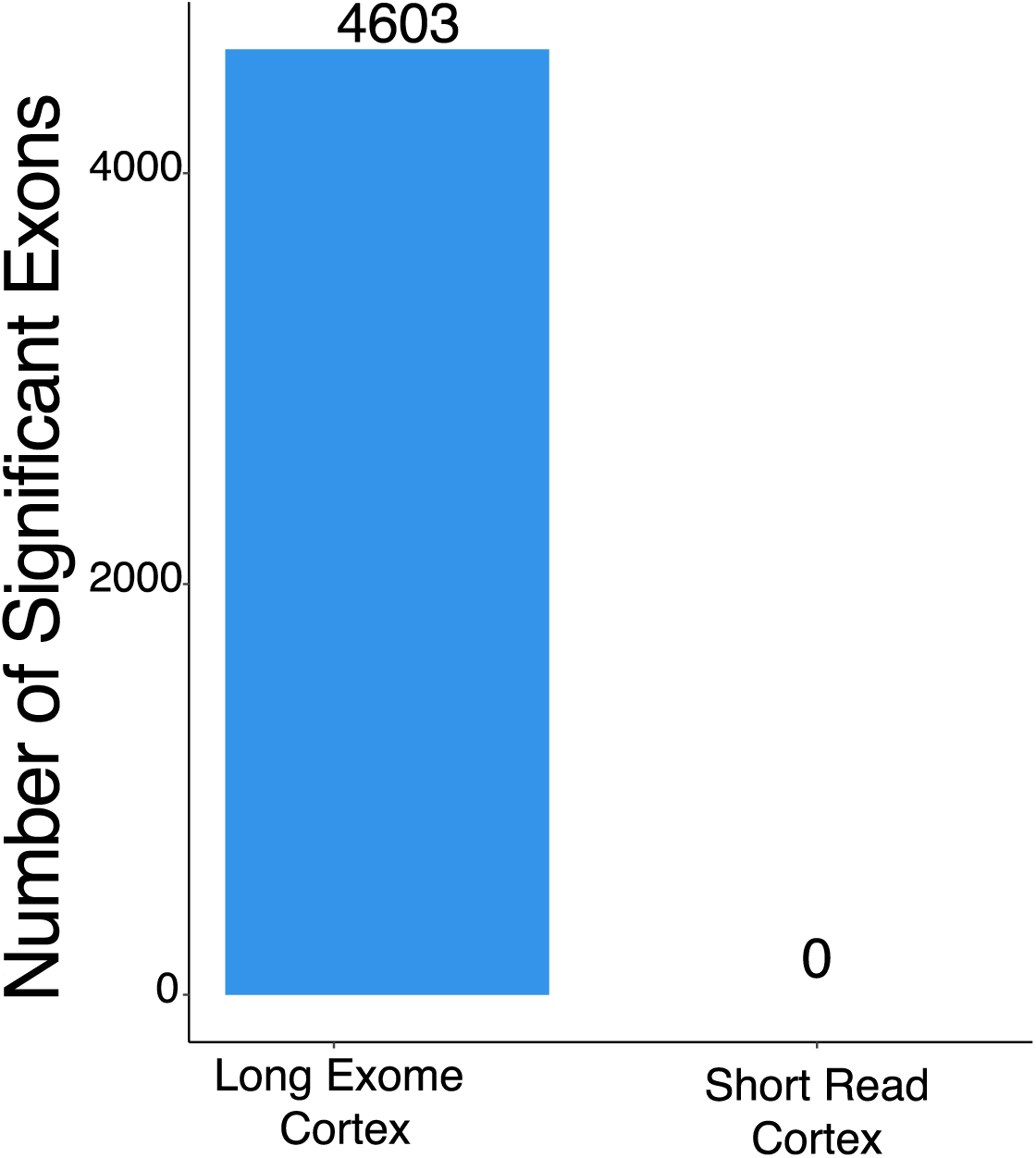
Number of significant exons identified to be regulated across age in the cortex by dataset. Long Exome Cortex: ONT data which is exome enriched and long-molecule selected. Short Read: Spliced short reads from the Naïve short read dataset.

**Figure S23.**
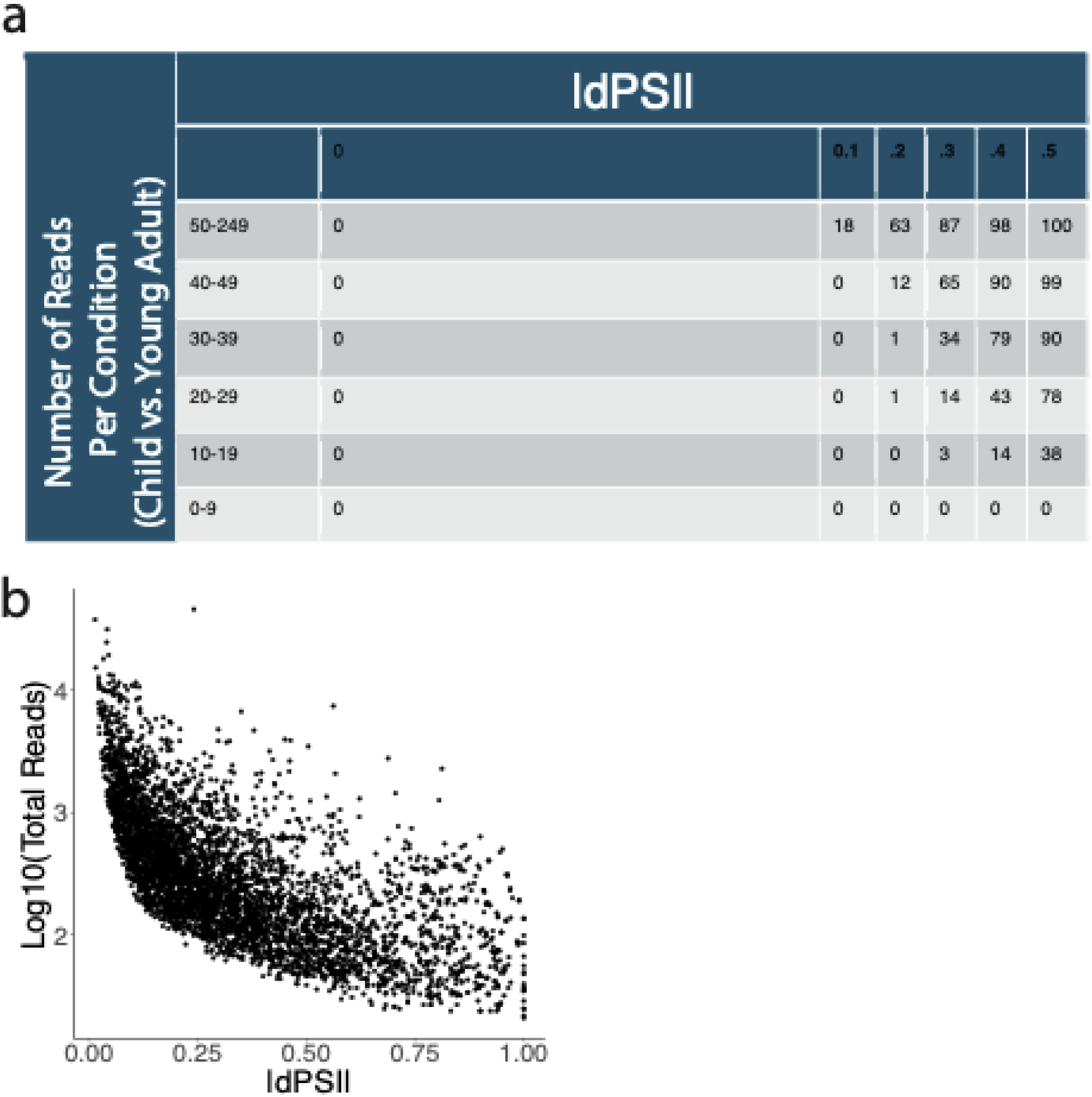
a) Matrices with 1000 reads per condition with predefined |dPSI| were downsampled to read counts of [0,9], [10,19], [20,29], [30,39], [40,49], [50,249] per condition. Values indicate the fraction of matrices that pass Benjamini-Yekutieli correction for multiple testing per |deltaPSI| and read number combination. b) Significant exons from the Cortex Child vs. Young Adult comparison plotted by |dPSI| and Log10(Total Reads).

**Figure S24.**
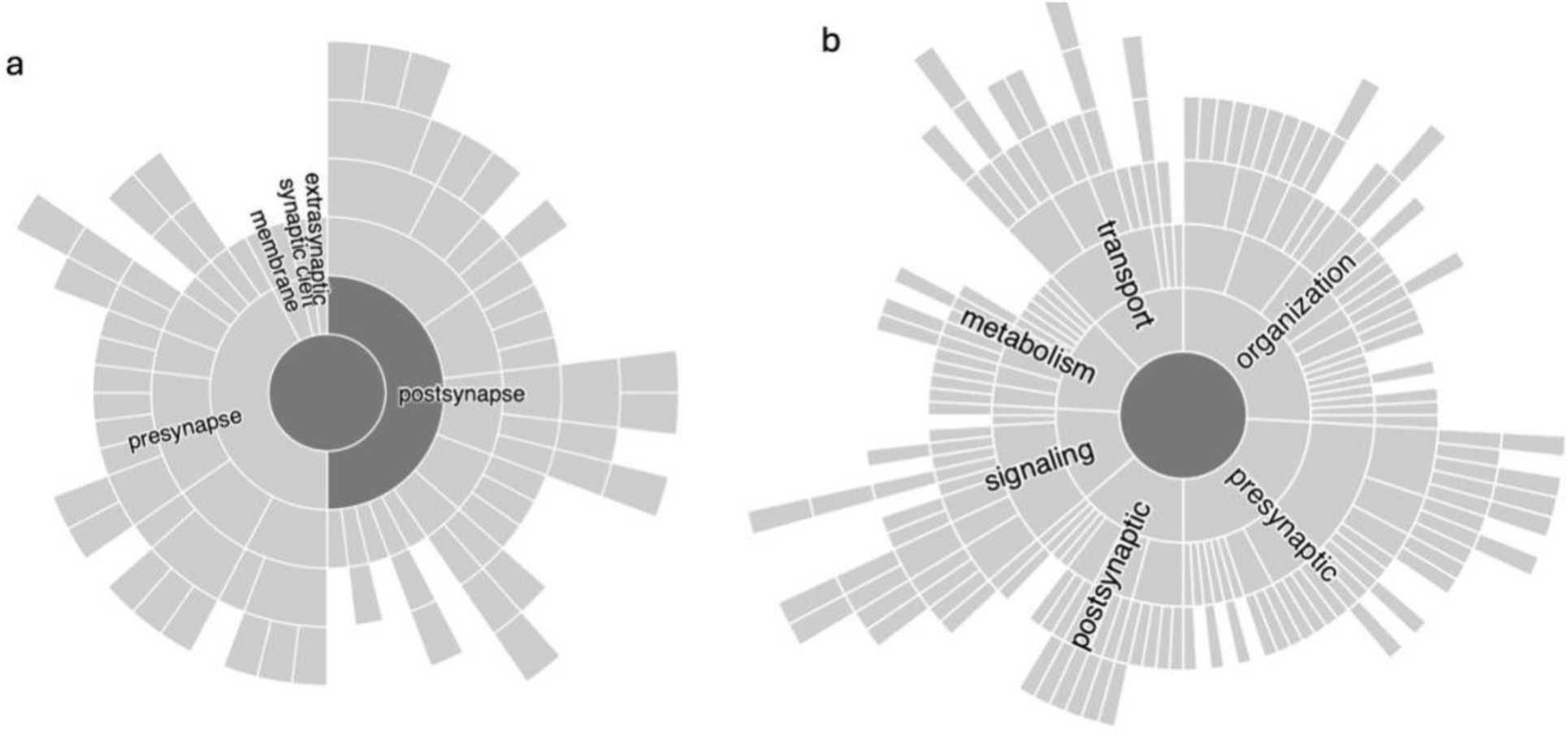
**a)** SynGo location enrichment analysis of genes which are differentially spliced in white matter across age. **b)** SynGo function enrichment analysis of genes which are differentially spliced in white matter across age.

**Figure S25.**
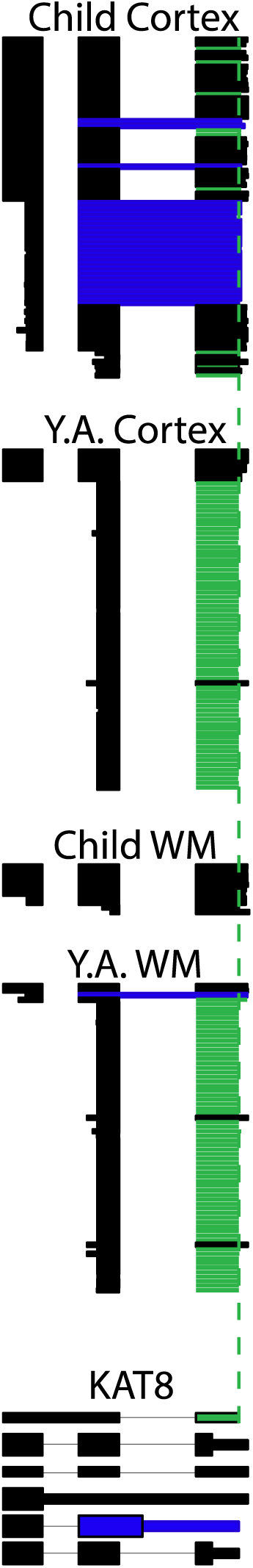
ScisorWiz plot of the gene KAT8. Top row: spliced, barcoded, and UMI corrected reads from the child age group which are located in the cortex. 2^nd^ Row: spliced, barcoded, and UMI corrected reads from the young adult age group which are located in the cortex. 3^rd^ Row: spliced, barcoded, and UMI corrected reads from the child age group which are located in white matter. Last Row: spliced, barcoded, and UMI corrected reads from the young adult age group which are located in white matter. Color indicates alternative last exon usage. Green: chr16:31,131,195-31,131,359; Blue: chr16:31,130,746-31,131,358. Black: All others. Green dotted line: aligns with start of exon chr16:31,131,195-31,131,359 (green label).

**Figure S26.**
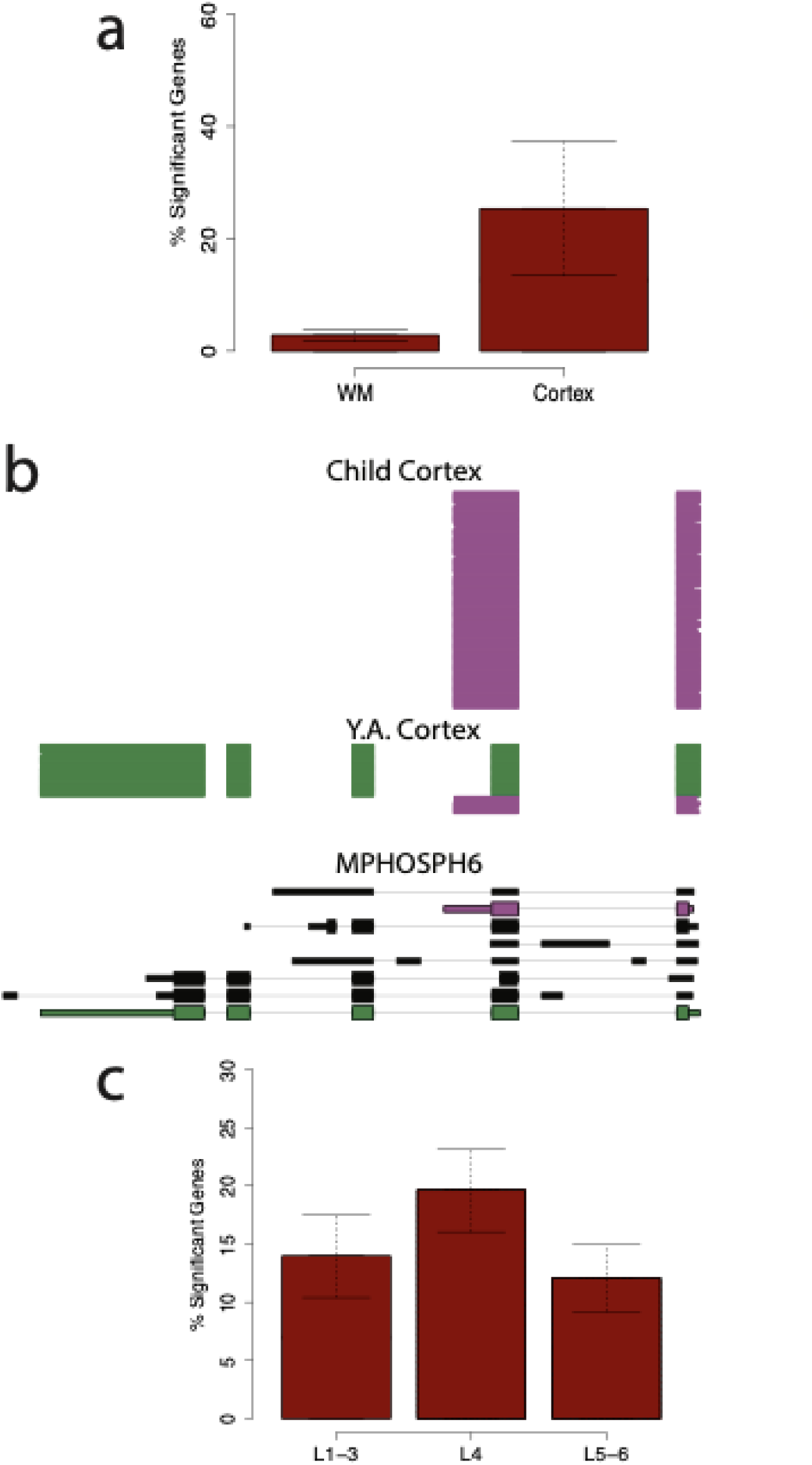
a) Percent significant genes which exhibit isoform changes across age, separated by brain region; WM: White Matter. b) ScisorWiz plot of the gene MPHOSPH6. Isoform1 is labeled in pink and Isoform2 is labeled in green. c) Percent significant genes which exhibit isoform changes separated by layer.

**Figure S27.**
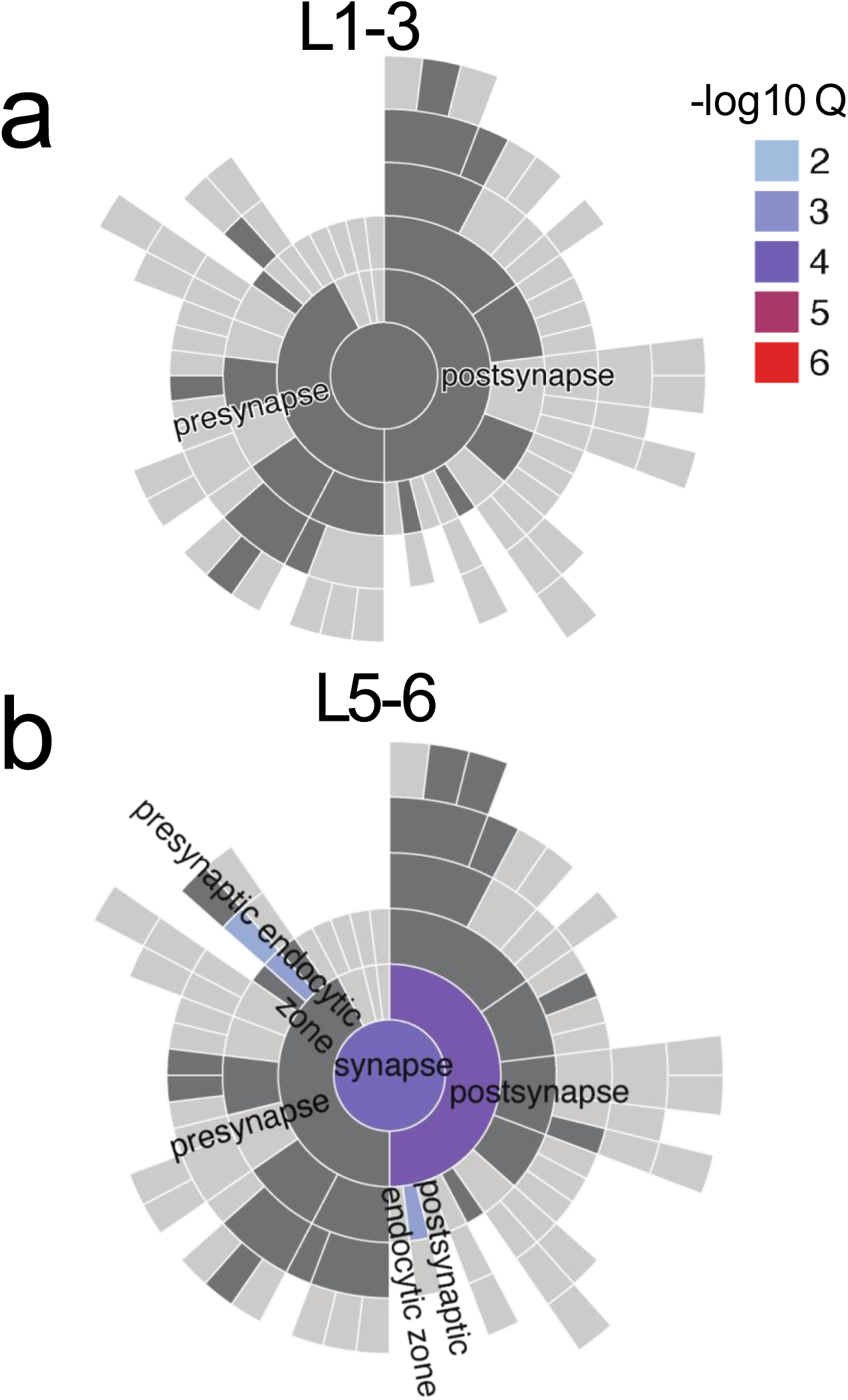
a) SynGo enrichment plots of genes with significant alternative exons in layers 1-3 across age. b) SynGo enrichment plots of genes with significant alternative exons in layers 5-6 across age.

**Figure S28.**
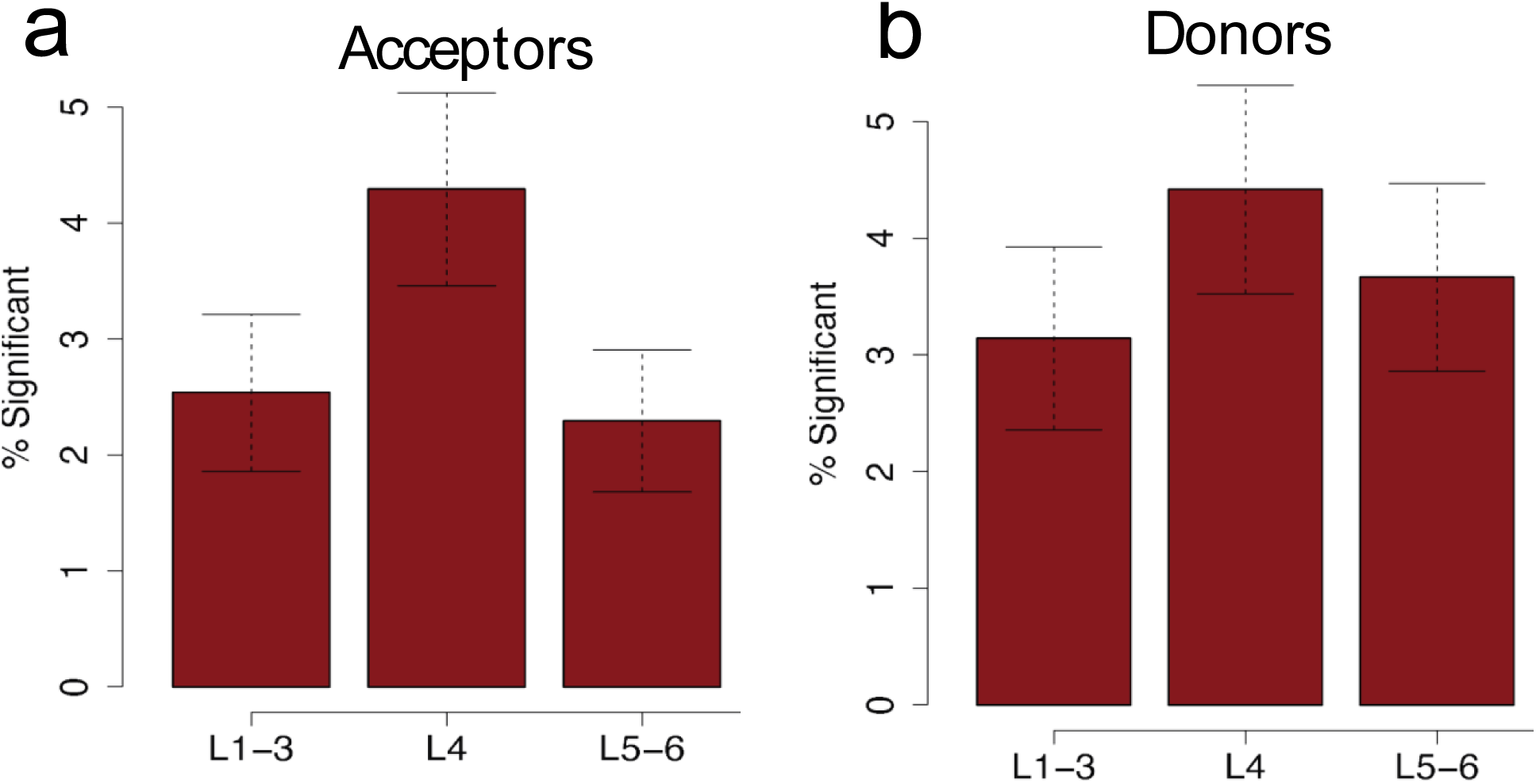
a) Percent significant alternative acceptor sites by layer. b) Percent significant alternative donor sites by layer.

**Figure S29.**
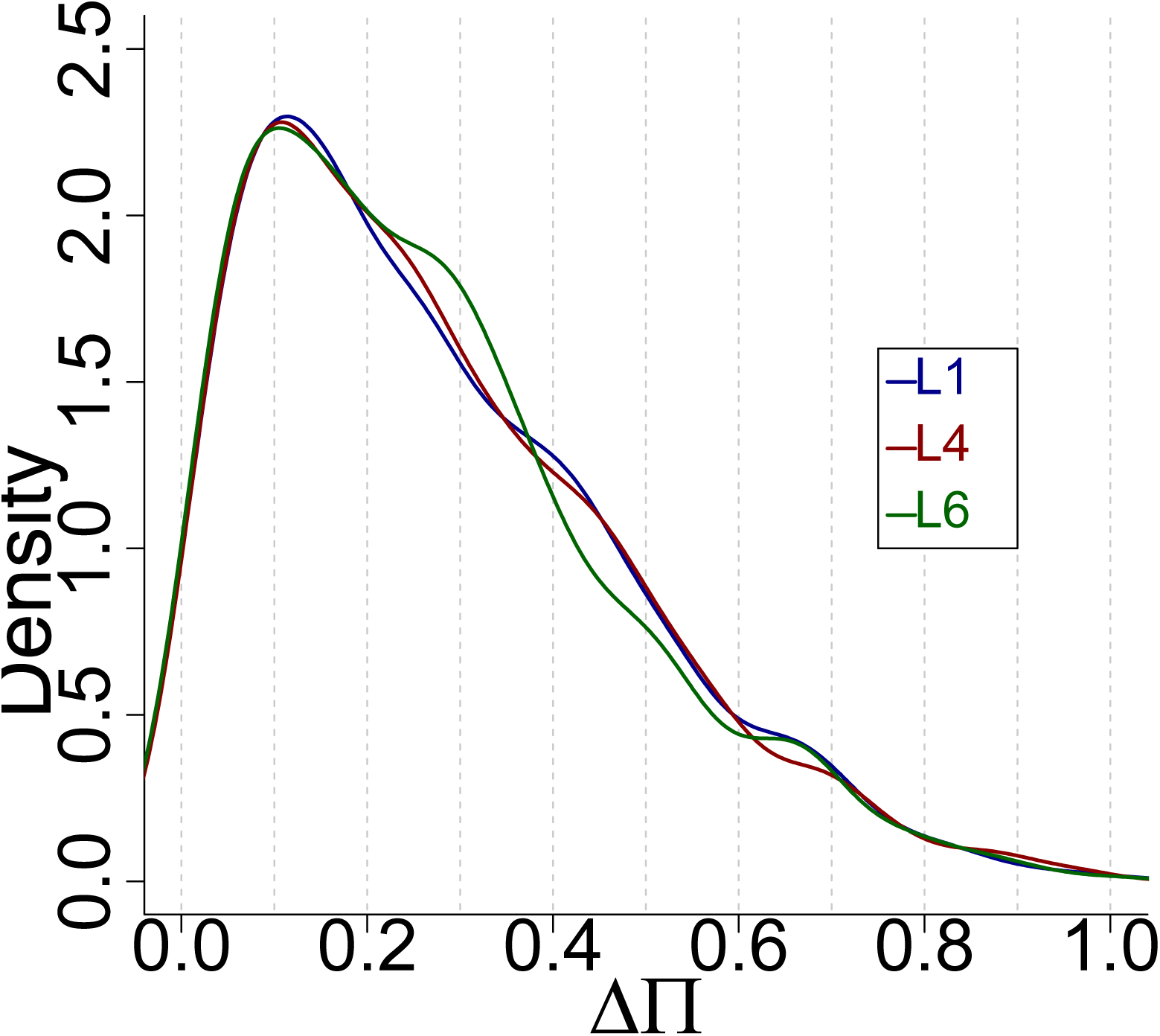
Density of |deltaPI| values identified for significant alternative Poly(A)-site usage. Lines are drawn by color to indicate layer; Blue = L1-3, Red=L4, Green=L5-6.

**Figure S30.**
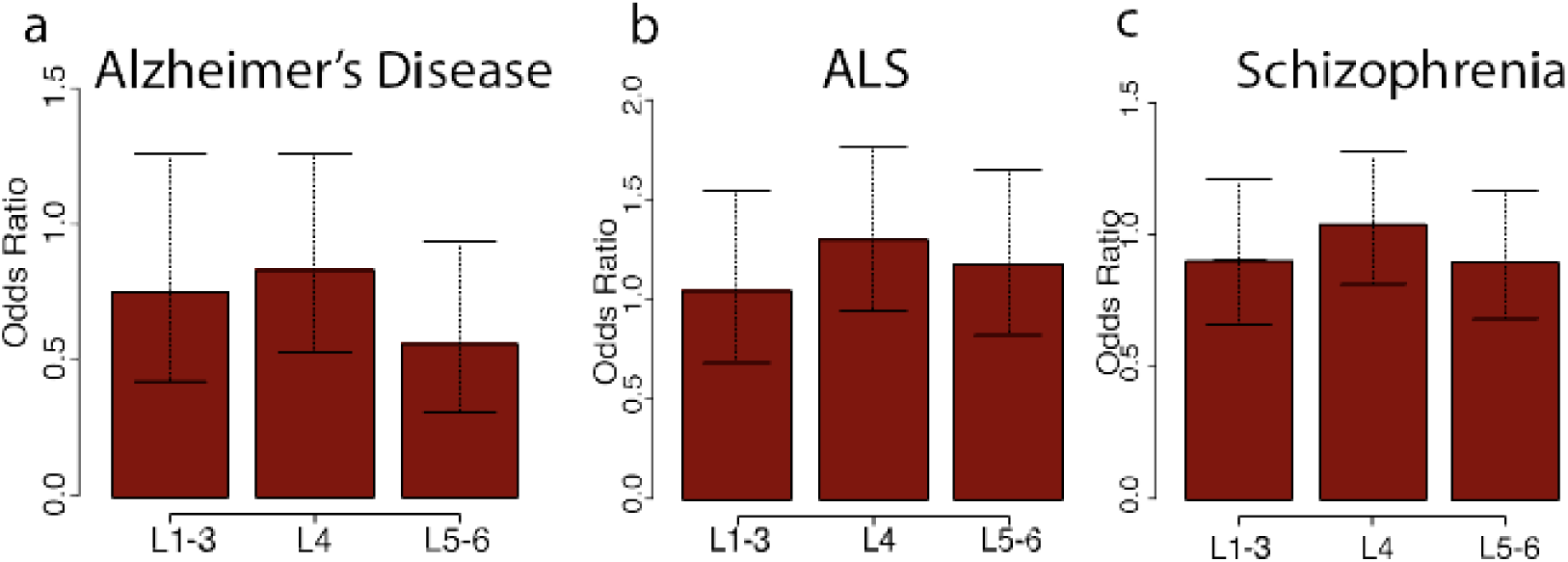
a) Odds ratio comparing significant group (|dPSI| > .5 and FDR < .05) v background group (|dPSI| < .1 and FDR > .05) of Alzheimer’s Diseases associated genes. b) Odds ratio comparing significant group v background group of ALS associated genes. c) Odds ratio comparing significant v background group of Schizophrenia associated genes.

**Figure S31.**
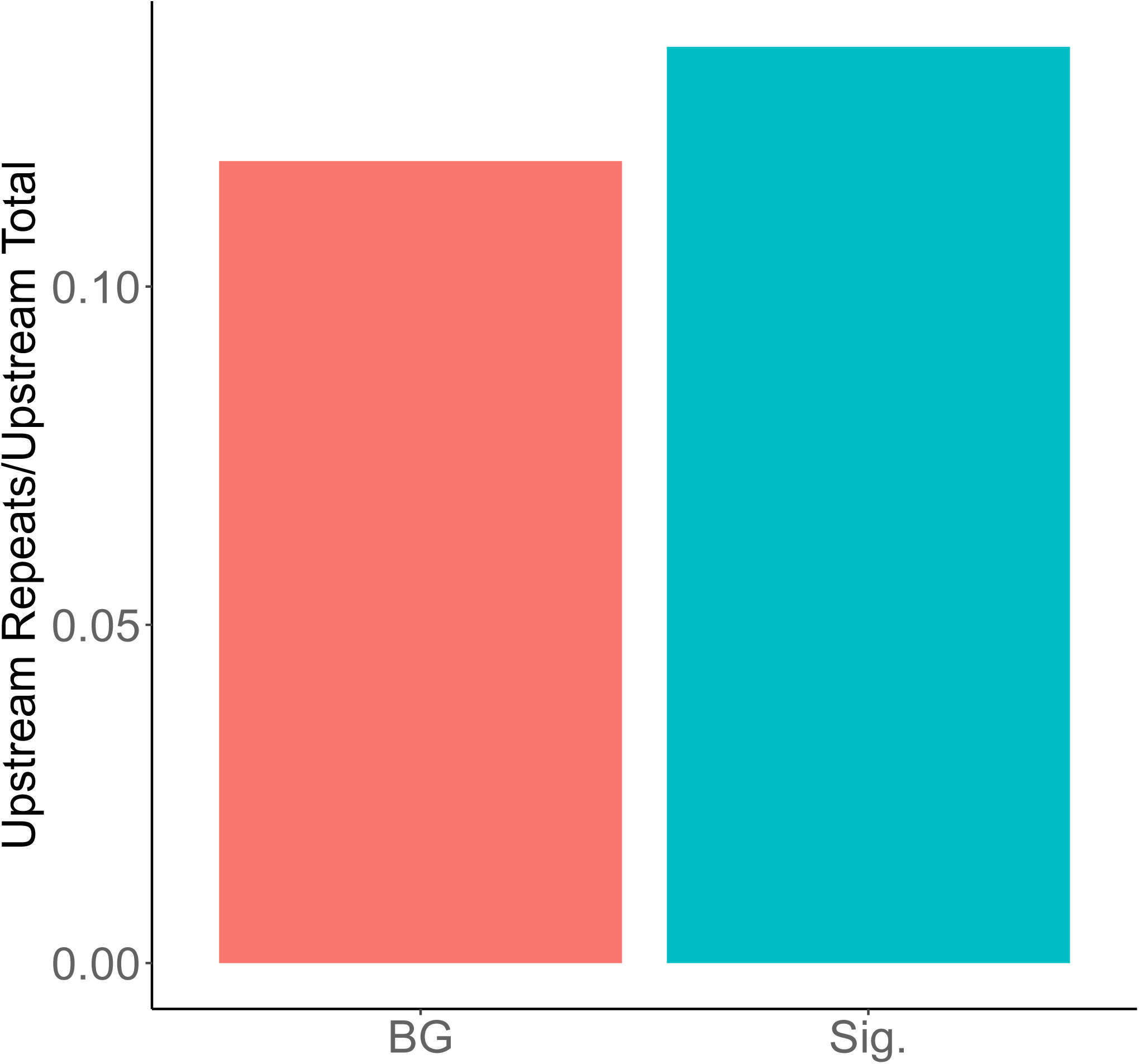
Ratio of sequences with repetitive elements found to total number of sequences per group. Sig: Sequences found upstream of exons with |dPSI| > 0.5 and FDR < .05. BG: Sequences found upstream of exons with |dPSI| < 0.05 and FDR > .05. 2-sided Fisher test, FDR = 0.334.

**Figure S32.**
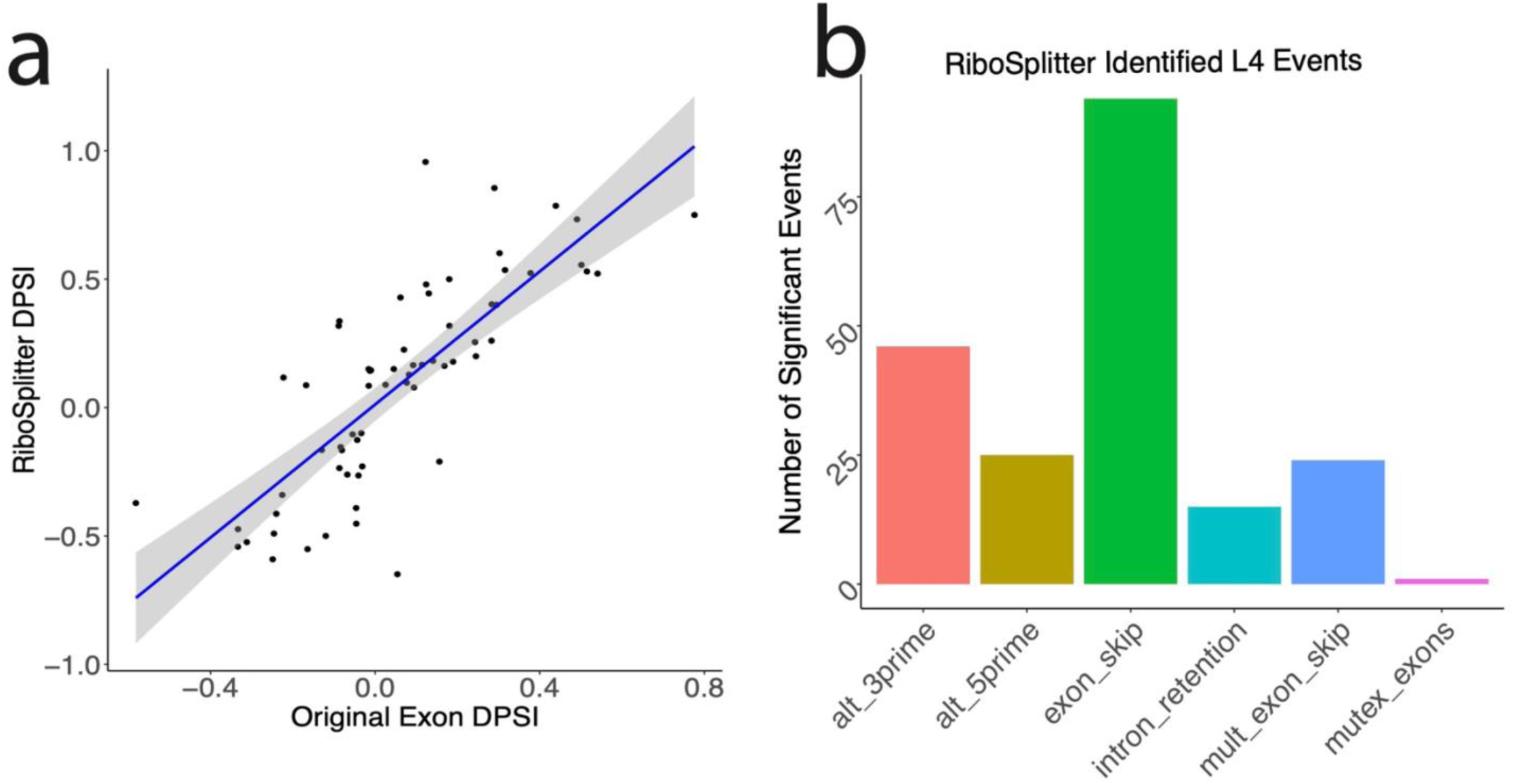
a) Correlation of the DPSIs of exons in L4 calculated as in Figure 6(a) compared to DPSIs calculated by RiboSplitter. Blue line represents the line of best fit. **b)** Number of significant events per category found by RiboSplitter.

## Notes

https://github.com/algbio/spl-IsoQuant.

